# Spatial and single-nucleus transcriptomic analysis of genetic and sporadic forms of Alzheimer’s Disease

**DOI:** 10.1101/2023.07.24.550282

**Authors:** Emily Miyoshi, Samuel Morabito, Caden M. Henningfield, Negin Rahimzadeh, Sepideh Kiani Shabestari, Sudeshna Das, Neethu Michael, Fairlie Reese, Zechuan Shi, Zhenkun Cao, Vanessa Scarfone, Miguel A. Arreola, Jackie Lu, Sierra Wright, Justine Silva, Kelsey Leavy, Ira T. Lott, Eric Doran, William H. Yong, Saba Shahin, Mari Perez-Rosendahl, Elizabeth Head, Kim N. Green, Vivek Swarup

## Abstract

The pathogenesis of Alzheimer’s disease (AD) depends on environmental and heritable factors, with remarkable differences evident between individuals at the molecular level. Here we present a transcriptomic survey of AD using spatial transcriptomics (ST) and single-nucleus RNA-seq in cortical samples from early-stage AD, late-stage AD, and AD in Down Syndrome (AD in DS) donors. Studying AD in DS provides an opportunity to enhance our understanding of the AD transcriptome, potentially bridging the gap between genetic mouse models and sporadic AD. Our analysis revealed spatial and cell-type specific changes in disease, with broad similarities in these changes between sAD and AD in DS. We performed additional ST experiments in a disease timecourse of 5xFAD and wildtype mice to facilitate cross-species comparisons. Finally, amyloid plaque and fibril imaging in the same tissue samples used for ST enabled us to directly link changes in gene expression with accumulation and spread of pathology.

## Introduction

Dating back to Santiago Ramón y Cajal, Paul Broca, and Korbinian Brodmann, it was revealed that the human brain is highly spatially organized at both macro- and microscopic levels, where both brain circuitry and function underlie this structural organization. Cajal’s famous drawings depicted the laminar organization of the brain and great morphological diversity across neurons. Advances in morphological, immunohistochemical, and electrophysiological methodology coupled with transcriptome sequencing have provided a modern framework for cataloging cortical cell types and unraveling their function at the circuit level ^1^. Single-cell (scRNA-seq) and single-nucleus RNA-sequencing (snRNA-seq) studies over the past few years have revealed that brain cell populations are highly heterogeneous at the molecular level ^2–11^. In Alzheimer’s disease (AD) brains, specific cell populations have been identified as under- or overrepresented relative to the cognitively healthy brain ^12–16^, and recent genomic studies have proposed an axis of selective vulnerability to resilience with regards to neuronal degeneration ^16, 17^. Holistically determining the functional consequences of disease-related changes of these molecularly distinct cell populations remains a challenge, and spatial information of these populations remains a critical piece to solving this puzzle.

Recently, several spatial profiling methods were developed with varying levels of resolution, throughput, and number of genes ^18–23^. Spatial transcriptomics (ST) relies on a grid of spatial spots with primers to uniquely barcode transcripts based on their spatial location and does not require pre-determined gene targets, thus allowing an unbiased assessment of gene expression changes. Further, this technique allows us to profile a whole coronally sectioned mouse hemisphere. However, the size of each spot is 55µm, resulting in transcriptomic profiles that may be from 1-10 cells. Therefore, in this study we generated both ST and snRNA-seq data to perform an integrated analysis of AD with cellular and spatial resolution. We paired our ST experiments with fluorescent amyloid imaging in the same tissue samples, allowing us to identify gene expression changes proximal to amyloid pathology. In principle, these datasets allow us to thoroughly characterize the cellular diversity and transcriptomic remodeling of the AD brain with underlying spatial context.

Here we examined the spatial and single-nucleus transcriptomes of clinical AD samples in both earlyand latestage pathology, as well as AD in Down syndrome (AD in DS). Although individuals with DS aged >65 years old have an 80% risk of dementia ^24^, only one previous study has profiled cell type-specific gene expression changes in DS brains ^25^. Further, despite shared features between AD in the general population and AD in DS ^26, 27^, there are currently no published single-cell or spatial transcriptomic studies examining both populations, and DS is a potentially advantageous group for clinical studies of AD. Focusing on AD in DS as a genetic form of AD, due to the vastly increased disease risk conferred by the triplication of *APP* on the 21st chromosome, provides opportunities for rigorous comparative analyses with the sporadic AD population and to further our understanding of the genetic underpinnings of sporadic AD. We uncovered spatial and cellular AD transcriptomic changes in both DS and the general population and additionally extended our analyses to a commonly used amyloid mouse model of AD, 5xFAD, by generating an additional spatial transcriptomic dataset. While mouse models are the most common non-human model systems used to probe AD biology, recent clinical trial failures of disease-modifying drugs in AD, which were successful in mouse models, have raised important questions about translatability of mouse findings to humans. While these failures can be attributed to many factors, including use of single-model systems for disease relevance and evolutionary distance between mouse and human, robust disease-specific translational targets can potentially be found using integrative approaches across human and mouse models. Our previous work in this area has led to the identification of regulators of neurodegenerative dementia and potential disease modifying drugs targeting those regulators ^28, 29^. Overall, our analysis identified spatially restricted and celltype-specific transcriptomic changes in different subtypes of AD, highlighting the value of employing a multi-faceted experimental and analytical approach using several models of investigation and multiple data modalities.

## Results

### Investigating the Alzheimer’s disease transcriptome with spatial and cellular resolution

We performed a cross-species study of spatially resolved gene expression changes in AD by generating ST data (10× Genomics Visium) from both postmortem human prefrontal cortical tissue (n=10 cognitively healthy controls, 9 early-stage AD, 10 late-stage AD, and 10 AD in DS; median 1,316 genes per spatial spot; n=115,251 ST spots; Table S1) and 5xFAD and wild type (WT) mouse brains (n=8-12 per group, 4 separate timepoints; median 2,438 genes per spatial spot; n=212,249 ST spots; Table S2) (Figures 1a, S1). We used BayesSpace ^31^, an unbiased clustering algorithm leveraging both transcriptomic and spatial information, to identify nine transcriptionally distinct clusters in our human dataset — 3 white matter (WM) clusters and 6 grey matter clusters encompassing the cortical layers — and 15 brain-region specific clusters in our mouse dataset (Figures 1b-c, S2,S3). We annotated these clusters based on the expression of known marker genes and their localization within the tissue sections and using unbiased cluster marker gene detection (Figure S4; Tables S3, S4). Due the heterogeneity inherent to postmortem human brain samples, we examined the distribution of the spatial spots by diagnosis for each cluster and found that most clusters are well represented across the diagnoses, except for WM3 in early-stage AD cases (Figure S1). Two mouse clusters lacked clear spatial localization, one being a mixture of erythrocytes and neurons based on marker gene expression, and the other a group of low quality spots based on UMIs, which we removed from our subsequent analyses (Figure S1).

**Figure 1.**
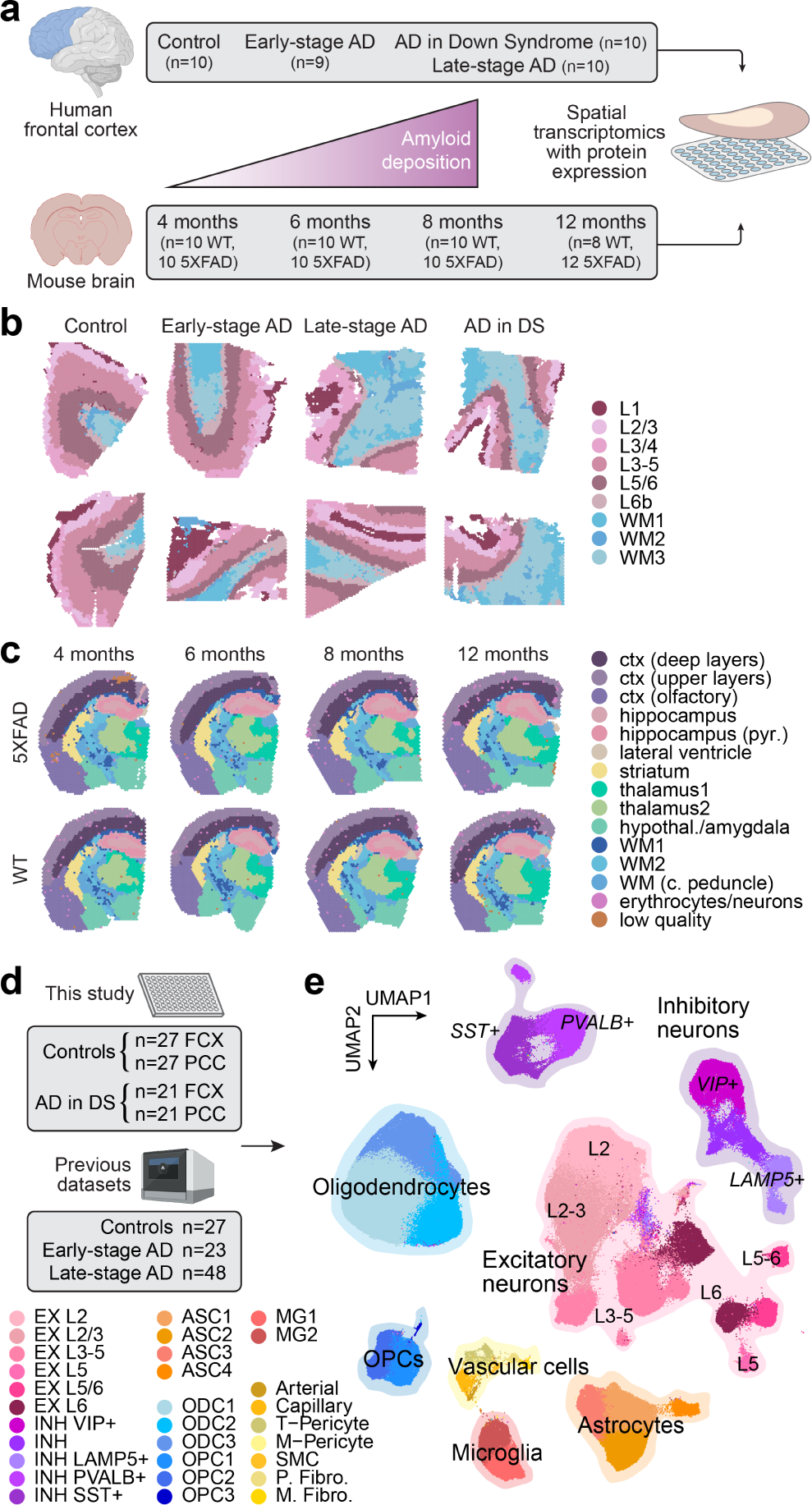
Spatial and single-nucleus transcriptomic analysis of genetic and sporadic forms of AD. **a**, We performed spatial transcriptomic experiments in the human frontal cortex and the mouse brain using 10X Genomics Visium. Human samples: n=10 cognitively normal controls; n=9 early-stage AD; n=10 late-stage AD; n=10 AD in DS. Mouse samples: n=10 WT and n=10 5xFAD aged 4 months; n=10 WT and n=10 5xFAD aged 6 months; n=10 WT and n=10 5xFAD aged 8 months; n=8 WT and n=12 5xFAD aged 12 months. **b**, Two representative human ST samples from each of the disease conditions, where each spot is colored by cortical annotations derived from unbiased spatial transcriptome clustering analysis. **c**, One representative mouse ST sample from WT and 5xFAD at each of the four time points, where each spot is colored by brain region annotations derived from unbiased spatial transcriptome clustering analysis. **d**, We performed single-nucleus RNA-seq (snRNA-seq) in the frontal cortex and posterior cingulate cortex from cognitively normal control donors (n=27 FCX, n=27 PCC) and AD in DS donors (n=21 FCX, n=21 PCC). We also included snRNA-seq data from three previous studies of the cortex in AD ^12–14^ (n=27 controls, n=23 early-stage AD, n=48 late-stage AD). **e**, Uniform manifold approximation and projection (UMAP) plot depicting a two-dimensional view of the cellular neighborhood graph of 585,042 single-nuclei transcriptome profiles. Each point in this plot represents one cell, colored by their cell-type annotations derived from Leiden clustering ^30^ analysis. Illustrations created with BioRender.com.

We also generated snRNA-seq data (Parse Biosciences, SPLiT-seq ^8^) from cognitively healthy controls (n=27 prefrontal cortex/FCX, 27 posterior cingulate cortex/PCC) and AD in DS (n=21 FCX, 21 PCC; 55 individuals total; Table S5). These brain regions were chosen for these experiments to cover a broad spectrum of the disease trajectory, as the PCC is affected in the early stages of disease while the FCX is affected in later stages ^32^. We used scANVI ^33^, a semi-supervised deep generative model with high performance in a recent comprehensive benchmarking study ^34^, to perform a reference-based integration of this dataset with snRNA-seq data from three previous studies of the AD cortex ^12–14^ (FCX; total n=27 cognitively healthy controls, 23 early-stage AD, and 48 late-stage AD) for a total of 585,042 nuclei after quality control (Figures 1d, S4; Methods). In this integrated dataset, our clustering analysis identified layer-specific excitatory neuron subpopulations (EX, n=229,041 cells); classical cortical inhibitory neuron subpopulations (INH, n=90,718); microglia (MG, n=20,197) and astrocyte (ASC, n=57,443) populations comprising a spectrum of homeostatic and activated cell states; multiple subtypes of oligodendrocyte precursor cells (OPC; n=23,053) and mature oligodendrocytes (ODC, n=153,182); and several rare vascular subpopulations as described in a previous transcriptomic study of AD ^35^ such as pericytes (PER, n=4,659), endothelial cells (END, n=3,637), fibroblasts (FBR, n=2,403), and smooth muscle cells (SMC, n=709, Figure 1e). Unbiased marker gene analysis of these cell clusters provided further context and support for their transcriptomic identity and underlying cell states (Figure S4, Table S6, Methods). Furthermore, we examined the expression of a post-mortem “ex vivo activation” signature, previously identified as an artifact of cellular stress during sample processing ^36^, and while we did find expression of select ex vivo activation genes like *HSPA1A*, *HSPA1B*, *FOS*, and *JUN*, the overall expression program score ^37^ was low (Figures S1, S4).

### Regional and cell type-specific gene expression changes in clinical Alzheimer’s disease

To identify disease-associated gene expression changes with cellular and spatial resolution, we performed differential expression analysis in each disease group (early-stage AD, late-stage AD, and AD in DS) with respect to controls for our human spatial and single-cell datasets (Figures S5, S6, S7; Tables S7-S12). We focused our analyses on the FCX, since we had both spatial and single-nucleus data for this region, and we found significant positive correlations between effect sizes for gene expression changes in the PCC and FCX across most cell clusters (Figures S8—S11). The extra copy of chr21 suggests overexpression of chr21 genes in our AD in DS samples, so we first examined differentially expressed genes (DEGs) by chromosome. We found that not all chr21 genes are upregulated in both our spatial and single-nucleus datasets, and upregulation was dependent on region or celltype (adjusted p-value < 0.05, Figure 2a-b). For example, *APP* expectedly is upregulated in AD in DS samples but interestingly is not significantly different from control samples in spatial cluster L3/4. Our findings are in line with Palmer & Liu et al.’s snRNA-seq study of DS ^25^. Moreover, we found many genes from other chromosomes that are also significantly changed in AD in DS (Figures 2a,b, S12, S13), indicating that triplication of chr21 initiates a cascade of transregulatory events that ripple throughout the entire genome. Our analysis identified substantial downregulation of genes in spatial cluster L3/4 across the diagnoses, which may reflect the preferential accumulation of pathology in cortical layer III ^38–40^.

**Figure 2.**
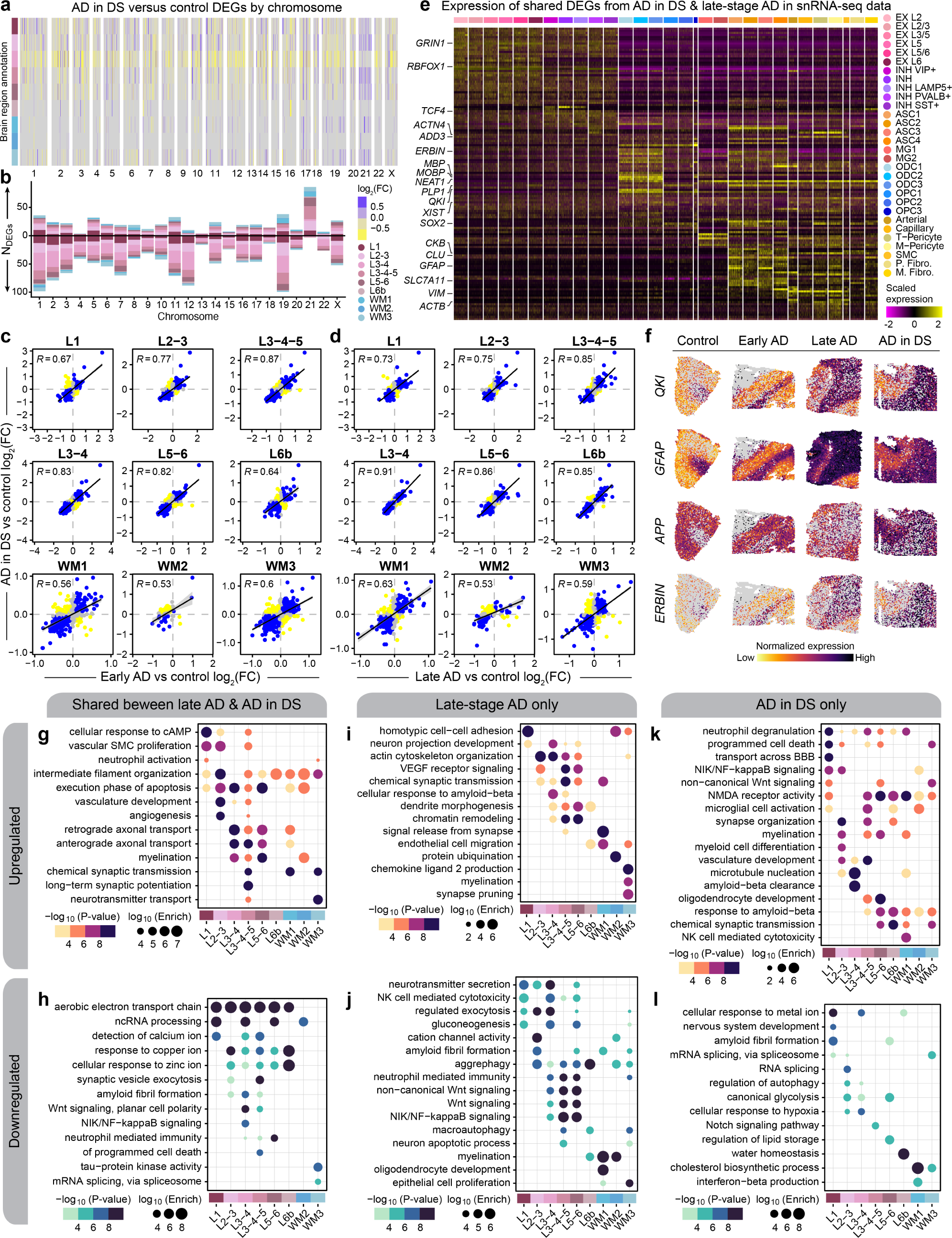
Shared and distinct gene expression signatures among subtypes of AD. **a**, Heatmap colored by effect size from the AD in DS versus control differential gene expression analysis, with genes stratified by chromosome and by spatial region. Statistically significant (FDR < 0.05) genes with an absolute average log2(fold-change) >= 0.25 in at least one region are shown. **b**, Stacked bar chart showing the number of AD in DS versus control DEGs in each spatial region stratified by chromosome. **c**, Comparison of differential expression effect sizes from early-stage AD versus control and AD in DS versus control. Genes that were statistically significant (adjusted p-value < 0.05) in either comparison were included in this analysis. Genes are colored blue if the direction is consistent, yellow if inconsistent, and grey if the absolute effect sizes were smaller than 0.05. Black line represents a linear regression with a 95% confidence interval shown in grey. Pearson correlation coefficients are shown in the upper left corner of each panel. **d**, Comparison of differential expression effect sizes from late-stage AD versus control and AD in DS versus control, layout as in panel (**c**). **e**, Heatmap showing the gene expression values in the snRNA-seq dataset of spatial DEGs shared between AD in DS and late-stage AD. **f**, Spatial feature plots of four selected DEGs (*QKI*, *GFAP*, *APP*, and *ERBIN*) in one representative sample from each disease group. **g-h**, Selected pathway enrichment results from DEGs that were upregulated (**g**) or downregulated (**h**) in both late-stage AD and AD in DS compared to controls. **i-j**, Selected pathway enrichment results from DEGs that were upregulated (**i**) or downregulated (**j**) in late-stage AD exclusively. **k-l**, Selected pathway enrichment results from DEGs that were upregulated (**k**) or downregulated (**l**) in AD in DS exclusively.

We next wanted to assess the similarities of disease-associated gene expression changes between diagnoses for each cluster. In our spatial data, although the majority of DEGs were unique to a diagnosis (early-stage AD, late-stage AD, or AD in DS), we still discovered many are conserved, including genes previously associated with AD, like *CST3*, *VIM*, *NEAT1*, and *CLU*. We also found significant and positive differential expression effect size correlations across spatial and single-nucleus clusters, except in smaller vascular clusters and OPC2 (Figures. 2c-d, S14, S15). Notably, these trends were stronger in grey matter clusters, compared to WM clusters, and this was matched in the single-nucleus data, where the correlations were also stronger in neuronal clusters relative to the glial clusters. We focused on the DEGs shared between late-stage AD and AD in DS, considering both conditions exhibit extensive amyloid and tau pathology. Disease-associated gene expression changes conserved between late-stage AD and AD in DS from the spatial data demonstrated strikingly region-specific patterns, and our snRNA-seq dataset revealed cell-type-specific expression of many spatial DEGs (Figure 2e-f). For example the gene *QKI*, previously identified as upregulated in AD ^41^, is upregulated only in upper cortical layers, although it is highly expressed in WM and oligodendrocytes. Gene ontology (GO) term enrichment of these shared DEGs revealed region-specific enrichment of AD-relevant biological pathways, such as upregulation of genes related to long-term potentiation in L3-5 and downregulation of those related to amyloid fibril formation in L3/4 and L3-5 (Figure 2g-h, Table S13). Genes associated with NIK/NF-κB signaling were downregulated in spatial cluster L3/4 and excitatory neuron clusters EX L2/3 and L3-5. However, we also found regional differences between AD in DS and late-stage AD; NIK/NF-κB signaling genes are upregulated in L1 and L2/3 exclusively in AD in DS samples (Figure 2i-l). Altogether, we identified conserved regional and cell-specific transcriptional changes between late-stage AD and AD in DS, supporting the utility of studying AD in DS to understand AD molecular changes in both DS and the general population.

In addition, we wanted to identify evolutionary-conserved AD transcriptional changes, as drug development largely relies on mouse models. However, mouse models of AD have been criticized for discrepancies with clinical AD. Considering previous literature shows baseline regional differences between human and mouse ^4, 42^, we hypothesized that there may be also regional differences in disease. In addition, we believe that focusing on the features shared between human and mouse will help forward *in vivo* research. We performed differential expression analysis on our mouse spatial transcriptomic dataset comparing 5xFAD versus WT within each spatial cluster for each timepoint (Figures S16, S17, S18, S19; Tables S14-17). This analysis revealed an increasing number of upregulated genes with age except in thalamic clusters, where the maximum was at the earliest timepoint (4 months, Figure S20). Upregulated genes at 4 months in the thalamus included disease-associated microglia (DAM) genes ^43^, like *Cst7*, *Tyrobp*, *Ctsd*, and *Trem2*, suggesting an early response to plaques localized to the thalamus, and with increasing age, we found upregulation of these genes across brain regions. We then examined the expression of DEGs identified in our human spatial dataset in a matched comparison with our mouse spatial dataset (Figure S21). We compared mouse cortical upper and deep layer clusters with the corresponding human cortical layer clusters (L1, L2/3, and L3/4; L3-5, L5/6, and L6b, respectively). White matter clusters were all individually compared with each other. We found significant and positive fold-change correlations in most cortical layer comparisons, except L1 and L6b. Like our comparison of AD in DS and late-stage AD, WM correlations were weaker, indicating that the 5xFAD model recapitulates some but not all clinical AD changes. Our analyses compiled a list of species-conserved, regional DEGs including some genes previously identified in disease-associated glia signatures ^43–45^.

### Multi-scale co-expression network analysis reveals spatial expression programs disrupted in AD

We performed a multi-scale co-expression network analysis in our ST dataset using hdWGCNA ^46^ (Methods). Co-expression networks enable elucidation of groups of genes (gene modules) that are highly co-expressed and are potentially co-regulated or involved in shared biological functions. We constructed co-expression networks and identified gene modules individually for each cortical layer cluster and white matter, yielding 166 modules from seven networks. Based on an approach previously employed in a brainwide network analysis in bulk RNA-seq ^47^, we defined a set of cortex-wide “meta-modules” by hierarchically clustering these 166 modules based on their similarity in gene expression patterns (module eigengenes, MEs) and their constituent gene sets (Figure 3a, Methods). This approach resulted in a unified set of 15 cortex-wide meta-modules and 166 fine-grain modules from individual spatial regions to define the spatial co-expression architecture of the cortex in AD (Figures 3a, S22, S23; Table S18). Using this compendium of gene-gene co-expression relationships across different spatial contexts, we interrogated systems-level differences in gene expression between disease groups and controls with differential module eigengene (DME) analysis (Figure 3a, Methods, Table S19).

**Figure 3.**
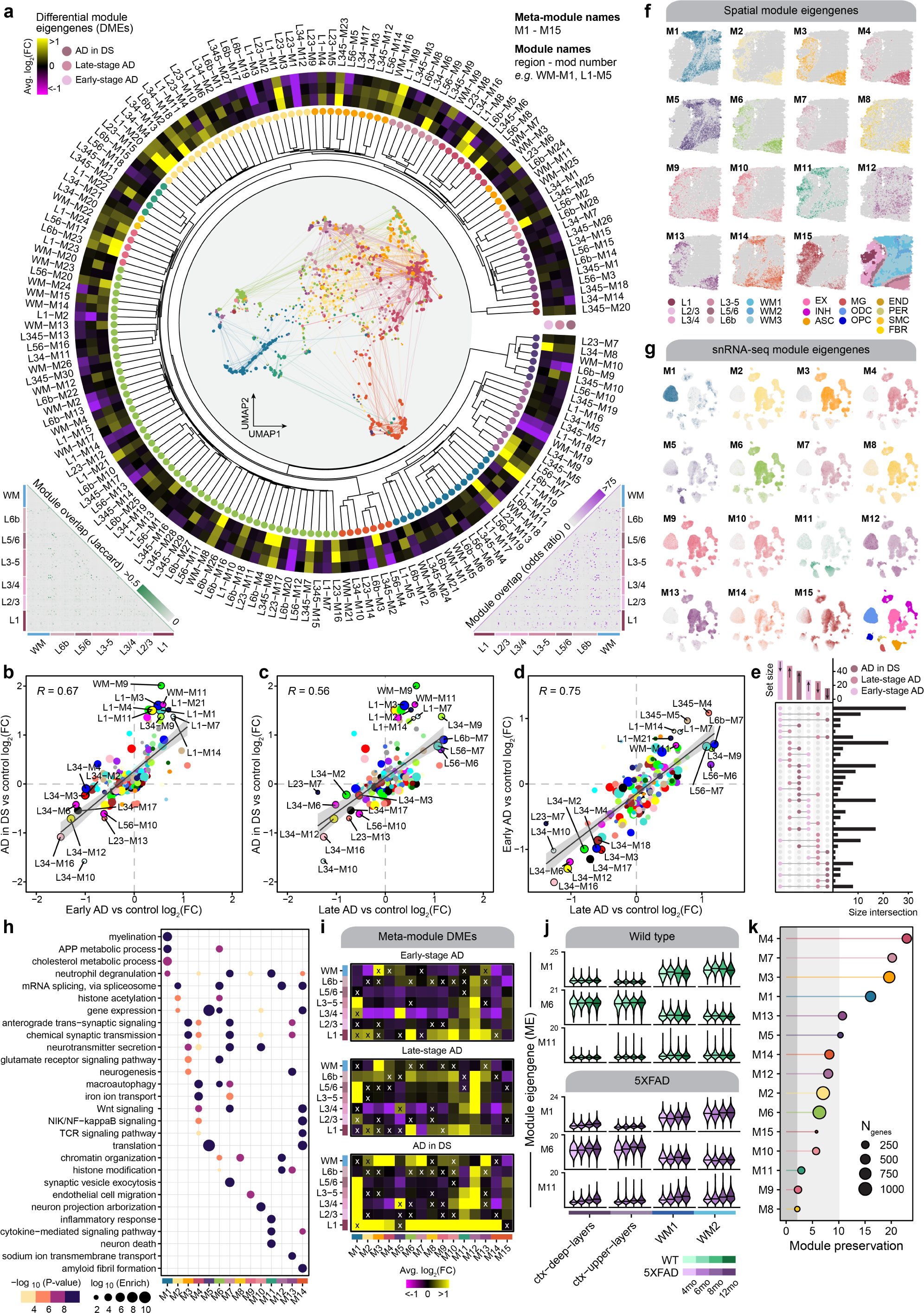
Multi-scale spatial co-expression network analysis reveals systems-level transcriptome alterations in AD. **a**, Dendrogram shows the of partitioning 166 gene co-expression modules into 15 brain-wide meta-modules. Network plot in the center represents a consensus co-expression network constructed from grey matter cortical clusters and white matter, where each dot represents a gene colored by meta-module assignment. Heatmap shows effect sizes from differential module eigengene (DME) testing in early-stage AD, late-stage AD, and AD in DS compared with controls. Triangular heatmaps show measures of distance between modules (bottom left: Jaccard index, bottom right: odds ratio). **b-d**, Comparison of effect sizes from differential module eigengenes testing in early-stage AD and AD in DS (**b**), late-stage AD and AD in DS (**c**), and late-stage AD and early-stage AD (**d**). Each dot represents a gene co-expression module, and selected modules are labeled. Black line represents a linear regression with a 95% confidence interval shown in grey. Pearson correlation coefficients are shown in the upper left corner of each panel. **e**, Upset plot showing the overlap between sets of differentially expressed modules in each disease group. **f**, Spatial expression profiles (module eigengenes, MEs) of the 15 meta-modules in one representative sample. Darker color corresponds to higher expression levels. A spatial plot with each spot colored by cluster assignment is shown in the bottom right corner as a visual aid. **g**, UMAP feature plots showing MEs of the 15 spatial meta-modules projected into the snRNA-seq dataset. Darker color corresponds to higher expression levels. The UMAP plot colored by major cell type is shown in the bottom right corner as a visual aid. **h**, Selected pathway enrichment results for each meta-module. **i**, Heatmaps showing the meta-module DME results for each cortical layer and white matter in the early-stage AD (top), late-stage AD (middle), and AD in DS (bottom) samples compared to controls. DME tests were performed similarly to those shown in panel (**a**). X symbol indicates that the test did not reach statistical significance. **j**, Violin plots showing MEs of selected meta-modules (M1, M6, and M11) in the 5xFAD mouse dataset, split by age group, with black line indicating the median ME. k, Lollipop plot showing module preservation analysis of the meta-modules projected into the 5xFAD mouse dataset. Z-summary preservation > 10: highly preserved; 10 > Z-summary preservation > 2: moderately preserved; 2 > Z-summary preservation: not preserved.

We compared the effect sizes between the different DME tests for each spatial module to assess the similarity in network-level changes between subtypes of AD (Figs. 3b-d). We broadly report similarities in the magnitude and direction of the DME results between the disease groups, indicating that the systems-level gene expression changes in AD in DS reflect those found in early-stage and late-stage AD, therefore complementing the similar comparative analysis of our DEGs. However, we also found that the largest set of unique differentially expressed modules were those downregulated in early-stage AD, suggesting that temporally transient systems-level expression changes greatly contribute to the molecular cascade underlying AD progression (Figure 3g). In addition, many modules demonstrate a larger effect size in early-stage AD, compared to late-stage and AD in DS.

We next sought to characterize the differences in cortexwide co-expression patterns by inspecting the 15 metamodules. We used the ST dataset and the snRNA-seq dataset to evaluate the spatial and cellular expression patterns of the meta-modules, noting that most of these expression signatures correspond to grey matter regions and neurons (Figs. 3h-i). These meta-modules were enriched for genes from biological processes pertaining to normal brain functions like myelination (M1) and chemical synaptic transmission (M3, M4, M7, M10, M13), as well as processes previously implicated in AD like glutamate signaling (M6), the inflammatory response (M11), and amyloid fibril formation (M14, Figure 3h, Table S20). We likewise performed DME analysis of the meta-modules to compare the disease groups to controls within each spatial cluster (Methods, Figure 3i). This analysis revealed that meta-module M6, which contains hub genes such as *APP*, *SCN2A*, and *CPE*, is consistently upregulated in cortical layer 1 (L1) in all three disease groups. While L1 is less densely populated with neurons than other cortical layers, module M6 is expressed primarily in neurons in our snRNA-seq data, together indicating key processes like APP metabolism, macroautophagy, and RNA splicing are altered in AD L1 neurons. On the other hand, meta-module M11 is expressed in glial cells and is upregulated across cortical upper layer clusters. M11 contains genes associated with the immune response and neuronal death, and its hub genes include members of the complement system (*C1QB*, *C3*), as well as genes previously identified in AD astrocytes ^12, 44^ (*SERPINA3*, *VIM*, *CD44*). In addition, we performed a crossspecies analysis of these co-expression networks identified in our human dataset by projecting the meta-modules to our 5xFAD ST dataset (Methods). This analysis enabled us to characterize the changes in module expression in different brain regions throughout aging in WT and 5xFAD mice (Figure S24). Expression levels of meta-modules M1, M6, and M11 were correlated with increasing age in 5xFAD mice but not in WT for several of these meta-modules, therefore representing changes associated with disease progression (Figure 3j). In particular, the gene expression signature of M11 appears to increase with pathology in both clinical AD and *in vivo* (Figure 3i-j). Furthermore, module preservation analysis ^48^ showed that the meta-modules identified in the human dataset were broadly preserved the mouse dataset (Figure 3k, Z-summary preservation > 2).

### Investigating sex differences within the AD in DS cohort

Understanding the effects of biological sex is particularly important in the context of AD as well as other neurological disorders, and previous AD studies have demonstrated sex differences in clinical manifestations, disease progression, risk factors, and gene expression ^13, 49–54^. However, there is limited knowledge as to how sex affects the AD transcriptome in the DS population. Here we performed differential gene expression analysis to investigate the sex differences in the transcriptome within our AD in DS cohort using our ST dataset (Figure 4a, Table S21, Methods). Given that our ST dataset had a greater number of female samples compared to male (n=7 female, n=3 male), we first sampled the ST spots in the female samples to match the total number in the male samples (Methods). We found gene expression differences across all chromosomes, and we discovered broad upregulation of genes in females compared to males across the spatial clusters (Figure 4b-d). Gene set overlap analysis demonstrated that many of the DEGs with the largest effect sizes were shared across multiple groups, such as the hemoglobin subunit beta gene *HBB* in females and *MOBP* in males (Figure 4e). Although *HBB* expression in female samples may be partly due to sample variability in blood contamination, hemoglobin is also expressed by neurons and glia. Further, a previous study found protein-protein interactions between Hbb and ATP synthase ^55^, and we also identified ATP synthase subunits (*ATP5PD* and *ATP5MPL*) as female DEGs. In addition, GO term enrichment analysis revealed that genes involved in inflammation, oxidative stress, and glucose metabolism are upregulated in females independent of brain region, whereas male DEGs are related to alternative splicing, chromatin organization, and cytoskeletal organization and transport (Figs. 4f-g, Table S22). Many DEGs were also exclusively found in only one or two spatial clusters (Figure 4e). Interestingly, amyloid-beta and amyloid fibril related processes were enriched in both the female and male specific DEGs. In males, this enrichment was restricted to L3-5, indicating regional specificity of sex differences in transcriptomic signatures related to amyloid processing (Figs. 4f-g). To complement our differential expression analysis comparing sex within AD in DS, we performed a similar analysis to identify differentially expressed co-expression modules for the 166 region-specific modules and the 15 cortex-wide meta-modules (Figs. 4h-i). This analysis highlighted many systems-level differences in the transcriptome between female and male samples; for instance meta-module M14 with genes related to Wnt and NIK/NF-κB signaling is upregulated in females across all spatial clusters.

**Figure 4.**
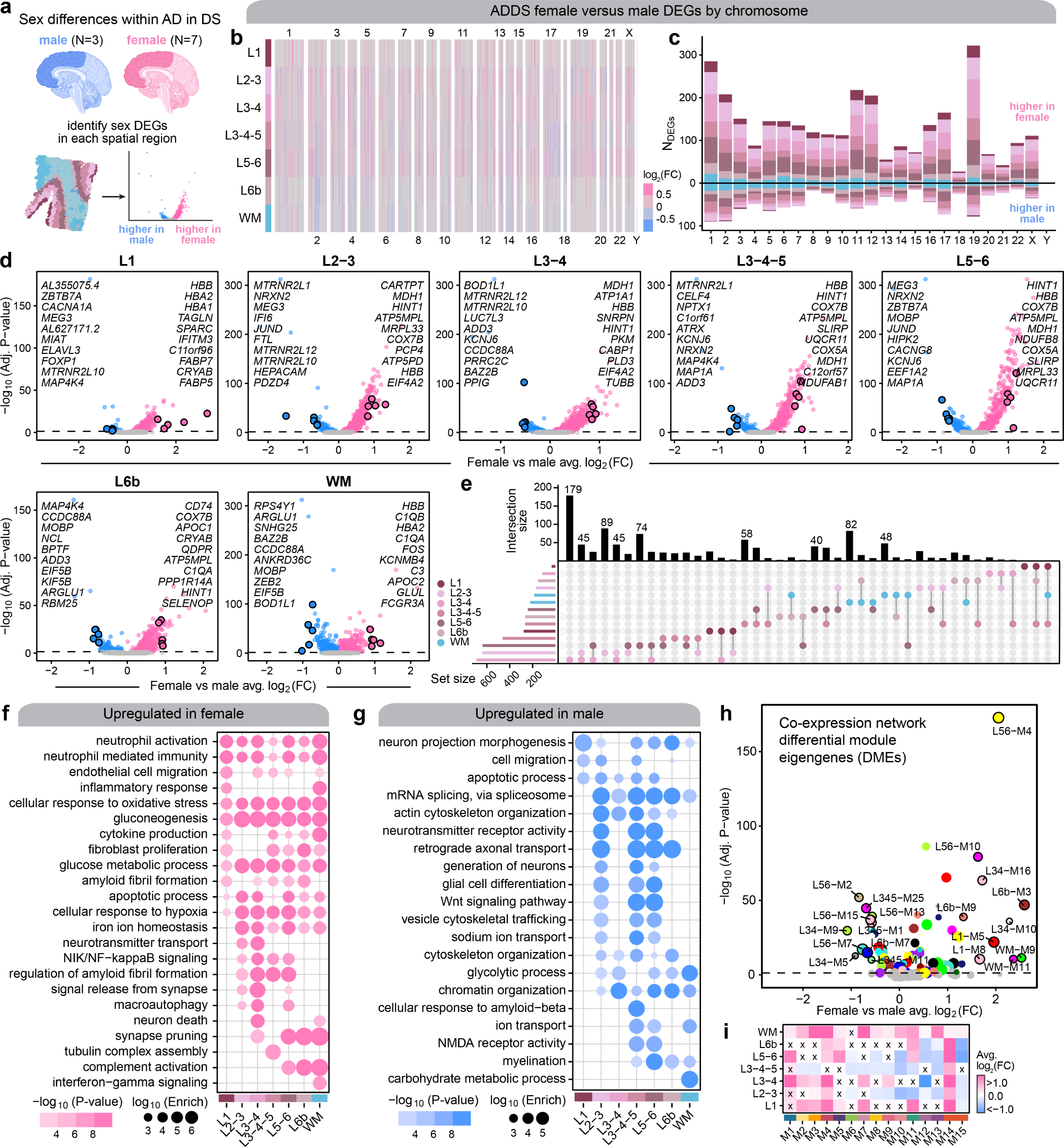
Sex differences within the AD in DS cohort. **a**, We performed differential expression analyses between female and male samples for each spatial cluster within the AD in DS cohort for our ST dataset. Since we had more female than male samples in the ST dataset, we subsampled the ST spots in the female samples to match the number from the male samples (Methods). **b**, Heatmap colored by effect size from the AD in DS female versus male differential gene expression analysis, with genes stratified by chromosome and by spatial region. Statistically significant (FDR < 0.05) genes with an absolute average log2(fold-change) >= 0.25 in at least one region are shown. **c**, Stacked bar chart showing the number of AD in DS female versus male DEGs in each spatial region stratified by chromosome. **d**, Volcano plots showing the effect size and significance level from the AD in DS female versus male differential expression tests in each of the spatial clusters. The top 10 and bottom 10 significant genes by effect size are labeled, excluding the set of genes that are commonly DE among all the spatial clusters. **e**, Upset plot showing the overlap between sets of DEGs in each spatial clusters. **f-g**, Selected pathway enrichment analysis from DEGs that were upregulated in female (**f**) or in male (**g**). **h**, Volcano plot showing the effect size and significance level from the AD in DS female versus male differential module eigengene (DME) analysis, with the top 10 and bottom 10 modules labeled. **i**, Heatmap showing the meta-module AD in DS female versus male DME results for each cortical layer and white matter. DME tests were performed similarly to those shown in panel (**h**). X symbol indicates that the test did not reach statistical significance.

### Spatial annotation of cell clusters to identify cell-cell signaling dysregulated in disease

While excitatory neuron subpopulations clustered and easily could be annotated by cortical layer, we were not able to stratify inhibitory neurons and glia by region with snRNA-seq alone. Therefore, we sought to leverage our spatial transcriptomic data to infer the region localization of our snRNA-seq clusters. We used CellTrek, an algorithm that integrates spatial and single-cell transcriptomic data to predict the spatial coordinates of a cell in a tissue section ^56^ For this analysis, we computed paired co-embeddings for each of the 39 human ST samples with the 48 snRNA-seq cortical samples generated in this study and used CellTrek to train a multivariate random forest model to predict the spatial coordinates of each cell using the shared embeddings (Methods, Figure 5a). We visualized these results in representative spatial samples by plotting each of the mapped snRNA-seq cells colored by their cell identity and split by major cell lineages (Figure 5a), as well as the expression of selected cortical layer marker genes side by side with the ST and mapped snRNA-seq datasets, confirming this approach assigns cell types to their known brain region (Figure 5b). Since this analysis was performed on each of the ST samples, each cell from the snRNA-seq data was mapped to multiple spatial contexts. Using this information, we computed a probability for each mapped cell to each of the spatial clusters. For example, this analysis uncovered that a subpopulation of *LAMP5*+ inhibitory neurons are WM interstitial neurons ^57^ (Figure 5c-d). Finally, for the snRNA-seq data we added a label for upper cortical layer, lower cortical layer, or white matter based on the spatial context that the cell was most frequently mapped to (Methods, Figure 5e). Furthermore, we inspected the predicted cell density distributions of specific cell subpopulations like subtypes of astrocytes and microglia in representative ST samples (Figure 5g). We also used these mappings to directly investigate the spatial and cellular contexts of systems-level transcriptomic signatures by visualizing the meta-module eigengenes in the predicted ST coordinates split by the major cell types (Figure 5f).

**Figure 5.**
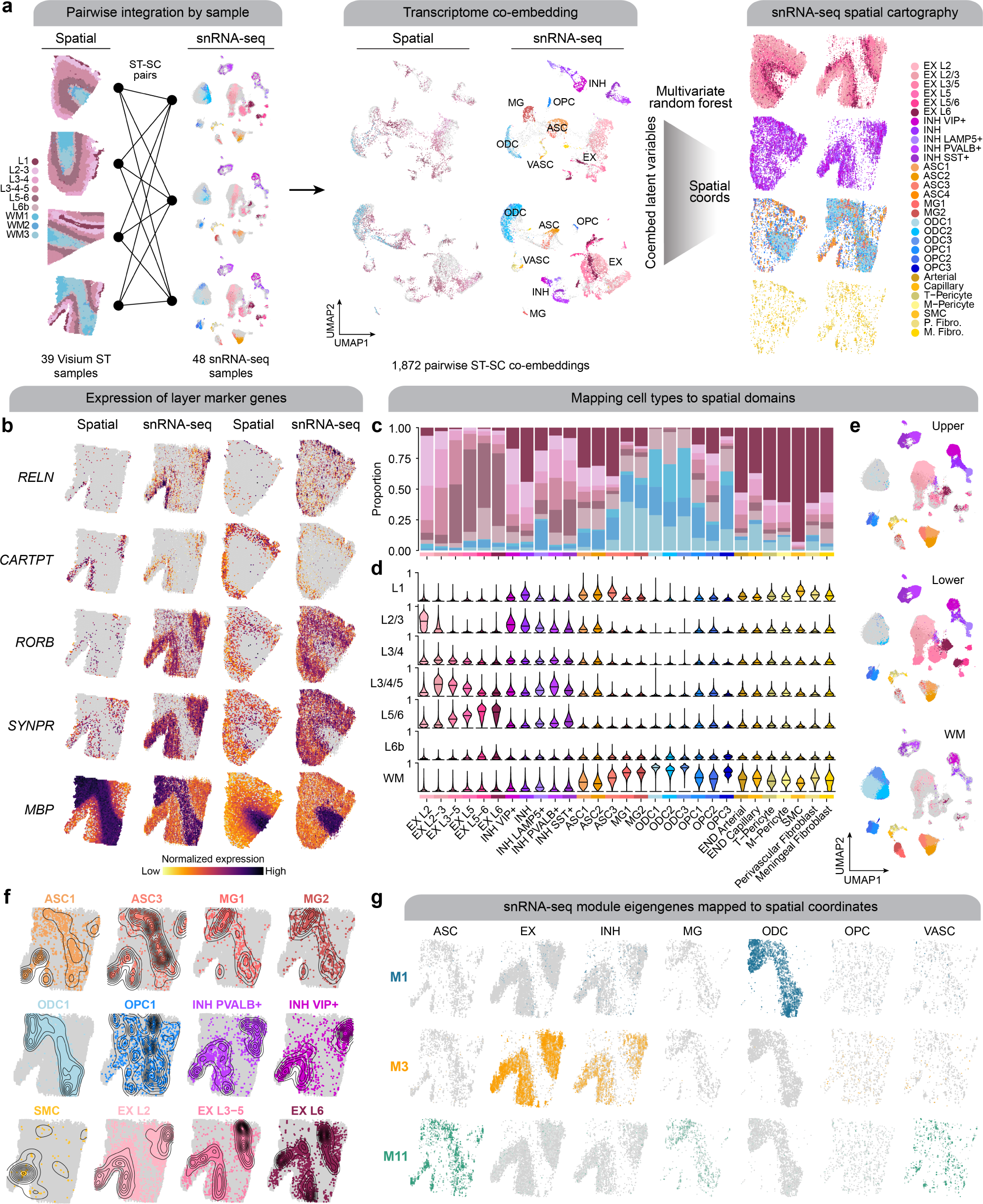
Systematic integration of spatial and single-nucleus expression profiles. **a**, We performed a systematic pairwise integration of biological samples profiled with spatial and single-nucleus transcriptomics (left). For all possible pairs of ST + snRNA-seq samples, we constructed a transcriptomic co-embedding (middle), and then we used a multivariate random forest model (CellTrek ^56^) to predict the spatial coordinates of each snRNA-seq cell in the given spatial context. The snRNA-seq dataset is shown on the right projected into two different spatial contexts (left: control sample; right: AD in DS sample), split by major cell lineages and colored by cell annotations. **b**, Spatial feature plots of selected layer-specific marker genes, shown side-by-side in the ST dataset and the snRNA-seq dataset projected into the spatial context for one AD in DS sample (left) and one control sample (right). **c**, Stacked bar plot showing the proportion of nuclei from each snRNA-seq cluster mapped to the spatial domains defined by the ST clustering. **d**, Violin plots showing the distribution of spatial domain mapping probabilities for nuclei from each of the snRNA-seq clusters. **e**, UMAP plot of the snRNA-seq dataset split by predicted spatial partitions into the upper cortical layers, lower cortical layers, or the white matter. **f**, Spatial density plot showing the snRNA-seq dataset in predicted spatial coordinates, highlighting selected cell populations. **g**, Spatial feature plots showing selected module eigengenes in the snRNA-seq dataset in predicted spatial coordinates.

With these spatial annotations of our snRNA-seq clusters, we next wanted to assess how cell-cell communication may be dysregulated in AD using CellChat ^58^. CellChat identifies signaling pathways between cells based on the expression of ligand and receptor pairs and performs statistical testing to compare the signaling patterns between different experimental groups. The additional spatial information allows us to distinguish these signaling changes as shortor longrange cellular communication, which is otherwise lost with snRNA-seq data alone. This analysis broadly uncovered differences in the strength of signaling interactions across different cell clusters between AD in DS and controls (Figure 6a). Across all the clusters, we found a reduced number of interactions in AD in DS compared to control, but an increased strength of the interactions (Figure 6b-c). We next performed an unsupervised machine-learning based analysis to inspect the functional changes in the cell-cell communication landscape of specific signaling pathways (Figure 6d) and to compute the relative information flow between AD in DS and controls (Figure 6e). We identified several signaling pathways up- or downregulated in disease (Figure 6f-h, Table S23) and highlight the NECTIN, ANGPTL, and CD99 signaling pathways. We found NECTIN signaling is downregulated in AD in DS. Nectins are involved in the formation and maintenance of synapses ^59–61^, and we discovered an absence of inhibitory neuronal NECTIN signaling, as well as a loss in several excitatory neuron subpopulations (Figures 6f-i). Though it is difficult to discern if these loss in NECTIN signaling is attributable to neurodegeneration or DS developmental changes, *NECTIN2* is a known AD risk gene ^62, 63^, indicating the importance of further studying the role of neuronal NECTIN signaling in AD. On the other hand, ANGPTL signaling is upregulated in AD in DS. In control samples, ANGPTL signaling features astrocyte clusters in cortical lower layers and the white matter (ASC1 and ASC3, respectively) communicating with a variety of cell types, including neurons, pericytes, and oligodendrocyte progenitor cells (OPCs), with the ligand *ANGPTL4* (Figures 6j-m). However in AD in DS, additional astrocytes, like ASC1 and ASC3 in the cortical upper layers, express *ANGPTL4*. Increased *ANGPTL4* expression has been previously observed in astrocytes from AD patients with vascular changes ^64^, and *ANGPTL4* is a hub gene of the metamodule M11, which we identified as upregulated with disease in the cortical upper layers. Interestingly, we also note a loss of astrocyte-inhibitory neuron ANGPTL communication with disease. Conversely, *CD99* is also a hub gene of M11, but CD99 signaling is downregulated in AD in DS (Figure S28). We discovered pericyte and astrocyte (ASC3) communication with excitatory and inhibitory neuronal populations is absent, suggesting that disrupted CD99-CD99L2 signaling may contribute to the neurovascular changes seen in AD. Notably, despite the overall downregulation of CD99 signaling in AD in DS, we also found CD99-PILRA signaling between cortical lower layer pericytes and microglia exclusively in AD in DS. PILRA, a myeloid inhibitory signaling receptor, has been associated with AD genetic risk ^65–68^; our findings implicate pericytes in the modulation of microglial function in AD via PILRA activity.

**Figure 6.**
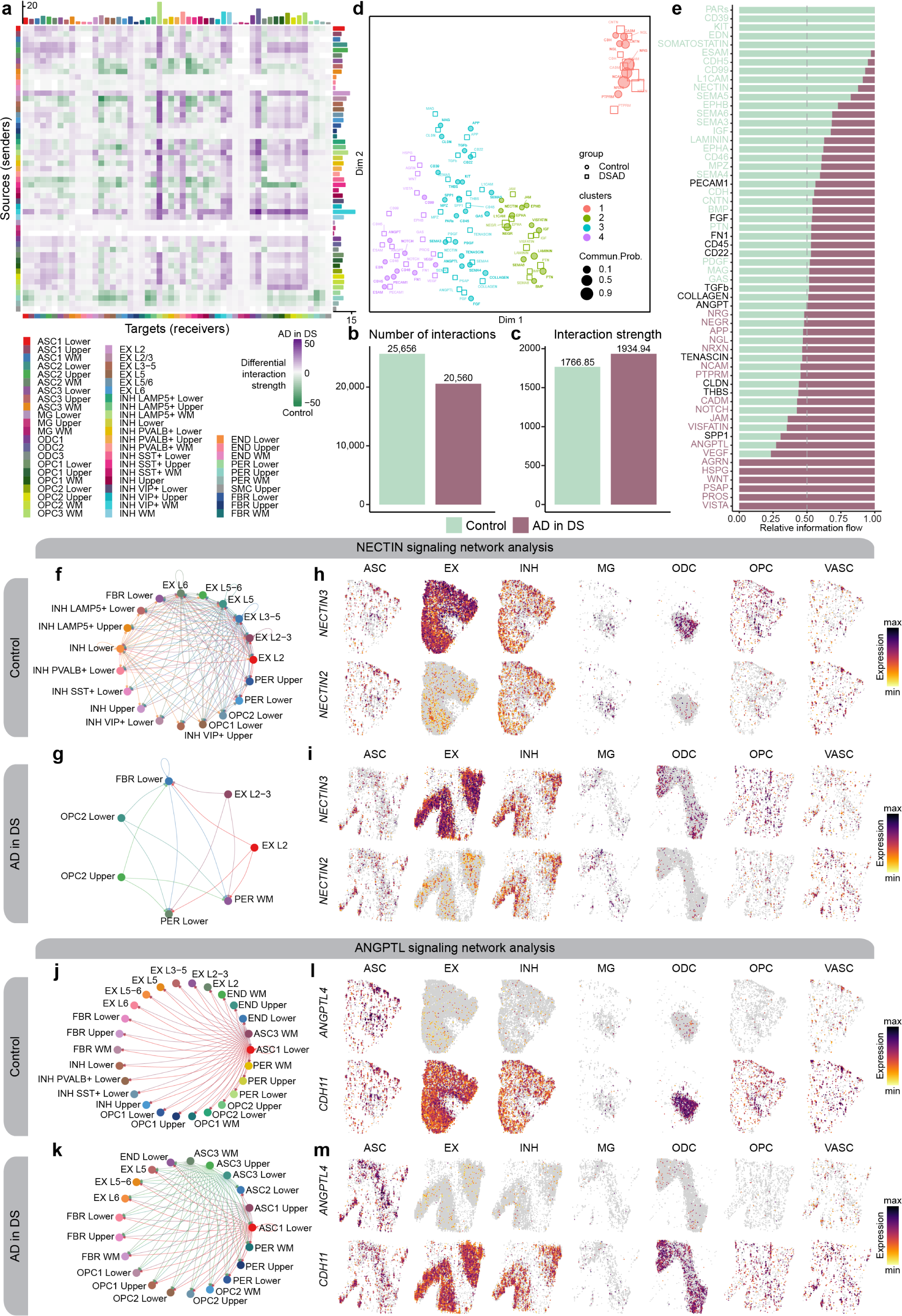
Altered cell-cell communication signaling networks in AD in DS. **a**, Heatmap showing the differential cell-cell communication (CCC) interaction strength between AD in DS and control. Each cell represents a snRNA-seq cell population, where rows correspond to signaling sources and columns correspond to signaling targets. Bar plots on the top and right show the sum of the incoming and outgoing signaling respectively. **b-c**, Bar plots showing the total number of CCC interactions (**b**) and interaction strength (**c**) for control and AD in DS. **d**, Joint dimensionality reduction and clustering of signaling pathways inferred from AD in DS and control data based on their functional similarity. Each point represents a signaling pathway. **e**, Bar plots showing signaling pathways with significant differences between AD in DS and controls, ranked based on their information flow (sum of communication probability among all pairs of cell populations in the network). **f-g**, Network plot showing the CCC signaling strength between different cell populations in AD in DS (**f**) and controls (**g**) for the NECTIN signaling pathway. **h-i**, Spatial feature plots of the snRNA-seq in predicted spatial coordinates for one control sample (**h**) and one AD in DS sample (**i**) for one ligand and one receptor in the NECTIN pathway. j-k, Network plot as in panels (**f-g**) in AD in DS (**j**) and controls (**k**) for the ANGPTL signaling pathway. **l-m**, Spatial feature plots as in panels (**h-i**) for one control sample (**l**) and one AD in DS sample (**m**) for the ANGPTL pathway.

### Imaging mass cytometry reveals protein-level expression changes in AD

To complement our transcriptomic analysis, we leveraged imaging mass cytometry (IMC) for multiplexed imaging of a panel of 23 proteins, allowing us to investigate the spatial expression patterns of these proteins (Figure 7a, Methods). Briefly, this technique uses a panel of metal-conjugated antibodies for multiplexed labeling, followed by fast UV-laser tissue ablation and cytometry by time of flight (CyTOF) to yield quantitative single-cell spatial proteomic data, and this approach has been previously applied in postmortem human brain tissue ^69^. This approach circumvents the constraints of conventional immunohistochemistry (IHC), which only allows for analysis of a handful of markers, restricting the understanding of cellular phenotypes, and avoids the autofluorescence issues inherent to aged human brain tissue. By devising and validating a highly multiplexed IMC approach, we were able to profile brain cell types including neurons, astrocytes, and microglia. Following tissue laser ablation, spatial expression data was acquired at approximately 1 um resolution via mass spectrometry analysis of metallic labels.

**Figure 7.**
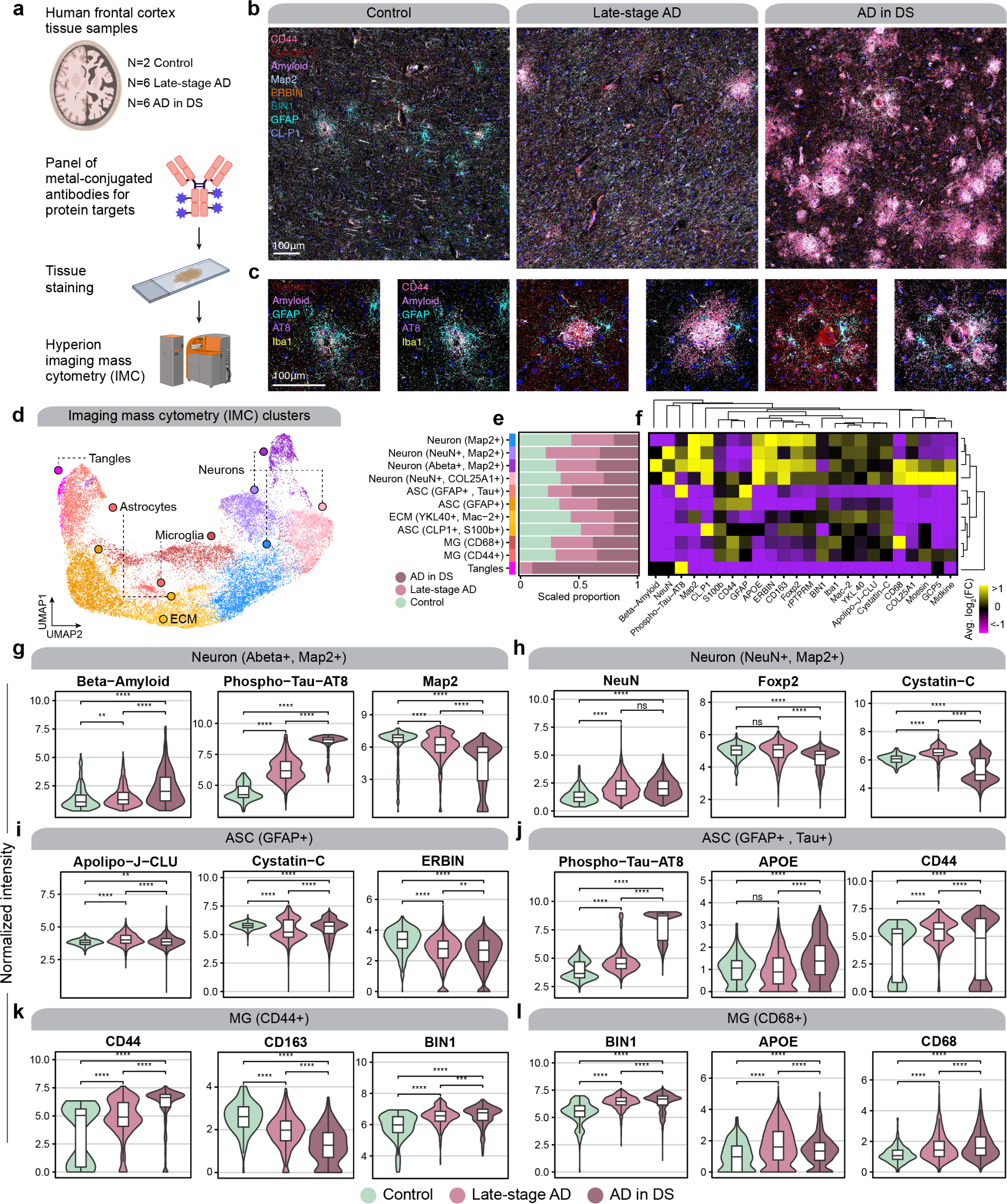
Imaging mass cytometry reveals spatial proteomic changes in AD. **a**, We performed imaging mass cytometry (IMC) in postmortem human cortical tissue (n=2 control, n=6 late-stage AD, n=6 AD in DS) using the Standard BioTools Hyperion Imaging System. **b**, Representative IMC images from control, late-stage AD, and AD in DS samples with select targets from the panel. **c**, Images as in panel (**b**) at higher magnification and focused around amyloid plaques. **d**, UMAP plot showing the unbiased clustering of segmented nuclei from the IMC dataset based on their protein intensity values. Each dot represents a segmented nucleus, colored by cluster assignment. **e**, Stacked bar plots showing the proportion of segmented nuclei assigned to each cluster stratified by disease groups. **f**, Heatmap showing the relative protein intensity of each protein in each IMC cluster. Dendrograms depict hierarchical clustering results based on these relative intensities. **g-l**, Violin plots showing the distribution of protein intensities for selected proteins in specified IMC clusters, stratified by disease groups. Wilcox test comparison results are overlaid on each plot. Not significant (ns), p > 0.05; * p <= 0.05; ** p <= 0.01; *** p <= 0.001, **** p<= 0.0001.

We used IMC to analyze the spatial protein expression patterns in gray matter and white matter cortical tissue samples from the same cohort used for our ST data (post-QC sample numbers: cognitively normal controls, n= 2; latestage AD, n=6; and AD in DS, n=6). The constructed images remarkably mirror images obtained from classical microscopy but with greater channel capacity (Figs. 7b-c). We applied a supervised machine learning approach to generate cell segmentation masks, allowing us to quantify protein intensities for individual cells, yielding 32,759 cellular profiles after quality controls (Methods). Using this single-cell protein data with spatial information, we performed an unbiased clustering analysis and revealed 11 distinct populations (Methods, Figure 7d). Our antibody panel did not include cell type marker proteins for all brain cell types due to limited antibody availability and compatibility, but we were able to recover proteomic profiles for astrocytes (GFAP+, S100b+), neurons (Map2+, NeuN+), and microglia (Iba1+), as well as populations that were enriched in extracellular matrix (ECM) proteins or phospho-tau (Figs. 7e-f).

We next used this dataset to perform statistical comparisons of protein abundances between disease groups in the different IMC clusters (Figs. 7g-l, Table S24, Methods). We found that the abundances of amyloid beta and phosphotau were significantly increased in neurons with disease, as expected, but microtubule-associated protein 2 (MAP2) decreased, indicative of neurodegeneration (Figure 7g). We found that abundance of Cystatin C (*CST3*), which is known to colocalize with amyloid beta, was significantly changed in microglia and astrocytes from late-stage AD and AD in DS (Figure 7i). In our spatial transcriptomic data, *CST3* was upregulated in both groups in spatial clusters L3-5 and L5/6; however, we did not find the same congruence in our snRNA-seq data. *CST3* was upregulated in only ASC1 for AD in DS and MG2 for late-stage AD. We additionally discovered CD44 is upregulated in astrocytes and microglia from both diagnoses (Figs. 7j-k). Similar to *CST3*, we found that *CD44* expression in the spatial transcriptomic data was more concordant than that in the snRNA-seq data. We expected CD44 upregulation in astrocytes, but we did not detect microglial upregulation of *CD44* at the single-nucleus transcriptome. These discrepancies may be due to the biological discrepancy between gene and protein expression or other technological and algorithmic limitations. However, we found a significant upregulation of clusterin (Apolipoprotein J, *CLU*) in both microglia and astrocyte subpopulations with increased CD44 protein expression in both late-stage AD and AD in DS. *CD44* and *CLU* are hub genes of meta-module M11, which we found upregulated in both 5xFAD and clinical AD samples, demonstrating these are co-regulated at both gene and protein level. Furthermore, one of our astrocyte clusters was marked by a high phospho-Tau burden, which was significantly elevated in disease compared to control, and was higher in AD in DS compared to late-stage AD (Figure 7j). In sum, our IMC experiment in light of our other -omics experiments highlights the importance of a multi-faceted approach to comprehensively understand the altered molecular signatures of the brain in neurodegeneration.

### Integrating spatial transcriptomics with fluorescent amyloid imaging to identify pathology-associated gene expression signatures

Amyloid pathology is a defining feature of AD, and there is a critical need to understand the development of amyloid pathology and the cellular response. Previous studies identified amyloid-associated cell subpopulations that are transcriptionally distinct ^43–45^ as well as plaque-associated gene expression changes ^70^. However, these studies did not investigate amyloid-associated gene expression changes regarding the conformational changes associated with plaque formation. We sought to distinguish the transcriptional changes that may be conformation-dependent by staining each ST tissue section with Amylo-glo for dense amyloidbeta plaques and the antibody OC for diffuse amyloid fibrils and fibrillar oligomers (Figure 8a-b). This yielded information about the size and spatial location of amyloid aggregates in the same coordinates as our gene expression data, and we developed a custom image analysis pipeline to integrate these amyloid quantifications with our ST datasets (Methods). We found that the distribution of amyloid pathology in the spatial clusters was generally consistent with neuropathological plaque staging, where amyloid pathology is primarily in the cortical upper layers before spreading to the deeper layers (Figure 8c). Using the amyloid imaging data together with our ST data, we developed a consistent strategy to identify amyloid-associated gene expression changes in our human and mouse ST datasets (Methods). Briefly, we performed geospatial statistical analysis ^71, 72^ to identify “hot spots” of plaques and fibrils (Figures S31, S30, S31, S32), and used generalized linear models (GLMs) to perform differential gene expression testing with respect to the amyloid hot spot intensity (Methods). Critically, this analysis was agnostic of disease status or mouse age groups to specifically identify genes associated with amyloid rather than those associated with other disease-related factors. By performing a consistent analysis in our mouse and human datasets, we reasoned that we could assess the similarity of amyloid-associated gene sets between human AD and 5xFAD mice.

**Figure 8.**
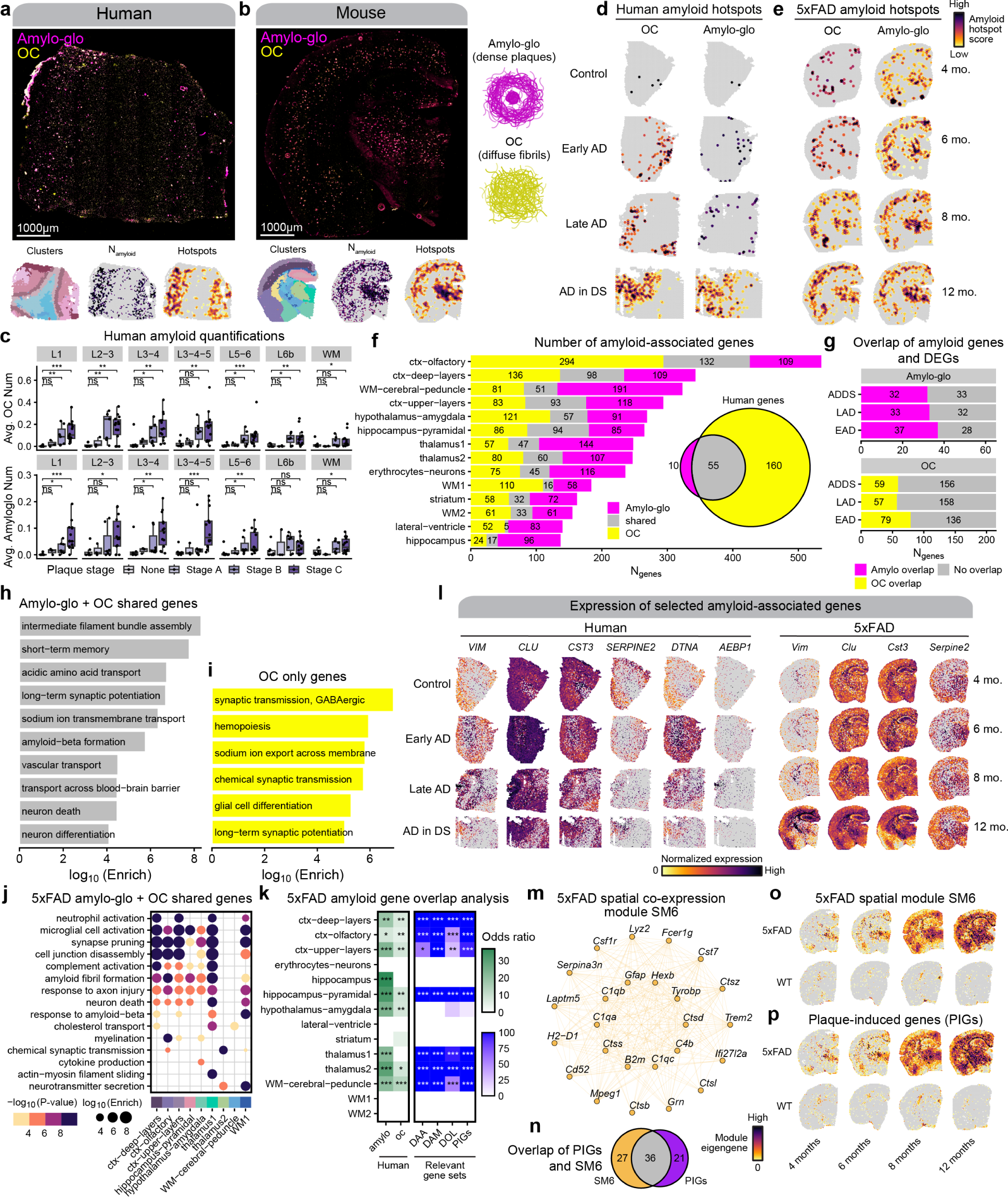
Identifying amyloid-associated gene expression signatures. **a-b**, Representative fluorescent whole-section images from one AD in DS (**a**) and one 12-month 5xFAD sample (**b**) stained with Amylo-glo and OC to mark dense amyloid plaques and diffuse amyloid fibrils respectively. ST data colored by cluster (left), amyloid quantification (middle), and amyloid hotspot analysis (right) are shown below the images. **c-d**, Spatial feature plots showing the amyloid hotspot analysis results (Getis-Ord Gi* statistic) for the human (**c**) and mouse (**d**) ST datasets for OC (left) and Amylo-glo (right). **e**, Box and whisker plots showing the distribution of amyloid quantifications in the human ST dataset, stratifying samples by their neuropathological plaque staging. Box boundaries and line correspond to the interquartile range (IQR) and median, respectively. Whiskers extend to the lowest or highest data points that are no further than 1.5 times the IQR from the box boundaries. **f**, Stacked bar plot showing the number of amyloid-associated genes from Amylo-glo, OC, and shared for each of the mouse ST clusters. Euler diagram shows the number of Amylo-glo- and OC-associated genes in the human dataset, and the overlap between these gene sets. g, Bar plots show the number of Amylo-glo- and OC-associated genes that overlap with disease DEGs. **h-i**, Selected pathway enrichment results from amyloid-associated genes that were shared between Amylo-glo and OC (**h**) and OC-specific (**i**) in the human ST dataset. **j**, Selected pathway enrichment results from amyloid-associated genes that were shared between Amylo-glo and OC for each cluster in the mouse ST dataset. **k**, Heatmap showing gene set overlap analysis results with the mouse amyloid-associated genes and human (left), and with other relevant gene sets (DAA: Disease-associated astrocytes ^44^; DAM: Disease-associated microglia ^43^; DOL: Disease-associated oligodendrocytes ^45^; PIGs: Plaque-induced genes ^70^). Fisher’s exact test results shown as follows: Not significant (ns), p > 0.05; * p <= 0.05; ** p <= 0.01; *** p <= 0.001, **** p<= 0.0001. **l**, Spatial feature plots of selected amyloid-associated genes in representative samples from the human (left) and mouse (right) ST datasets. **m**, Network plot showing module hub genes from the mouse spatial co-expression network module SM6. **n**, Euler plot showing the gene set overlap of the SM6 module and the PIGs module from Chen et al. ^70^. **o-p** Spatial feature plots showing the module eigengenes for the SM6 module (**o**) and the PIGs module (**p**) in representative samples of the mouse ST dataset.

Since amyloid depositions appear in grey matter much more frequently than in white matter, for the human dataset we focused our analysis on the gray matter regions (Methods). With this approach, we identified 65 amyloid plaqueassociated genes and 215 amyloid fibril-associated genes in the human dataset (Figure 8f, Table S25). Gene set overlap analysis showed that 55 genes were common between the plaqueand fibril-associated genes (Figure 8f, Fisher’s exact test p-value = 1.8e-115, odds ratio = 1248.92). Biological processes such as intermediate filament assembly, long-term potentiation, neuron death, and transport across the blood brain barrier were enriched in these shared genes (Figure 8g, Table S26). Notably, these genes included meta-module M11’s hub genes *CLU* and *VIM*. This significant overlap is not surprising as amyloid fibrils comprise plaques. However, the large number of genes exclusively associated with fibrillar amyloid represent transcriptional changes prior to plaque assembly, and are related to synaptic function and hemopoiesis (Figure 8h). A few genes were also members of M11, including *AEBP1* and *DTNA*, both previously found associated with AD ^73–75^. Further, we found that many, but not all, of these amyloid-associated genes are DEGs we identified in each diagnosis, demonstrating the increased spatial granularity of this analysis (Figure 8i).

In the mouse dataset, we identified amyloid-associated genes separately within each of the 14 spatial clusters. From this analysis, we found a total 1,829 plaque-associated genes and 1,759 fibril-associated genes (Figure 8f, Table S27). We found that some of the amyloid-associated gene sets had significant overlaps between those found across different spatial clusters, but most of these genes were spatially restricted to only one of these groups. Furthermore, we found broad overlaps between the sets of plaque and fibril-associated genes within each spatial cluster, with the largest overlaps found in the hippocampus pyramidal and the upper cortical layer clusters (Figure 8f). GO term enrichment analysis of mouse amyloid-associated genes associated these genes to immune and inflammation related processes like microglial activation, as well as neurodegeneration terms like axon injury, neuron death, and response to amyloid beta (Figure 8j, Table S28). Interestingly, genes associated only with OC staining included AD GWAS risk gene *Adamts7*, as well as immune-related genes, such as *Itgb2*, *Cd53*, and *Il33,* suggesting a unique immune signature preceding plaque formation. We next compared the amyloid-associated gene sets identified in this study with plaque-induced gene (PIG) module from another amyloid mouse model ^70^, in addition to other relevant gene signatures like disease-associated microglia (DAMs ^43^), astrocytes (DAAs ^44^), and oligodendrocytes (DOLs ^45^), which were originally identified in 5xFAD mice. We performed gene set overlap analysis, and we computed gene signature scores for these gene sets in the mouse and human ST dataset (Methods), revealing significant overlaps between our mouse amyloid-associated genes with these published gene sets (Figure 8k). Since the PIG module was previously identified using co-expression network analysis, we performed a similar network analysis in our 5xFAD ST dataset to see if we could recover a similar expression program using hdWGCNA ^46^ (Methods, Figures S33, S34, S35). One of the gene modules identified in this analysis (SM6) was enriched for genes related to microglial activation and amyloid-beta response, and the expression of this module increased with age in 5xFAD but not WT mice (Figure S33). Further, we found that this module significantly overlapped with the PIG module, and that the expression of the PIG module also only increased with age in 5xFAD mice (Figures S36, S37). Altogether, these findings demonstrate that these amyloid-associated gene expression changes are highly reproducible and are mouse model-agnostic.

We next directly assessed the overlap between the amyloid-associated genes identified in the human dataset with those from the mouse dataset to investigate the evolutionary conservation of amyloid-associated transcriptional changes (Figure 8k). We found modest yet significant overlaps between some of these gene sets, with a generally higher degree of overlap in the plaque-associated genes (Figure 8k). Shared OC-associated genes in both mouse and human cortex include neurofilaments *NEFH* and *NEFM*, as well as *ALDOC*, previously identified as amyloid-associated in cerebrospinal fluid ^76^, and *MAFB*, which was implicated in AD microglial gene regulation ^77^. Perhaps surprisingly, we noted a general lack of microglial genes in the human amyloid-associated genes, whereas several of these mouse amyloid gene sets, including those from cortical regions, contained microglial genes like *Trem2* and *Tyrobp*. However, we identified *MEF2C*, a microglial transcription factor associated with AD genetic risk ^65, 66^, in the shared amyloid genes. Furthermore, these shared amyloid genes highlighted meta-module M11 with the hub genes *VIM* and *CLU*. We found that M11’s gene expression signature mirrors amyloid deposition, indicating this meta-module represents species-conserved amyloid-associated biological processes (Figures S23, S24.

## Discussion

Single-cell sequencing has uncovered cell type and cell state-specific changes in disease, yielding novel insights into disease pathophysiology and gene targets for further study. However, a critical limitation of these methods is their inability to capture spatial information, and many disorders have spatially defined characteristics. AD pathology progresses in a predictably anatomical manner that may be linked to brain circuitry. We generated spatial transcriptomic data from postmortem human brain tissue samples of clinical AD, encompassing early- and late-stage AD in the general population, as well as AD in DS. Due to technical limitations, we could not profile gene expression at a macroscopic level of multiple brain regions, and our samples had varying amounts of cortical layers and white matter. We additionally generated spatial transcriptomic data from the 5xFAD mouse model of AD, allowing us to simultaneously assess multiple brain regions and the temporal dynamics of disease progression, with the added benefit of reduced sample heterogeneity. Both our human and mouse spatial datasets are the largest of our knowledge (n=119 total), altogether representing a rich data resource for the AD community.

We identified regional differential gene expression changes shared between AD in the general population and DS, in line with previous accounts of shared genetic, clinical, and biomarker features ^27, 78, 79^. Further, we generated the first snRNA-seq study of AD in DS and integrated previously published AD studies to identify cellular changes conserved between both populations. We leveraged our novel transcriptomic data of AD in DS to investigate the role of biological sex on the AD transcriptome in the DS population, revealing distinct biological signatures between males and females. In addition, identification of cross-species disease changes are of particular interest for mouse model development and preclinical translation. Therefore, we evaluated the expression of DEGs identified in clinical AD in the 5xFAD, identifying regional evolutionary-conserved disease gene expression changes. Furthermore, we integrated fluorescent imaging data to reveal amyloid-associated genes shared between both species. Although previous 5xFAD studies identified amyloid-associated subpopulations and genes, not all may be recapitulated in clinical AD, and this has been previously highlighted in human AD snRNA-seq studies examining the DAM signature. While this may be due to singlecell vs single-nucleus comparisons ^80^, a previous snRNA-seq study using human AD and 5xFAD samples ^14^ in addition to our previous re-analysis ^12^ demonstrated that this may be a species difference. In our present study, we present a list of species-conserved amyloid-associated genes, including previously identified and novel genes, to prioritize gene targets with high translational value and extend our knowledge of plaque formation by identifying conformation-specific gene expression changes. However, we also note that the present resolution of spatial transcriptomics (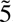5um) may have limited our findings.

In addition, we performed systems-level analyses using both our spatial and single-cell datasets to gain further insight into the AD transcriptome. We leveraged a newly published method, CellTrek ^56^, to provide spatial coordinates for our snRNA-seq populations and the cell-cell communication analytical package CellChat ^58^ to reveal disease-associated cellular communication changes. Specifically, changes in ANGPTL and CD99 signaling highlighted astrocyte modulation of brain vascular integrity in AD and pinpointed downstream targets of AD astrocyte phenotype changes. The multi-scale co-expression network analysis utilizing hdWGCNA ^46^ identified 166 gene modules across different cortical layers that we then refined into 15 cortex-wide “meta-modules”. Importantly, these modules exposed not only spatial, but also temporal patterns of gene expression; we found significant downregulation of modules specifically in early-stage AD, indicating dynamic shifts in gene expression that may play a pivotal role in the progression of AD. Comparison of these modules in 5xFAD mice also demonstrated these are cross-species AD dysregulated transcriptomic programs, and we integrated our snRNA-seq data to determine the cellular contexts of these spatial gene programs, uncovering both neuronal and glial components that may be integral to disease biology. In particular, the glial meta-module M11 is upregulated in the cortical upper layers with disease and contains DEGs shared between sporadic AD and AD in DS. The ligands *CD99* and *ANGPTL4*, identified in our cell signaling analyses, are hub genes in M11. We also found additional hub genes in our amyloid-associated genes, observing the expression of M11 at regions with amyloid deposition in both mouse and human samples, altogether emphasizing a critical role of M11’s associated biological processes and genes in AD pathophysiology.

Finally, we generated a spatial proteomic dataset to examine the protein expression of genes we identified in our transcriptomic datasets, in addition to other AD-related proteins. Notably we found that findings in the spatial transcriptomic data were upheld at the protein level; however, we could not confirm layer-specific expression due to technical limitations. snRNA-seq results were less concordant with the proteomic data. Although this may be due to profiling only nuclear RNA, resultant protein expression may also be affected by posttranscriptional and posttranslational regulatory changes, as well as protein trafficking, which may not be captured in transcriptomic data. High-throughput methods to assess protein expression are also still limited, and the availability of effective antibodies remains a large impediment in the study of proteins. In spite of these limitations, we found that the proteomic data confirmed that M11 represents key AD co-regulated genes, as we found the encoded proteins of both hub genes *CD44* and *CLU* upregulated in AD glia, demonstrating our integrative analytical approach uncovers functional information that enriches our understanding of disease-relevant gene targets.

Our work offers a pioneering exploration into the spatial and temporal dynamics of gene expression in the disease pathogenesis of both sporadic AD and AD in DS by using spatial transcriptomic data from postmortem human brain tissues and the 5xFAD mouse model. This study additionally highlights the significant roles of specific cell-types in AD and identifies key targets of cellular phenotype changes. The application of co-expression network analysis allowed the identification of spatial-temporal patterns of gene coexpression across different cortical layers, notably pointing to significant downregulation of certain gene modules during early-stage AD. This adds new dimensions to our understanding of AD’s pathophysiology, emphasizing the dynamic shifts in gene expression and the critical involvement of both neuronal and glial components in disease progression. Furthermore, the study presents a collection of species-conserved amyloid-associated genes, creating a foundation for future studies in AD and potentially facilitating the translation of in vitro and mouse model findings to clinical applications.

## ACKNOWLEDGEMENTS

Funding for this work was provided by National Institutes on Aging, Neurological Disorders and Stroke, and Drug Abuse grants 1RF1AG071683, P01NS084974-06A1, 1U01DA053826, U54 AG054349-06 (MODEL-AD), and 3U19AG068054-02S, Adelson Medical Research Foundation funds to V.S, T32AG000096-38 to E.M., and the National Institute on Aging predoctoral fellowship 1F31AG076308-01 to S.M. We thank the UCI Genomics Research and Technology Hub for providing their facilities and sequencing our single-nucleus RNA-seq and spatial transcriptomics libraries. This work utilized the infrastructure for high-performance and high-throughput computing, research data storage and analysis, and scientific software tool integration built, operated, and updated by the Research Cyberinfrastructure Center (RCIC) at the University of California, Irvine (UCI). We also thank Ram Stanciauskas and Ruby De Casas for their technical assistance.

## AUTHOR CONTRIBUTIONS

E.M, S.M., and V.S. conceptualized this study. The manuscript was written by E.M., S.M and V.S. with assistance and approval from all authors. E.M. generated the spatial transcriptomic data with assistance from C.M.H. and S.K.S. E.M. and Va.Sc. generated the spatial proteomic data with assistance from J.L. E.M. and S.K.S. generated the snRNA-seq libraries with assistance from C.M.H. and M.A.A. S.D. performed dissections and RNA isolation. S.M. performed bioinformatics and network analysis on human and mouse datasets with assistance from E.M., N.R., F.R., Z.S., and Z.C. Va.Sc. also helped with spatial proteomic data analyses. E.M., C.M.H., N.M., M.A.A., S.K.S. and V.S. processed mouse samples. N.M., S.S., C.M.H., and K.N.G. generated the mouse colony. N.M., S.S., C.M.H., and S.D. performed genotyping. S.W, I.T.L., J.S, E.D., W.H.Y., K.L, M.P-R and E.H. provided human brain samples from UCI ADRC.

## DATA AVAILABILITY

All raw and processed spatial transcriptomics and single-nucleus RNA sequencing data have been deposited into the National Center for Biotechnology Information Gene Expression Omnibus (GEO) database and will be made publicly available upon publication.

## CODE AVAILABILITY

The data analysis code used for this study is available on GitHub at https://github.com/swaruplabUCI/ADDS_paper.

## Methods

### Postmortem human brain tissue

Human brain tissue from prefrontal cortex and posterior cingulate cortex was obtained from UC Irvine’s Alzheimer’s Disease Research Center and the NIH NeuroBioBank under UCI’s Institutional Review Board (IRB). Samples were assigned to groups based on both NFT and plaque staging, in addition to clinical diagnoses. Samples were also selected based upon several covariates, including age, sex, race, postmortem interval (PMI), RNA integrity number (RIN), and disease comorbidity. RIN values were obtained by isolating total RNA with the Zymo Direct-zol RNA Isolation kit and running on the Agilent TapeStation 4200. Sample information is available in Table S1.

### Mouse brain tissue

All mouse work was approved by the Institutional Animal Care and Use (IACUC) committee at UCI. 5xFAD hemizygous (C57BL16) and wildtype littermates were bred and housed until sacrifice at 4, 6, 8, and 12 months. Sample information is available in Table S2. For genotyping, we used the following primers (for PSEN1): 5’ –AAT AGA GAA CGG CAG GAG CA –3’ (Forward), 5’ –GCC ATG AGG GCA CTA ATC AT –3’ (Reverse). Mice were euthanized by carbon dioxide inhalation. After PBS transcardiac perfusion, one brain hemisphere was flash frozen in isopentane chilled with dry ice for RNA analyses, while the other hemisphere was fixed in 4% paraformaldehyde for immunohistochemistry.

### Single-nucleus RNA-sequencing

Single-nucleus isolations were performed in randomized groups of 12 samples. 50mg fresh frozen postmortem human brain tissue was homogenized in Nuclei EZ Lysis buffer (NUC101-1KT, Sigma-Aldrich) and incubated for 5 min before being passed through a 70µm filter. Samples were then centrifuged at 500 g for 5 min at 4°C and resuspended in additional lysis buffer for 5 min. After another centrifugation at 500 g for 5 min at 4°C, samples were incubated in Nuclei Wash and Resuspension buffer (NWR, 1xPBS, 1% BSA, 0.2U/µl RNase inhibitor) for 5 min. To remove myelin contaminants, we prepared sucrose gradients with Nuclei PURE Sucrose Buffer and Nuclei PURE 2M Sucrose Cushion Solution from the Nuclei PURE Nuclei Isolation Kit (NUC-201, Sigma-Aldrich), and samples were carefully overlaid and centrifuged at 13,000 g for 45 min at 4°C. Samples were then washed in NWR before processing with the Nuclei Fixation Kit (SB1003, Parse Biosciences). After nuclei fixation and permeabilization, samples were cryopreserved with DMSO until day of library preparation. Library preparations were performed as 4 batches of 24 samples, with an additional batch to increase numbers of nuclei/sample for 16 samples. We generated single-nucleus libraries with the WTK Whole Transcriptome Kit (SB2001, Parse Biosciences). cDNA library quantification and quality were assessed with Qubit dsDNA HS assay kit (Q32851, Invitrogen) and D5000 HS kit (5067-5592, 5067-5593; Agilent) or D1000 HS kit (5067-5584, 5067-5585; Agilent) for the Agilent TapeStation 4200. Libraries were sequenced using Illumina Novaseq 6000 S4 platform using 100bp paired-end sequencing for a sequencing depth of 50,000 read pairs/cell.

### Spatial transcriptomics

Fresh frozen tissue samples were sectioned on a HM525NX cryostat (Fisher) at -15°C for 10µm thick sections that are immediately mounted onto 10× Genomics Visium slides. Slides were individually stored in slide mailers (sealed airtight in a plastic bag) at -80°C until staining. We followed 10× Genomics Methanol Fixation, Immunofluorescence Staining & Imaging for Visium Spatial Protocols (Rev C), except after tissue sections were fixed in methanol and blocked, the sections were incubated with Amylo-glo (1:100; TR-300, Biosensis) for 20 min. Sections were then incubated with the primary antibody OC (1:500 for mouse, 1:200 for human; polyclonal, AB2286, Millipore) and respective secondary antibody (1:400; goat anti-rabbit secondary antibody Alexa Fluor 488, Life Tech or Alexa Fluor 647, Life Tech). Immediately after immunostaining, capture areas were imaged on a widefield Nikon Ti2-E microscope at 20X magnification. Spatial transcriptomic libraries were then generated from the tissue sections according to the 10× Genomics Visium User Guide (Rev E). Library quantification, quality check, and sequencing were performed as previously described, but sequencing depth was based on an estimated 60% tissue area coverage per sample for 50,000 read pairs per covered spot.

### Spatial proteomics

Primary antibodies were formulated carrier-free except for YKL-40, which contained glycerol and was purified before metal conjugation with Amicon 10K Buffer Exchange Columns (UFC501096, EMD Millipore). All antibody concentrations were obtained using a Nanodrop 2000c Spectrophotometer and formulated with a final stock concentration of 0.5mg/ml. All antibodies were conjugated using Standard BioTool’s (SBT, formerly Fluidigm) Maxpar X8 metal conjugation protocol with Maxpar metal labeling kits (201300, SBT).

Fixed and cryoprotected tissue was sectioned on a HM525NX cryostat (Fisher) at -15°C for 14µm thick sections onto Fisher Superfrost Plus slides. Slides were stored at -80°C until staining, sealed airtight in a plastic bag. We followed the fresh frozen staining protocol from SBT; however, since the tissue was previously fixed, we skipped the fixation step. Slides were transferred on dry ice to incubate at 37°C for 5 min on a PCR machine, similar to the 10× Genomics Visium protocol. Sections were washed in PBS 3 times for 5 min before drawing a hydrophobic barrier. After the hydrophobic barrier dried, we incubated the sections with 3% BSA in PBS with 0.2% Triton X-100 for 45 min at room temperature. We then incubated the sections with the primary antibody cocktail diluted in 0.5% BSA/PBS with 0.2% Triton X-100 overnight at 4°C. The antibodies and dilutions used in the primary antibody cocktail are in Table S29. Sections were then washed in PBS with 0.2% Triton X-100 twice for 8 min before incubating with the iridium intercalator (1:100 in PBS; 201192A, SBT) for 30 min at room temperature. We then washed the sections in water twice for 5 min before allowing them to air dry before ablation. Hyperion Imaging System (SBT) was tuned prior to ablation using Hyperion Tuning Slide (201088, SBT) for optimal instrument performance. Ablations were performed with an ablation energy of four with a reference energy of zero in 1000×1000µm regions of interest, with the exception of one due to unexpected consumption of Argon gas that resulted in a 1000×922 acquisition.

### Preprocessing gene expression data

For the snRNA-seq dataset, we aligned sequencing reads to the reference transcriptome (GRCH38) and quantified gene expression using splitpipe (Parse Biosciences) in each of the five snRNA-seq experiments. We quantified and corrected ambient RNA signal present in our samples using Cellbender ^81^ remove-background (v 0.2.0). Heterotypic barcode collisions were inferred in each snRNA-seq experiment using Scrublet ^82^ (v 0.2.3) with default settings. We merged the individual snRNA-seq experiments into a single anndata (v 0.8.0) object, totaling 611,999 barcodes and 29,889 genes before additional quality control (QC) filtering. For each snRNA-seq experiment, we removed barcodes in the 95th percentile for number of UMIs detected, doublet score from Scrublet, and percentage of mitochondrial reads. We also applied dataset-wide cutoffs to remove barcodes with less than or equal to 250 UMIs, greater than or equal to 50,000 total UMIs, and greater than or equal to 10% mitochondrial reads. For one of the snRNA-seq experiments, we applied a more stringent filter to remove cells with less than or equal to 500 UMIs, and greater than or equal to 5% mitochondrial reads. We retained 431,534 barcodes for downstream analysis.

The 10X Genomics Loupe Browser image alignment tool was used to select Visium ST spots that intersected the tissue based on the fluorescent images. Sequencing reads from the human and mouse Visium experiments were processed using the 10X Genomics Spaceranger (v 1.2.1) pipeline, with GRch38 and MM10 as the respective reference transcriptomes. Spaceranger count was used to align sequencing reads to the reference, quantify gene expression, and perform a preliminary clustering analysis for each sample. Unlike the snRNA-seq dataset, we did not filter out additional spots based on sequencing QC metrics. The UMI counts matrices and fluorescent images for the human and mouse samples were combined into merged Seurat ^83–85^ objects for the respective species.

### Initial snRNA-seq data analysis

Following QC filtering, we processed the snRNA-seq dataset using SCANPY ^86^ and scVI ^87^. The UMI counts matrix was first normalized using the functions `sc.pp.normalize_total` and `sc.ppl.log1p`. We set up the anndata object to train the scVI model using snRNA-seq as the batch key and the following additional continuous and categorical covariates: sample ID, diagnosis, brain region, age at death, percentage of mitochondrial counts, number of UMI, postmortem interval (PMI) and RNA integrity number (RIN). We set up the scVI model with two hidden layers, 128 nodes per layer, a 30-dimensional latent embedding after the encoder phase, and a dropout rate of 0.1. We trained the model over 50 epochs and noted a flattened loss curve by the end of the training procedure. The latent embedding learned from the scVI model accounts for the batch effects and additional covariates specified in the model setup step, and we used this embedding for Leiden clustering and UMAP ^88^ dimensionality reduction in SCANPY. With a resolution parameter of 1.5 we identified 43 clusters. We inspected gene expression patterns in these clusters for a panel of canonical CNS cell type marker genes to assign major cell type labels to each cluster. We also checked the distribution of QC metrics in each cluster to identify outlier clusters. Six clusters (7, 29, 33, 35, 50, 51) were removed from downstream analysis as QC outliers, or due to presence of potential doublets. We recomputed the UMAP and Leiden clustering (resolution = 1.2) after filtering these clusters, yielding 29 clusters. Glutamatergic neuron clusters were annotated based on expression of known cortical layer marker genes, and GABAergic neuron clusters were annotated based on expression of known markers (*VIP*, *SST*, *PVALB*, *LAMP5*). At this stage, non-neuronal cell clusters were simply labeled by their major cell type (astrocytes, microglia, oligodendrocytes, oligodendrocyte progenitors, vascular cells). To identify subpopulations in non-neuronal cells, we performed subclustering analysis in each of the major non-neuronal cell populations (microglia, astrocytes, oligodendrocytes, vascular cells). Each group was isolated in its own anndata object, and Leiden clustering was performed (see GitHub repository for subclustering parameters).

### Reprocessing publicly available single-nucleus gene expression datasets

We obtained sequencing data from obtained three published snRNA-seq studies ^12–14^ of AD. Sequencing data from the Mathys et al. 2019 and Zhou et al. 2020 datasets were downloaded from Synapse (syn18485175 and syn21670836), and the data from the Morabito et al. 2021 study generated by our own group was not re-downloaded. We used a uniform pipeline to process each of these datasets, with slightly varying parameters which are noted in our GitHub repository. This pipeline and the resulting anndata objects are identical to those used in another study from our group ^46^, and we reiterate the main analysis steps here. Sequencing reads were pseudoaligned to the reference transcriptome (GRch38) and gene expression was quantified using the count function from kallisto bustools ^89^. Ambient RNA signal was corrected in UMI counts matrices for each sample using Cellbender ^81^ remove-background, and we used Scrublet ^82^ to identify barcodes attributed to more than one cell. Individual samples were then merged into one anndata object for each of the three studies. Analogous to the snRNA-seq data we generated in this study, we performed percentile filtering based on the following QC metrics: doublet score, number of UMI per cell, and percentage of mitochondrial reads per cell. The downstream processing was performed using SCANPY ^86^. Gene expression was normalized using the functions `sc.pp.normalize_total` and `sc.pp.log1p`, resulting in a ln(CPM) transformation of the input UMI counts data. Highly variable features were identified using `sc.pp.highly_variable_genes`, which were then scaled to unit variance and centered at zero using sc.pp.scale. Linear dimensionality reduction was performed on the scaled expression matrix using PCA with the function sc.tl.pca. Harmony ^90^ was used to batch correct the PCA representation with the function `sc.external.pp.harmony_integrate`. A cell neighborhood graph was computed based on the harmony representation using sc.pp.neighbors, followed by Leiden ^30^ clustering and non-linear UMAP dimensionality reduction with sc.tl.leiden and sc.tl.umap respectively. Canonical CNS cell type marker genes were used to assign coarse-grain identities to each cluster, and to identify additional doublet clusters that passed our previous filtering steps. We inspected the distribution of the QC metrics in each cluster and removed outlier clusters. After filtering additional low-quality clusters, we ran UMAP and Leiden clustering again to result in the final processed anndata object for each dataset.

### Spatial transcriptomics clustering analysis

In the human and mouse ST datasets, we grouped spots into biologically relevant clusters by accounting for transcriptome measurements and spatial coordinates. The BayesSpace ^31^ clustering algorithm uses a low-dimensional representation of the transcriptome with a spatial prior to encourage assignment of neighboring spots in the same cluster. Critically, BayesSpace produces a single unified clustering across many different ST experiments, rather than separate clustering and annotation for each ST slide. Seurat objects were converted to the SingleCellExperiment format using the function as.SingleCellExperiment. Absolute and relative spatial coordinates were stored in the meta-data compartment of the SingleCellExperiment objects to inform the BayesSpace model of the spatial information, ensuring to offset each sample such that there was no overlap. Each dataset was log normalized and linear dimensionality reduction was performed with PCA using the function spatialPreprocess from the BayesSpace R package. Harmony ^90^ batch correction was applied on the basis of individual samples using the RunHarmony function. For the human dataset, we ran BayesSpace clustering using the spatialCluster function in the BayesSpace R package, varying the q parameter (number of resulting clusters) from 5 through 10. We inspected the output of each clustering and found that q=9 produced results that were most consistent with the underlying anatomy of the cortex, allowing us to annotate clusters based on cortical layers and white matter. Similarly, we ran BayesSpace clustering on the mouse dataset varying the q parameter between 10 and 20, and we selected q=15 for downstream analysis.

### Reference-based integration of snRNA-seq datasets

We performed reference-based integration of the snRNA-seq dataset from the current study with the three published AD snRNA-seq datasets. Using our new snRNA-seq dataset as the reference, we projected the three published datasets into the reference latent space using scANVI ^33^, and we performed transfer learning to predict cell identities using scArches ^91^. While scANVI shares similarities with the scVI model that we previously used to process our snRNA-seq data, it is a semi-supervised model that leverages cell annotations in the reference dataset to inform the latent representation of the query dataset. We trained the scANVI model separately for each of the query datasets using the class scvi.model.SCANVI, training for 100 epochs in each case. For each query dataset, this process resulted in a low-dimensional representation of the transcriptome in the latent space originally learned from the reference snRNA-seq dataset with the `model.get_latent_representation` function, and predicted cell annotation labels from the model.predict function. We merged the reference dataset with the three query datasets, and we ran UMAP on the scANVI latent representation to visually represent the unified dataset in two dimensions.

### Cluster marker gene analysis

We performed cluster marker gene analysis for each snRNA-seq cluster using the snRNA-seq dataset generated in this study. For this analysis, we employed a “one-versus-rest” strategy to systematically perform differential gene expression analysis comparing each cluster to all the other clusters. We used MAST ^92^ as our differential expression model for this test, accounting for the sequencing batch and the number of UMI per nuclei as covariates in our model. We a similar strategy to perform cluster marker gene analysis in our human and mouse Visium spatial transcriptomics datasets where we used the biological sample ID and the number of UMI per spot as model covariates for MAST.

### Differential gene expression analysis

We systematically performed differential gene expression analysis in each of our datasets to compare the disease conditions with controls. For all our differential gene expression tests, we use MAST ^92^ as the underlying model, which has shown favorable results in recent benchmarks for differential expression analysis in datasets with multiple sequencing batches98. For the snRNA-seq dataset generated in this study, we compared gene expression between the AD in DS and cognitively normal control groups for each cluster and major cell type, and we performed this analysis separately for the two brain regions profiled in this study (frontal cortex and posterior cingulate cortex). We used the sequencing batch, number of UMI per nuclei, sample postmortem interval (PMI), and sample RNA integrity number (RIN) as model covariates for these tests. We used a similar strategy for the differential expression analysis of the previously published snRNA-seq datasets ^12–14^ to compare late-stage AD samples with cognitively normal controls, accounting for study of origin and number of UMI per nuclei as model covariates. We compared gene expression between cognitively normal controls and the three experimental groups (early-stage AD, late-stage AD, and AD in DS) separately in our human spatial transcriptomics dataset. For these comparisons, we used RIN, PMI, number of UMI per spot, date of sequencing library preparation, and sequencing batch as model covariates. In the mouse spatial transcriptomics dataset, we performed differential gene expression analysis to compare gene expression between 5xFAD and wild type mice in each ST cluster, and we stratified this analysis by each age group (4, 6, 8, and 12 months). For the mouse analysis, we used the number of UMI per spot, sequencing batch, date of sacrifice, and date of sequencing library preparation as model covariates. We visualized the results of the differential expression tests as volcano plots to show the statistical significance and the effect sizes for each gene. For the human datasets, we also visualized the differential expression results using a heatmap stratified by chromosome to inspect the contribution of each chromosome to the overall set of DEGs.

Within the human snRNA-seq and ST datasets, we compared the results of the differential expression analyses across the different conditions within each snRNA-seq and ST group. For these comparisons we computed Pearson correlations and performed linear regression on the differential expression effect sizes for the set of genes that were significantly different between condition and control for either of the analyses. We visualized these results using scatter plots, and indicated genes that were consistent or inconsistent in terms of their effect sizes across the different comparisons. We used the EnrichR ^93^ R package (version 3.1) to identify biological pathways and processes that are enriched in our sets of DEGs, and in sets of DEGs that were shared across different differential expression tests.

### Hierarchical co-expression network analysis in human spatial transcriptomics

We performed hierarchical coexpression network analysis across different cortical layers and white matter in our human ST dataset using the R package hdWGCNA ^46, 94^ (version 0.2.19). Prior to network analysis, we computed pseudo-bulk gene expression profiles for each of our 39 ST samples in the for the cortical layer clusters and white matter (L1, L2-3, L3-4, L3-4-5, L5-6, L6b, WM), and we calculated log2(CPM) (counts per million mapped reads) normalized expression values. Genes were retained for network analysis if they were expressed above 0 UMI counts in at least 5% of cells in any of the spatial annotations, thereby retaining 10,199 genes. We selected soft-power threshold parameters for each spatial region by computing the scale-free topology model fit at for different soft-power values, and using the smallest parameter which yielded a model fit greater than 0.8. Co-expression networks were then computed separately for the seven spatial regions using the hdWGCNA function ConstructNetwork, using a minimum module size of 50, a merge cut height of 0.1 for dynamic tree cutting ^95^, the soft power thresholds as previously described, and all other parameters set to the default values. This process yielded 166 gene co-expression modules across the seven spatial regions, and we assigned a unique name to each module based on a combination of their spatial region of origin and a numeric identifier. For example, module “L1-M7” is the seventh module originating from spatial region L1. We calculated gene expression summary values, called “module eigengenes” (MEs) for each of these 166 modules in each of the ST spots using the hdWGCNA function ModuleEigengenes, applying a Harmony95 correction to the MEs based on biological sample of origin. We then computed eigengene-based connectivity (kME) for each gene using the hdWGCNA function ModuleConnectivity.

We next sought to perform a hierarchical analysis of these co-expression networks using a strategy similar to a previous study ^47^. First, we calculated similarity metrics between pairs of co-expression modules that arose from different spatial regions by computing gene overlap statistics such as Jaccard similarity (J) using the R package GeneOverlap (version 1.34.0). Next, we calculated pairwise Pearson correlations between each of the 166 co-expression modules within each of the seven brain regions, and we kept the component-wise maximum of these correlations (E). Using these two measures of similarity, we computed a module-module dissimilarity matrix D = 1 –(E + (3J)/4). We then computed Euclidean distance between modules in this matrix and performed hierarchical clustering with the R function hclust. We refer to the 15 groups of modules that arose from this hierarchical clustering analysis as “meta-modules”. Similar to our previous co-expression network analysis, we computed MEs and kMEs for these meta-modules to summarize gene expression in the ST spots and to quantify the eigengene-based connectivity of each gene. For cases where a single gene was assigned to more than one meta-module, we re-assigned the gene to only the meta-module where the gene had the highest eigengene-based connectivity. Additionally, we computed a consensus topological overlap matrix (TOM) based on the seven TOMs from the co-expression analysis in each spatial regions, and we visualized this consensus TOM as a UMAP plot using the hdWGCNA function RunModuleUMAP. We identified enriched biological processes in each of the meta-modules using the EnrichR ^93^ R package (version 3.1) to overlap these gene sets with those associated via Gene Ontology (GO).

We performed differential module eigengene (DME) analysis to compare the differences in the expression of each coexpression module between disease groups (early-AD, late-AD, and AD in DS) with cognitively normal controls. This analysis was done using the hdWGCNA function FindDMEs, using a Wilcoxon rank sum test for the comparisons. Like differential gene expression analysis, DME analysis results in measures of statistical significance and effect sizes for each co-expression module across each comparison. For the 166 region-specific co-expression modules, we performed DME analysis only within the region that the modules were derived from. Alternatively, for the meta-modules we performed DME analysis within each of the spatial regions. After performing these tests, we sought to compare the results of these tests across the different disease conditions to identify similarities and differences among the co-expression patterns. For the 166 co-expression modules, we computed Pearson correlations and linear regressions comparing the effect sizes of the DME results and visualized the results as scatter plots. We next tested the overlap between sets of co-expression modules that were significantly differentially expressed between the groups using the UpSetR ^96^ package.

To inspect the activity of these spatially derived co-expression networks in other related contexts, we projected the 166 co-expression modules and the 15 meta-modules into our snRNA-seq and mouse ST datasets using the hdWGCNA function ProjectModules. This function computes MEs in a query dataset for sets of genes from co-expression modules that were found in a different reference dataset. In this case, the reference dataset is the human ST dataset and the query datasets are the human snRNA-seq and the mouse ST datasets. We coarsely inspected the distributions of the meta-modules in specific cell types within the human snRNA-seq dataset with UMAP visualizations where each cell is colored by the projected ME value. In the mouse ST dataset, we inspected these trends using violin plots stratified by age group and genotype. Furthermore, in the mouse dataset performed module preservation analysis ^48^ to assess the reproducibility of these modules across species using the hdWGCNA function ModulePreservation.

### Co-expression network analysis in mouse spatial transcriptomics

We used hdWGCNA ^46^ to perform a brain-wide gene co-expression network analysis in our mouse ST dataset, similar to the analysis from the mouse brain ST dataset from Chen et al. ^70^ which was used to identify the “plaque-induced gene” (PIG) network. First, since co-expression network analysis is sensitive to noise in sparse datasets, we computed “meta-spots” by merging the transcriptomes of adjacent ST spots in a grid pattern for each of the 80 ST samples. Genes were retained for network analysis if they were expressed above 0 UMI in at least 5% of spots originating from any of the different brain regions, yielding a total of 12,579 genes. After performing a soft-power threshold search similar to our human co-expression network analysis, we computed the co-expression network topological overlap matrix (TOM) and identified gene modules using the hdWGCNA function ConstructNetwork with default parameters. In total, this process yielded 10 brain-wide spatial co-expression modules. We computed module eigengenes (MEs) for these modules using the hdWGCNA function ModuleEigengenes, applying a Harmony ^90^ correction to the MEs based on the sequencing batch. Next, we computed eigengene-based connectivity (kME) for each gene using the hdWGCNA function ModuleConnectivity. We visualized the co-expression network in two dimensions by computing a UMAP representation of the TOM with the hdWGCNA function RunModuleUMAP. We used the EnrichR ^93^ R package (version 3.1) to identify biological processes from Gene Ontology (GO) that were enriched in these co-expression modules.

We sought to compare these co-expression modules with other relevant gene sets. For this analysis we used the same gene sets described in the Quantifying gene expression signatures of disease-relevant gene sets section. We performed pairwise Pearson correlations of the MEs with the UCell ^37^ scores of these gene signatures to assess the similarity of gene expression patterns using the hdWGCNA function ModuleTraitCorrelation. We next computed gene overlap statistics between the sets of genes in each co-expression modules with these other gene sets using the R package GeneOverlap. To further our comparison with the spatial co-expression modules from the Chen et al. study, such as the PIG module, we used the hdWGCNA function ProjectModules to quantify gene expression patterns of the Chen et al. modules in our mouse ST dataset. We then computed pairwise Pearson correlations between the MEs for the co-expression modules derived from our mouse ST dataset to those derived from the Chen et al. dataset.

### Sex differences within the AD in DS cohort

We performed additional differential gene expression tests to identify sex differences within the AD in DS cohort, using a similar strategy as the other differential expression tests with MAST ^92^ as the underlying model. For the snRNA-seq dataset, we compared expression between nuclei from female versus male samples in each cell type and each cell cluster, and we used sequencing batch, the number of UMI per nuclei, and the postmortem interval (PMI) as the model covariates. Since the spatial transcriptomics dataset had a greater number of female samples (7) as compared to male (3), before running the differential expression analysis we first down sampled the dataset (stratified by biological sample) such that the number of spots from the female samples matched that from the male samples. We then performed differential expression analysis with MAST using the number of UMI per spot and the PMI as the model covariates. We inspected the overlap between sets of DEGs between the different spatial regions using the R package UpSetR ^96^ (version 1.4.0). We used the EnrichR R package to identify biological processes enriched in DEGs in each spatial region. To complement the differential expression analysis, we also used the hdWGCNA R package to perform differential module eigengene (DME) analysis to compare the expression of our co-expression modules between female and male samples in each spatial region.

### Spatial mapping of snRNA-seq data

We mapped our snRNA-seq dataset into spatial coordinates using the R package CellTrek ^56^ (v 0.0.94). Briefly, the CellTrek pipeline enables spatial mapping of single-cell transcriptomes by creating an integrated co-embedding of ST and single-cell data, followed by a multivariate random forest model to predict the biological coordinates from the shared feature space. In our testing, we found that this algorithm was limited in that it could not scale to large datasets comprising hundreds of thousands of single cells. Additionally, this algorithm only maps data to a single ST slide at a time. We also found that the CellTrek algorithm only provided predicted coordinates for a subset of the input single cell transcriptomes. For these reasons, we mapped our snRNA-seq frontal cortex data to the human ST dataset in a pairwise fashion for each snRNA-seq sample and each ST sample. For a given pair of ST and snRNA-seq samples, we constructed an integrated co-embedding using the CellTrek function `traint` with default parameters. We then iteratively mapped the single-cell transcriptomes into the ST coordinates using the celltrek function over three iterations. The second iteration only included cells that were not mapped in the first iteration, and the third iteration only included cells that were not mapped in the first or second iterations. We then computed the Euclidean distance between each mapped cell and each of the ST spots, and we labeled each cell with a spatial annotation based on the most frequently observed annotation among the labels of the ten closest spots. After running the pairwise CellTrek mappings, we compiled the results into a single table. In sum, this process yielded multiple spatial coordinates and multiple annotations for each cell across the 39 human ST samples in this study. Given that these tissue samples varied in their grey and white matter content, the CellTrek mappings and inferred spatial annotations are generally not consistent across the ST samples. To come up with a consensus regional annotation across the different spatial mappings, we excluded the mappings from ST samples which were excessively high in white matter or grey matter content. We computed a metric summarizing the grey to white matter ratio in each ST sample by counting the number of grey matter spots and white matter spots, taking the difference, and dividing by the total number of spots. Positive values indicate higher grey matter content, while negative values indicate higher white matter content. We excluded samples with greater than 0.9 and less than -0.3, thereby retaining mappings from 34 of the ST samples. For each cell, we counted the number of times it was mapped to each spatial region, and labeled the cell based on the most frequently mapped region across the different samples. We further simplified these spatial annotations by upper cortical, lower cortical, or white matter regions.

### Cell-cell signaling analysis

We performed cell-cell signaling analysis in our snRNA-seq frontal cortex dataset with CellChat ^58^ (v 1.1.3), using the predicted spatial annotations in addition to cell type labels. The human CellChatDB ligandreceptor interaction database was used for this analysis. To facilitate downstream comparisons of the signaling networks in DSAD versus control samples, we ran the CellChat workflow separately based on disease status. The CellChat object was created using the normalized gene expression matrix and the cell type annotations with the predicted spatial regions from CellTrek, removing any cell groups with fewer than 30 cells. We then ran the recommended CellChat workflow using the following functions: `identifyOverExpressedGenes`, `identifyOverExpressedInteractions`, `projectData`, `computeCommunProb`, `filterCommunication`, `subsetCommunication`, `computeCommunProbPathway`, `aggregateNet`, and `netAnalysis_computeCentrality`. The DSAD and control CellChat objects were merged into one object using the mergeCellChat function. We compared the signaling networks across conditions both functionally and structurally using the computeNetSimilarityPairwise function. Furhter, we used the rankNet function to compute the relative information flow changes between AD in DS and control across all signaling pathways. We identified differentially expressed ligands and receptors as well as their signaling pathways using the identifyOverExpressedGenes function, visualizing selected results with the `netVisual_bubble` function.

### Quantifying gene expression signatures of disease-relevant gene sets

We used the UCell ^37^ R package (version 2.2.0) to quantify gene expression signatures of several relevant gene sets with the function `AddModuleScore_UCell`. The following gene sets were used for this analysis; homeostatic microglia ^43^, disease-associated microglia ^43^, disease-associated astrocytes ^44^, disease-associated oligodendrocytes ^45^, plaque-induced genes ^70^. The full list of genes within each gene set used for this analysis can be found on our GitHub repository.

### Integration of amyloid imaging data and spatial transcriptomic data

Since we stained the brain sections used for ST with amylo-glo and OC, we developed a custom data analysis pipeline to identify gene expression changes associated with amyloid-beta plaque depositions in our human and mouse ST datasets. For this analysis, our data processing pipeline was uniform among the human and mouse datasets. We used custom automated imaging analysis protocols (General Analysis protocols on NIS-Elements) to obtain Amylo-glo+ and OC+ binaries thresholding by intensity and size, as well as accounting for autofluorescence/non-specific staining by negative thresholding based on an empty channel. We exported the following values for each binary: Area (µm2), Diameter, center X and Y coordinates. Only 5xFAD samples were used for the mouse samples, as there is no amyloid pathology in WT mice. Samples with high background were excluded. The image analysis of the amylo-glo and OC fluorescent images gives us the coordinates and sizes of stained amyloid bodies which can then be directly compared to the Visium ST data. We separately counted the number of amyloid aggregates stained with amylo-glo or OC that overlapped each of the ST spots. We then calculated the number of Amylo-glo or OC+ binaries per spot by testing for an intersection between a spot and binary.

The radius of a spatial spot was calculated according to values provided by 10× Genomics, where a spot is 55µm with a 100µm distance between spot centers, and expanded the radius of a spot to account for the gap between spots. To account for the size of each amyloid aggregate, we also computed the sum of the areas of all amyloid aggregates overlapping each ST spot. We next used the R package Voyager ^97^ (version 1.0.10) to perform hot spot analysis by computing the Getis-Ord Gi* ^71, 72^ statistics for the amylo-glo and OC area scores in each ST sample.

### Identifying amyloid-associated gene expression signatures in human and mouse

We used the Getis-Ord Gi* ^71, 72^ hot spot statistics for amyloid aggregates stained with amylo-glo and OC to identify gene expression signatures associated with amyloid aggregation in the human and mouse ST datasets. We used generalized linear models (GLMs) to identify genes that were significantly altered in expression with respect to the amyloid hot spot statistics. This analysis was done using the `fit_-models` function from the R package monocle3^98^ (version 1.3.1). We used the biological sample of origin and the number of UMI per spot as model covariates. Further, this analysis was performed separately for the amylo-glo and OC hot spot statistics, since these two stains identify different forms of amyloid aggregates. Since our mouse brain dataset profiled an entire brain hemisphere and the clusters broadly corresponded to different major brain regions, we also performed this analysis separately for each of the clusters in the mouse ST dataset. Alternatively, the human dataset contains ST profiles only in the frontal cortex grey matter and white matter. Amyloid aggregation tends to primarily occur in the grey matter, as seen in our hot spot analysis, and for this reason we chose to exclude the human white matter ST spots from the analysis. After running the GLM, we computed Pearson correlations between the amyloid hot spot scores and gene expression for the significant results. We consider genes to be amyloid-associated if there was a significant result from the GLM (FDR < 0.05) and a positive correlation between gene expression and the amyloid score.

For each of these sets of amyloid-associated genes, we performed biological pathway enrichment analysis using the R package EnrichR ^93^ (version 3.1) with gene sets from the Gene Ontology database. In both the human and mouse datasets, we computed overlap statistics between the sets of amyloid-associated genes from amylo-glo and OC using the R package GeneOverlap (version 1.34.0). Further, we computed overlap statistics between each set of amyloid-associate genes and other disease relevant gene sets which were previously described in the Quantifying gene expression signatures of disease-relevant gene sets section. We also performed a gene overlap analysis to compare the set of amyloid-associated genes between the human and mouse datasets.

### Spatial proteomics data analysis

SBT MCD files were imported into Visiopharm Software version 2022.03. Image classes were created for training and included background, nuclei, and nuclei boarder. Nuclei detect AI App (version 2023.01.2.13695), a pre-trained deep learning app developed by Visiopharm was used to detect nuclei with Ir191 and Ir193 nuclear channels. Single cell data was exported to a .tsv file for further analysis.

We performed an unbiased clustering analysis of our imaging mass cytometry (IMC) spatial proteomic dataset using the R package Seurat ^83–85^ (version 4.3.0). We first created a Seurat object using the protein intensity by nuclei segments matrix as the input, and then we log-normalized this matrix using the Seurat function `NormalizeData`. We next performed dimensionality reduction by scaling and centering the data with the `ScaleData` function, performing principal component analysis (PCA) with the `RunPCA` function. To correct for the sample-specific differences in our protein intensity data, we ran Harmony ^90^ to correct the PCA matrix prior to running Louvain clustering and UMAP. We then performed a one-versus-rest marker test (Wilcoxon rank sum test) with the Seurat function `FindAllMarkers` to identify proteins which were significantly expressed in each cluster in order to annotate them with cell-type labels. Following our cell-type annotation, we performed additional Wilcoxon rank sum tests to compare the different experimental groups (cognitively normal controls, late-stage AD, AD in DS) within each cluster for each protein in our panel.

**Figure S1. Quality control of the human and mouse spatial transcriptomics datasets.**

**Figure S2. Clustering results in the human ST dataset.**

**Figure S3. Clustering results in the mouse ST dataset.**

**Figure S4. Quality control of the integrated snRNA-seq dataset.**

**Figure S5. Early-stage AD versus control DEGs in the ST dataset.**

**Figure S6. Late-stage AD versus control DEGs in the ST dataset.**

**Figure S7. AD in DS versus control DEGs in the ST dataset.**

**Figure S8. Late-stage AD versus control DEGs in the snRNA-seq dataset.**

**Figure S9. AD in DS versus control DEGs in the FCX snRNA-seq dataset.**

**Figure S10. AD in DS versus control DEGs in the PCC snRNA-seq dataset**

**Figure S11. Comparison of snRNA-seq AD in DS versus control DEG effect sizes in the PCC and FCX**

**Figure S12. snRNA-seq DEGs examined by chromosome**

**Figure S13. Spatial transcriptomic DEGs examined by chromosome**

**Figure S14. Comparison of spatial transcriptomic DEG effect sizes between early-stage AD and late-stage AD**

**Figure S15. Comparison of snRNA-seq DEG effect sizes between sporadic AD and AD in DS**

**Figure S16. 5xFAD versus wild type DEGs at four months of age.**

**Figure S17. 5xFAD versus wild type DEGs at six months of age.**

**Figure S18. 5xFAD versus wild type DEGs at eight months of age.**

**Figure S19. 5xFAD versus wild type DEGs at twelve months of age.**

**Figure S20. Number of 5xFAD DEGs and GO term enrichment analysis.**

**Figure S21. Comparisons of human and mouse DEGs.**

**Figure S22. Module hub gene networks for the human ST co-expression meta-modules.**

**Figure S23. Module eigengenes for the human ST meta-modules.**

**Figure S24. Module eigengenes of human co-expression meta-modules in the mouse ST dataset.**

**Figure S25. Cell-type deconvolution results in the human ST dataset.**

**Figure S26. Cell-type deconvolution results in the mouse ST dataset.**

**Figure S27. Differential cell-cell signaling between AD in DS and control.**

**Figure S28. CD99 signaling changes between AD in DS and control.**

**Figure S29. Integration of amyloid plaque imaging (Amylo-glo) and hotspot analysis in the human ST dataset**

**Figure S30. Integration of amyloid fibril imaging (OC) and hotspot analysis in the human ST dataset.**

**Figure S31. Integration of amyloid plaque imaging (Amylo-glo) and hotspot analysis in the mouse ST dataset**

**Figure S32. Integration of amyloid fibril imaging (OC) and hotspot analysis in the mouse ST dataset**

**Figure S33. Co-expression network analysis in the mouse ST dataset**

**Figure S34. Module hub gene networks from the mouse co-expression network analysis**

**Figure S35. Module eigengenes in representative samples from the mouse ST dataset**

**Figure S36. Module-trait correlation analysis in the mouse co-expression network**

**Figure S37. Investigating the gene co-expression modules from Chen et al in our 5xFAD ST dataset**

## Bibliography

1. BRAIN Initiative Cell Census Network (BICCN), BRAIN Initiative Cell Census Network (BICCN) Corresponding authors, Edward M Callaway, Hong-Wei Dong, Joseph R Ecker, Michael J Hawrylycz, Z Josh Huang, Ed S Lein, John Ngai, Pavel Osten, et al. A multimodal cell census and atlas of the mammalian primary motor cortex. Nature, 598(7879):86–102, 2021. ISSN 0028-0836. doi: 10.1038/s41586-021-03950-0.

2. Blue B. Lake, Rizi Ai, Gwendolyn E. Kaeser, Neeraj S. Salathia, Yun C. Yung, Rui Liu, Andre Wildberg, Derek Gao, Ho-Lim Fung, Song Chen, et al. Neuronal subtypes and diversity revealed by single-nucleus RNA sequencing of the human brain. Science, 352(6293):1586–1590, 2016. ISSN 0036-8075. doi: 10.1126/science.aaf1204.

3. Spyros Darmanis, Steven A. Sloan, Ye Zhang, Martin Enge, Christine Caneda, Lawrence M. Shuer, Melanie G. Hayden Gephart, Ben A. Barres, and Stephen R. Quake. A survey of human brain transcriptome diversity at the single cell level. Proceedings of the National Academy of Sciences, 112(23):7285–7290, 2015. ISSN 0027-8424. doi: 10.1073/pnas1507125112.

4. Rebecca D. Hodge, Trygve E. Bakken, Jeremy A. Miller, Kimberly A. Smith, Eliza R. Barkan, Lucas T. Graybuck, Jennie L. Close, Brian Long, Nelson Johansen, Osnat Penn, et al. Conserved cell types with divergent features in human versus mouse cortex. Nature, 573 (7772):61–68, 2019. ISSN 0028-0836. doi: 10.1038/s41586-019-1506-7.

5. Bosiljka Tasic, Vilas Menon, Thuc Nghi Nguyen, Tae Kyung Kim, Tim Jarsky, Zizhen Yao, Boaz Levi, Lucas T Gray, Staci A Sorensen, Tim Dolbeare, et al. Adult mouse cortical cell taxonomy revealed by single cell transcriptomics. Nature Neuroscience, 19(2):335–346, 2016. ISSN 1097-6256. doi: 10.1038/nn.4216.

6. Bosiljka Tasic, Zizhen Yao, Lucas T. Graybuck, Kimberly A. Smith, Thuc Nghi Nguyen, Darren Bertagnolli, Jeff Goldy, Emma Garren, Michael N. Economo, Sarada Viswanathan, et al. Shared and distinct transcriptomic cell types across neocortical areas. Nature, 563(7729): 7729–72, 2018. ISSN 0028-0836. doi: 10.1038/s41586-018-0654-5.

7. Zizhen Yao, Cindy T.J. van Velthoven, Thuc Nghi Nguyen, Jeff Goldy, Adriana E. Sedeno-Cortes, Fahimeh Baftizadeh, Darren Bertagnolli, Tamara Casper, Megan Chiang, Kirsten Crichton, et al. A taxonomy of transcriptomic cell types across the isocortex and hippocampal formation. Cell, 184(12):3222–3241.e26, 2021. ISSN 0092-8674. doi: 10.1016/j.cell.2021.04.021.

8. Alexander B. Rosenberg, Charles M. Roco, Richard A. Muscat, Anna Kuchina, Paul Sample, Zizhen Yao, Lucas T. Graybuck, David J. Peeler, Sumit Mukherjee, Wei Chen, et al. Single-cell profiling of the developing mouse brain and spinal cord with split-pool barcoding. Science, 360(6385):176–182, 2018. ISSN 0036-8075. doi: 10.1126/science.aam8999.

9. Amit Zeisel, Ana B. Muñoz-Manchado, Simone Codeluppi, Peter Lönnerberg, Gioele La Manno, Anna Juréus, Sueli Marques, Hermany Munguba, Liqun He, Christer Betsholtz, et al. Cell types in the mouse cortex and hippocampus revealed by single-cell RNA-seq. Science, 347(6226):1138–1142, 2015. ISSN 0036-8075. doi: 10.1126/science.aaa1934.

10. Rachel C. Bandler, Ilaria Vitali, Ryan N. Delgado, May C. Ho, Elena Dvoretskova, Josue S. Ibarra Molinas, Paul W. Frazel, Maesoumeh Mohammadkhani, Robert Machold, Sophia Maedler, et al. Single-cell delineation of lineage and genetic identity in the mouse brain. Nature, 601(7893):404–409, 2022. ISSN 0028-0836. doi: 10.1038/s41586-021-04237-0.

11. Trygve E. Bakken, Nikolas L. Jorstad, Qiwen Hu, Blue B. Lake, Wei Tian, Brian E. Kalmbach, Megan Crow, Rebecca D. Hodge, Fenna M. Krienen, Staci A. Sorensen, et al. Comparative cellular analysis of motor cortex in human, marmoset and mouse. Nature, 598(7879):111–119, 2021. ISSN 0028-0836. doi: 10.1038/s41586-021-03465-8.

12. Samuel Morabito, Emily Miyoshi, Neethu Michael, Saba Shahin, Alessandra Cadete Martini, Elizabeth Head, Justine Silva, Kelsey Leavy, Mari Perez-Rosendahl, and Vivek Swarup. Single-nucleus chromatin accessibility and transcriptomic characterization of Alzheimer’s disease. Nature Genetics, 53(8):1143–1155, 2021. ISSN 1061-4036. doi: 10.1038/s41588-021-00894-z.

13. Hansruedi Mathys, Jose Davila-Velderrain, Zhuyu Peng, Fan Gao, Shahin Mohammadi, Jennie Z. Young, Madhvi Menon, Liang He, Fatema Abdurrob, Xueqiao Jiang, et al. Singlecell transcriptomic analysis of Alzheimer’s disease. Nature, 570(7761):332–337, 2019. ISSN 0028-0836. doi: 10.1038/s41586-019-1195-2.

14. Yingyue Zhou, Wilbur M. Song, Prabhakar S. Andhey, Amanda Swain, Tyler Levy, Kelly R. Miller, Pietro L. Poliani, Manuela Cominelli, Shikha Grover, Susan Gilfillan, et al. Human and mouse single-nucleus transcriptomics reveal TREM2-dependent and TREM2-independent cellular responses in Alzheimer’s disease. Nature Medicine, 26(1):131–142, 2020. ISSN 1078-8956. doi: 10.1038/s41591-019-0695-9.

15. Shun-Fat Lau, Han Cao, Amy K. Y. Fu, and Nancy Y. Ip. Single-nucleus transcriptome analysis reveals dysregulation of angiogenic endothelial cells and neuroprotective glia in Alzheimer’s disease. Proceedings of the National Academy of Sciences, 117(41):25800–25809, 2020. ISSN 0027-8424. doi: 10.1073/pnas.2008762117.

16. Kun Leng, Emmy Li, Rana Eser, Antonia Piergies, Rene Sit, Michelle Tan, Norma Neff, Song Hua Li, Roberta Diehl Rodriguez, Claudia Kimie Suemoto, et al. Molecular characterization of selectively vulnerable neurons in Alzheimer’s disease. Nature Neuroscience, 24 (2):276–287, 2021. ISSN 1097-6256. doi: 10.1038/s41593-020-00764-7.

17. Marcos Otero-Garcia, Sameehan U. Mahajani, Debia Wakhloo, Weijing Tang, Yue-Qiang Xue, Samuel Morabito, Jie Pan, Jane Oberhauser, Angela E. Madira, Tamara Shakouri, et al. Molecular signatures underlying neurofibrillary tangle susceptibility in Alzheimer’s disease. Neuron, 2022. ISSN 0896-6273. doi: 10.1016/j.neuron.2022.06.021.

18. Lambda Moses and Lior Pachter. Museum of spatial transcriptomics. Nature Methods, pages 1–13, 2022. ISSN 1548-7091. doi: 10.1038/s41592-022-01409-2.

19. Samuel G. Rodriques, Robert R. Stickels, Aleksandrina Goeva, Carly A. Martin, Evan Murray, Charles R. Vanderburg, Joshua Welch, Linlin M. Chen, Fei Chen, and Evan Z. Macosko. Slide-seq: A scalable technology for measuring genome-wide expression at high spatial resolution. Science, 363(6434):1463–1467, 2019. ISSN 0036-8075. doi: 10.1126/science.aaw1219.

20. Sanjay R. Srivatsan, Mary C. Regier, Eliza Barkan, Jennifer M. Franks, Jonathan S. Packer, Parker Grosjean, Madeleine Duran, Sarah Saxton, Jon J Ladd, Malte Spielmann, et al. Embryo-scale, single-cell spatial transcriptomics. Science, 373(6550):111–117, 2021. ISSN 0036-8075. doi: 10.1126/science.abb9536.

21. Patrik L. Ståhl, Fredrik Salmén, Sanja Vickovic, Anna Lundmark, José Fernández Navarro, Jens Magnusson, Stefania Giacomello, Michaela Asp, Jakub O. Westholm, Mikael Huss, et al. Visualization and analysis of gene expression in tissue sections by spatial transcriptomics. Science, 353(6294):78–82, 2016. ISSN 0036-8075. doi: 10.1126/science.aaf2403.

22. Xiao Wang, William E. Allen, Matthew A. Wright, Emily L. Sylwestrak, Nikolay Samusik, Sam Vesuna, Kathryn Evans, Cindy Liu, Charu Ramakrishnan, Jia Liu, et al. Three-dimensional intact-tissue sequencing of single-cell transcriptional states. Science, 361(6400), 2018. ISSN 0036-8075. doi: 10.1126/science.aat5691.

23. Rongqin Ke, Marco Mignardi, Alexandra Pacureanu, Jessica Svedlund, Johan Botling, Carolina Wählby, and Mats Nilsson. In situ sequencing for RNA analysis in preserved tissue and cells. Nature Methods, 10(9):857–860, 2013. ISSN 1548-7091. doi: 10.1038/nmeth.2563.

24. M. McCarron, P. McCallion, E. Reilly, P. Dunne, R. Carroll, and N. Mulryan. A prospective 20-year longitudinal follow-up of dementia in persons with Down syndrome. Journal of Intellectual Disability Research, 61(9):843–852, 2017. ISSN 0964-2633. doi: 10.1111/jir.12390.

25. Carter R. Palmer, Christine S. Liu, William J. Romanow, Ming-Hsiang Lee, and Jerold Chun. Altered cell and RNA isoform diversity in aging Down syndrome brains. Proceedings of the National Academy of Sciences, 118(47):e2114326118, 2021. ISSN 0027-8424. doi: 10.1073/pnas.2114326118.

26. Alessandra C. Martini, Thomas J. Gross, Elizabeth Head, and Mark Mapstone. Beyond amyloid: Immune, cerebrovascular, and metabolic contributions to Alzheimer disease in people with Down syndrome. Neuron, 2022. ISSN 0896-6273. doi: 10.1016/j.neuron.2022.04.001.

27. Juan Fortea, Eduard Vilaplana, Maria Carmona-Iragui, Bessy Benejam, Laura Videla, Isabel Barroeta, Susana Fernández, Miren Altuna, Jordi Pegueroles, Víctor Montal, et al. Clinical and biomarker changes of Alzheimer’s disease in adults with Down syndrome: a cross-sectional study. The Lancet, 395(10242):1988–1997, 2020. ISSN 0140-6736. doi: 10.1016/s0140-6736(20)30689-9.

28. Vivek Swarup, Flora I. Hinz, Jessica E. Rexach, Ken-ichi Noguchi, Hiroyoshi Toyoshiba, Akira Oda, Keisuke Hirai, Arjun Sarkar, Nicholas T. Seyfried, Chialin Cheng, et al. Identification of evolutionarily conserved gene networks mediating neurodegenerative dementia. Nature Medicine, 25(1):152–164, 2019. ISSN 1078-8956. doi: 10.1038/s41591-018-0223-3.

29. Vivek Swarup, Timothy S. Chang, Duc M. Duong, Eric B. Dammer, Jingting Dai, James J. Lah, Erik C.B. Johnson, Nicholas T. Seyfried, Allan I. Levey, and Daniel H. Geschwind. Identification of Conserved Proteomic Networks in Neurodegenerative Dementia. Cell Reports, 31(12):107807, 2020. ISSN 2211-1247. doi: 10.1016/j.celrep.2020.107807.

30. V. A. Traag, L. Waltman, and N. J. van Eck. From Louvain to Leiden: guaranteeing well-connected communities. Scientific Reports, 9(1):5233, 2019. doi: 10.1038/s41598-019-41695-z.

31. Edward Zhao, Matthew R. Stone, Xing Ren, Jamie Guenthoer, Kimberly S. Smythe, Thomas Pulliam, Stephen R. Williams, Cedric R. Uytingco, Sarah E. B. Taylor, Paul Nghiem, et al. Spatial transcriptomics at subspot resolution with BayesSpace. Nature Biotechnology, pages 1–10, 2021. ISSN 1087-0156. doi: 10.1038/s41587-021-00935-2.

32. Saroshi Minoshima, Bruno Giordani, Stanley Berent, Kirk A. Frey, Norman L. Foster, and David E. Kuhl. Metabolic reduction in the posterior cingulate cortex in very early Alzheimer’s disease. Annals of Neurology, 42(1):85–94, 1997. ISSN 0364-5134. doi: 10.1002/ana.410420114.

33. Chenling Xu, Romain Lopez, Edouard Mehlman, Jeffrey Regier, Michael I Jordan, and Nir Yosef. Probabilistic harmonization and annotation of single-cell transcriptomics data with deep generative models. Molecular Systems Biology, 17(1):e9620, 2021. ISSN 1744-4292. doi: 10.15252/msb.20209620.

34. Malte D. Luecken, M. Büttner, K. Chaichoompu, A. Danese, M. Interlandi, M. F. Mueller, D. C. Strobl, L. Zappia, M. Dugas, M. Colomé-Tatché, and Fabian J. Theis. Benchmarking atlas-level data integration in single-cell genomics. Nature Methods, 19(1):41–50, 2022. ISSN 1548-7091. doi: 10.1038/s41592-021-01336-8.

35. Andrew C. Yang, Ryan T. Vest, Fabian Kern, Davis P. Lee, Maayan Agam, Christina A. Maat, Patricia M. Losada, Michelle B. Chen, Nicholas Schaum, Nathalie Khoury, et al. A human brain vascular atlas reveals diverse mediators of Alzheimer’s risk. Nature, 603(7903):885–892, 2022. ISSN 0028-0836. doi: 10.1038/s41586-021-04369-3.

36. Samuel E. Marsh, Alec J. Walker, Tushar Kamath, Lasse Dissing-Olesen, Timothy R. Hammond, T. Yvanka de Soysa, Adam M. H. Young, Sarah Murphy, Abdulraouf Abdulraouf, Naeem Nadaf, et al. Dissection of artifactual and confounding glial signatures by single-cell sequencing of mouse and human brain. Nature Neuroscience, 25(3):306–316, 2022. ISSN 1097-6256. doi: 10.1038/s41593-022-01022-8.

37. Massimo Andreatta and Santiago J. Carmona. UCell: Robust and scalable single-cell gene signature scoring. Computational and Structural Biotechnology Journal, 19:3796–3798, 2021. ISSN 2001-0370. doi: 10.1016/j.csbj.2021.06.043.

38. H. Braak and E. Braak. Neuropathological stageing of Alzheimer-related changes. Acta Neuropathologica, 82(4):239–259, 1991. ISSN 0001-6322. doi: 10.1007/bf00308809.

39. Patrick R. Hof, John H. Morrison, and Kevin Cox. Quantitative analysis of a vulnerable subset of pyramidal neurons in Alzheimer’s disease: I. Superior frontal and inferior temporal cortex. Journal of Comparative Neurology, 301(1):44–54, 1990. ISSN 0021-9967. doi: 10.1002/cne.903010105.

40. Nanet Willumsen, Teresa Poole, Jennifer M. Nicholas, Nick C. Fox, Natalie S. Ryan, and Tammaryn Lashley. Variability in the type and layer distribution of cortical Aβ pathology in familial Alzheimer’s disease. Brain Pathology, 32(3):e13009, 2022. ISSN 1015-6305. doi: 10.1111/bpa.13009.

41. Bryn Farnsworth, Christiane Peuckert, Bettina Zimmermann, Elena Jazin, Petronella Kettunen, and Lina Sors Emilsson. Gene Expression of Quaking in Sporadic Alzheimer’s Disease Patients is Both Upregulated and Related to Expression Levels of Genes Involved in Amyloid Plaque and Neurofibrillary Tangle Formation. Journal of Alzheimer’s Disease, 53 (1):209–219, 2016. ISSN 1387-2877. doi: 10.3233/jad-160160.

42. Hongkui Zeng, Elaine H. Shen, John G. Hohmann, Seung Wook Oh, Amy Bernard, Joshua J. Royall, Katie J. Glattfelder, Susan M. Sunkin, John A. Morris, Angela L. Guillozet-Bongaarts, et al. Large-Scale Cellular-Resolution Gene Profiling in Human Neocortex Reveals Species-Specific Molecular Signatures. Cell, 149(2):483–496, 2012. ISSN 0092-8674. doi: 10.1016/j.cell.2012.02.052.

43. Hadas Keren-Shaul, Amit Spinrad, Assaf Weiner, Orit Matcovitch-Natan, Raz Dvir- Szternfeld, Tyler K. Ulland, Eyal David, Kuti Baruch, David Lara-Astaiso, Beata Toth, et al. A Unique Microglia Type Associated with Restricting Development of Alzheimer’s Disease. Cell, 169(7):1276–1290.e17, 2017. ISSN 0092-8674. doi: 10.1016/j.cell.2017.05.018.

44. Naomi Habib, Cristin McCabe, Sedi Medina, Miriam Varshavsky, Daniel Kitsberg, Raz Dvir- Szternfeld, Gilad Green, Danielle Dionne, Lan Nguyen, Jamie L. Marshall, et al. Diseaseassociated astrocytes in Alzheimer’s disease and aging. Nature Neuroscience, 23(6):701–706, 2020. ISSN 1097-6256. doi: 10.1038/s41593-020-0624-8.

45. Mor Kenigsbuch, Pierre Bost, Shahar Halevi, Yuzhou Chang, Shuo Chen, Qin Ma, Renana Hajbi, Benno Schwikowski, Bernd Bodenmiller, Hongjun Fu, et al. A shared diseaseassociated oligodendrocyte signature among multiple CNS pathologies. Nature Neuroscience, 25(7):876–886, 2022. ISSN 1097-6256. doi: 10.1038/s41593-022-01104-7.

46. Samuel Morabito, Fairlie Reese, Negin Rahimzadeh, Emily Miyoshi, and Vivek Swarup. hdWGCNA identifies co-expression networks in high-dimensional transcriptomics data. Cell Reports Methods, 6 2023. doi: 10.1016/j.crmeth.2023.100498.

47. Christopher L. Hartl, Gokul Ramaswami, William G. Pembroke, Sandrine Muller, Greta Pintacuda, Ashis Saha, Princy Parsana, Alexis Battle, Kasper Lage, and Daniel H. Geschwind. Coexpression network architecture reveals the brain-wide and multiregional basis of disease susceptibility. Nature Neuroscience, pages 1–11, 2021. ISSN 1097-6256. doi: 10.1038/s41593-021-00887-5.

48. Peter Langfelder, Rui Luo, Michael C. Oldham, and Steve Horvath. Is My Network Module Preserved and Reproducible? PLoS Computational Biology, 7(1):e1001057, 2011. ISSN 1553-734X. doi: 10.1371/journal.pcbi.1001057.

49. Logan Dumitrescu, Lisa L Barnes, Madhav Thambisetty, Gary Beecham, Brian Kunkle, William S Bush, Katherine A Gifford, Lori B Chibnik, Shubhabrata Mukherjee, Philip L De Jager, et al. Sex differences in the genetic predictors of Alzheimer’s pathology. Brain, 142 (9):2581–2589, 2019. ISSN 0006-8950. doi: 10.1093/brain/awz206.

50. Jaclyn M. Eissman, Logan Dumitrescu, Emily R. Mahoney, Alexandra N. Smith, Shubhabrata Mukherjee, Michael L. Lee, Phoebe Scollard, Seo-Eun Choi, William S. Bush, Corinne D. Engelman, et al. Sex differences in the genetic architecture of cognitive resilience to Alzheimer’s disease. Brain, 145(7):awac177–, 2022. ISSN 0006-8950. doi: 10.1093/brain/awac177.

51. Karen Irvine, Keith R. Laws, Tim M. Gale, and Tejinder K. Kondel. Greater cognitive deterioration in women than men with Alzheimer’s disease: A meta analysis. Journal of Clinical and Experimental Neuropsychology, 34(9):989–998, 2012. ISSN 1380-3395. doi: 10.1080/13803395.2012.712676.

52. Shawn D. Gale, Leslie Baxter, and Juliann Thompson. Greater memory impairment in dementing females than males relative to sex-matched healthy controls. Journal of Clinical and Experimental Neuropsychology, 38(5):527–533, 2016. ISSN 1380-3395. doi: 10.1080/13803395.2015.1132298.

53. Paul Hollingworth, Marian L. Hamshere, Valentina Moskvina, Kimberley Dowzell, Pamela J. Moore, Catherine Foy, Nicola Archer, Aoibhinn Lynch, Simon Lovestone, Carol Brayne, et al. Four Components Describe Behavioral Symptoms in 1,120 Individuals with Late-Onset Alzheimer’s Disease. Journal of the American Geriatrics Society, 54(9):1348–1354, 2006. ISSN 0002-8614. doi: 10.1111/j.1532-5415.2006.00854.x.

54. Jong-Chan Park, Hanbyeol Lim, Min Soo Byun, Dahyun Yi, Gihwan Byeon, Gijung Jung, Yu Kyeong Kim, Dong Young Lee, Sun-Ho Han, and Inhee Mook-Jung. Sex differences in the progression of glucose metabolism dysfunction in Alzheimer’s disease. Experimental & Molecular Medicine, 55(5):1023–1032, 2023. ISSN 1226-3613. doi: 10.1038/s12276-023-00993-3.

55. Nolan Brown, Kholoud Alkhayer, Robert Clements, Naveen Singhal, Roger Gregory, Sausan Azzam, Shuo Li, Ernest Freeman, and Jennifer McDonough. Neuronal Hemoglobin Expression and Its Relevance to Multiple Sclerosis Neuropathology. Journal of Molecular Neuroscience, 59(1):1–17, 2016. ISSN 0895-8696. doi: 10.1007/s12031-015-0711-6.

56. Runmin Wei, Siyuan He, Shanshan Bai, Emi Sei, Min Hu, Alastair Thompson, Ken Chen, Savitri Krishnamurthy, and Nicholas E. Navin. Spatial charting of single-cell transcriptomes in tissues. Nature Biotechnology, pages 1–10, 2022. ISSN 1087-0156. doi: 10.1038/s41587-022-01233-1.

57. Goran Sedmak and Miloš Judaš. White Matter Interstitial Neurons in the Adult Human Brain: 3% of Cortical Neurons in Quest for Recognition. Cells, 10(1):190, 2021. doi: 10.3390/cells10010190.

58. Suoqin Jin, Christian F. Guerrero-Juarez, Lihua Zhang, Ivan Chang, Raul Ramos, Chen-Hsiang Kuan, Peggy Myung, Maksim V. Plikus, and Qing Nie. Inference and analysis of cell-cell communication using CellChat. Nature Communications, 12(1):1088, 2021. doi: 10.1038/s41467-021-21246-9.

59. Kiyohito Mizutani, Muneaki Miyata, Hajime Shiotani, Takeshi Kameyama, and Yoshimi Takai. Nectins and Nectin-like molecules in synapse formation and involvement in neurological diseases. Molecular and Cellular Neuroscience, 115:103653, 2021. ISSN 1044-7431. doi: 10.1016/j.mcn.2021.103653.

60. Kiyohito Mizutani, Muneaki Miyata, Hajime Shiotani, Takeshi Kameyama, and Yoshimi Takai. Nectin-2 in general and in the brain. Molecular and Cellular Biochemistry, 477(1):167–180, 2022. ISSN 0300-8177. doi: 10.1007/s11010-021-04241-y.

61. Johanna Tomorsky, Philip R. L. Parker, Chris Q. Doe, and Cristopher M. Niell. Precise levels of nectin-3 are required for proper synapse formation in postnatal visual cortex. Neural Development, 15(1):13, 2020. doi: 10.1186/s13064-020-00150-w.

62. Alireza Nazarian, Anatoliy I. Yashin, and Alexander M. Kulminski. Genome-wide analysis of genetic predisposition to Alzheimer’s disease and related sex disparities. Alzheimer’s Research & Therapy, 11(1):5, 2019. doi: 10.1186/s13195-018-0458-8.

63. Yanfa Sun, Jingjing Zhu, Dan Zhou, Saranya Canchi, Chong Wu, Nancy J. Cox, Robert A. Rissman, Eric R. Gamazon, and Lang Wu. A transcriptome-wide association study of Alzheimer’s disease using prediction models of relevant tissues identifies novel candidate susceptibility genes. Genome Medicine, 13(1):141, 2021. doi: 10.1186/s13073-021-00959-y.

64. Ananya Chakraborty, Alwin Kamermans, Bert van het Hof, Kitty Castricum, Ed Aanhane, Jack van Horssen, Victor L Thijssen, Philip Scheltens, Charlotte E Teunissen, Ruud D Fontijn, et al. Angiopoietin like-4 as a novel vascular mediator in capillary cerebral amyloid angiopathy. Brain, 141(12):3377–3388, 2018. ISSN 0006-8950. doi: 10.1093/brain/awy274.

65. Alzheimer Disease Genetics Consortium (ADGC), The European Alzheimer’s Disease Initiative (EADI), (CHARGE)„ Cohorts for Heart and Aging Research in Genomic Epidemiology Consortium, (GERAD/PERADES)„ Genetic and Environmental Risk in AD/Defining Genetic, Polygenic and Environmental Risk for Alzheimer’s Disease Consortium, Brian W Kunkle, Benjamin Grenier-Boley, Rebecca Sims, Joshua C Bis, Vincent Damotte, Adam C Naj, et al. Genetic meta-analysis of diagnosed Alzheimer’s disease identifies new risk loci and implicates Aβ, tau, immunity and lipid processing. Nature Genetics, 51(3):414–430, 2019. ISSN 1061-4036. doi: 10.1038/s41588-019-0358-2.

66. European Alzheimer’s Disease Initiative (EADI), (GERAD), Genetic and Environmental Risk in Alzheimer’s Disease, Alzheimer’s Disease Genetic Consortium (ADGC), (CHARGE), Cohorts for Heart and Aging Research in Genomic Epidemiology, Jean-Charles Lambert, Carla A Ibrahim-Verbaas, Denise Harold, Adam C Naj, Rebecca Sims, Céline Bellenguez, et al. Meta-analysis of 74,046 individuals identifies 11 new susceptibility loci for Alzheimer’s disease. Nature Genetics, 45(12):1452–1458, 2013. ISSN 1061-4036. doi: 10.1038/ng.2802.

67. Nisha Rathore, Sree Ranjani Ramani, Homer Pantua, Jian Payandeh, Tushar Bhangale, Arthur Wuster, Manav Kapoor, Yonglian Sun, Sharookh B. Kapadia, Lino Gonzalez, et al. Paired Immunoglobulin-like Type 2 Receptor Alpha G78R variant alters ligand binding and confers protection to Alzheimer’s disease. PLoS Genetics, 14(11):e1007427, 2018. ISSN 1553-7390. doi: 10.1371/journal.pgen.1007427.

68. Karin Lopatko Lindman, Caroline Jonsson, Bodil Weidung, Jan Olsson, Janardan P. Pandey, Dmitry Prokopenko, Rudolph E. Tanzi, Göran Hallmans, Sture Eriksson, Fredrik Elgh, and Hugo Lövheim. PILRA polymorphism modifies the effect of APOE4 and GM17 on Alzheimer’s disease risk. Scientific Reports, 12(1):13264, 2022. doi: 10.1038/s41598-022-17058-6.

69. Chun Chen, David McDonald, Alasdair Blain, Ashwin Sachdeva, Laura Bone, Anna L. M. Smith, Charlotte Warren, Sarah J. Pickett, Gavin Hudson, Andrew Filby, et al. Imaging mass cytometry reveals generalised deficiency in OXPHOS complexes in Parkinson’s disease. npj Parkinson’s Disease, 7(1):39, 2021. ISSN 2373-8057. doi: 10.1038/s41531-021-00182-x.

70. Wei-Ting Chen, Ashley Lu, Katleen Craessaerts, Benjamin Pavie, Carlo Sala Frigerio, Nikky Corthout, Xiaoyan Qian, Jana Laláková, Malte Kühnemund, Iryna Voytyuk, et al. Spatial Transcriptomics and In Situ Sequencing to Study Alzheimer’s Disease. Cell, 182(4):976–991.e19, 2020. ISSN 0092-8674. doi: 10.1016/j.cell.2020.06.038.

71. Arthur Getis and J. K. Ord. The Analysis of Spatial Association by Use of Distance Statistics. Geographical Analysis, 24(3):189–206, 1992. ISSN 0016-7363. doi: 10.1111/j.1538-4632.1992.tb00261.x.

72. J. K. Ord and Arthur Getis. Local Spatial Autocorrelation Statistics: Distributional Issues and an Application. Geographical Analysis, 27(4):286–306, 1995. ISSN 0016-7363. doi: 10.1111/j.1538-4632.1995.tb00912.x.

73. Nicholas T. Seyfried, Eric B. Dammer, Vivek Swarup, Divya Nandakumar, Duc M. Duong, Luming Yin, Qiudong Deng, Tram Nguyen, Chadwick M. Hales, Thomas Wingo, et al. A Multi-network Approach Identifies Protein-Specific Co-expression in Asymptomatic and Symptomatic Alzheimer’s Disease. Cell Systems, 4(1):60–72.e4, 2017. ISSN 2405-4712. doi: 10.1016/j.cels.2016.11.006.

74. Matthew J. Simon, Marie X. Wang, Charles F. Murchison, Natalie E. Roese, Erin L. Boespflug, Randall L. Woltjer, and Jeffrey J. Iliff. Transcriptional network analysis of human astrocytic endfoot genes reveals region-specific associations with dementia status and tau pathology. Scientific Reports, 8(1):12389, 2018. doi: 10.1038/s41598-018-30779-x.

75. Masahiro Shijo, Hiroyuki Honda, Satoshi O. Suzuki, Hideomi Hamasaki, Masaaki Hokama, Nona Abolhassani, Yusaku Nakabeppu, Toshiharu Ninomiya, Takanari Kitazono, and Toru Iwaki. Association of adipocyte enhancer-binding protein 1 with Alzheimer’s disease pathology in human hippocampi. Brain Pathology, 28(1):58–71, 2018. ISSN 1015-6305. doi: 10.1111/bpa.12475.

76. Daniel J. Panyard, Justin McKetney, Yuetiva K. Deming, Autumn R. Morrow, Gilda E. Ennis, Erin M. Jonaitis, Carol A. Van Hulle, Chengran Yang, Yun Ju Sung, Muhammad Ali, et al. Large-scale proteome and metabolome analysis of CSF implicates altered glucose and carbon metabolism and succinylcarnitine in Alzheimer’s disease. Alzheimer’s & Dementia, 2023. ISSN 1552-5260. doi: 10.1002/alz.13130.

77. Ashlyn G. Anderson, Brianne B. Rogers, Jacob M. Loupe, Ivan Rodriguez-Nunez, Sydney C. Roberts, Lauren M. White, J. Nicholas Brazell, William E. Bunney, Blynn G. Bunney, Stanley J. Watson, et al. Single nucleus multiomics identifies ZEB1 and MAFB as candidate regulators of Alzheimer’s disease-specific cis-regulatory elements. Cell Genomics, page 100263, 2023. ISSN 2666-979X. doi: 10.1016/j.xgen.2023.100263.

78. V. P. Prasher, S. G. Sajith, S. D. Rees, A. Patel, S. Tewari, N. Schupf, and W. B. Zigman. Significant effect of APOE epsilon 4 genotype on the risk of dementia in Alzheimer’s disease and mortality in persons with Down Syndrome. International Journal of Geriatric Psychiatry, 23(11):1134–1140, 2008. ISSN 0885-6230. doi: 10.1002/gps.2039.

79. Joseph H. Lee, Annie J. Lee, Lam-Ha Dang, Deborah Pang, Sergey Kisselev, Sharon J. Krinsky-McHale, Warren B. Zigman, José A. Luchsinger, Wayne Silverman, Benjamin Tycko, et al. Candidate gene analysis for Alzheimer’s disease in adults with Down syndrome. Neurobiology of Aging, 56:150–158, 2017. ISSN 0197-4580. doi: 10.1016/j.neurobiolaging.2017.04.018.

80. Nicola Thrupp, Carlo Sala Frigerio, Leen Wolfs, Nathan G. Skene, Nicola Fattorelli, Suresh Poovathingal, Yannick Fourne, Paul M. Matthews, Tom Theys, Renzo Mancuso, et al. Single-Nucleus RNA-Seq Is Not Suitable for Detection of Microglial Activation Genes in Humans. Cell Reports, 32(13):108189, 2020. ISSN 2211-1247. doi: 10.1016/j.celrep.2020.108189.

81. Stephen J. Fleming, John C. Marioni, and Mehrtash Babadi. CellBender removebackground: a deep generative model for unsupervised removal of background noise from scRNA-seq datasets. bioRxiv, page 791699, 2019. doi: 10.1101/791699.

82. Samuel L. Wolock, Romain Lopez, and Allon M. Klein. Scrublet: Computational Identification of Cell Doublets in Single-Cell Transcriptomic Data. Cell Systems, 8(4):281–291.e9, 2019. ISSN 2405-4712. doi: 10.1016/j.cels.2018.11.005.

83. Andrew Butler, Paul Hoffman, Peter Smibert, Efthymia Papalexi, and Rahul Satija. Integrating single-cell transcriptomic data across different conditions, technologies, and species. Nature Biotechnology, 36(5):411–420, 2018. ISSN 1087-0156. doi: 10.1038/nbt.4096.

84. Tim Stuart, Andrew Butler, Paul Hoffman, Christoph Hafemeister, Efthymia Papalexi, William M. Mauck, Yuhan Hao, Marlon Stoeckius, Peter Smibert, and Rahul Satija. Comprehensive Integration of Single-Cell Data. Cell, 177(7):1888–1902.e21, 2019. ISSN 0092-8674. doi: 10.1016/j.cell.2019.05.031.

85. Yuhan Hao, Stephanie Hao, Erica Andersen-Nissen, William M. Mauck, Shiwei Zheng, Andrew Butler, Maddie J. Lee, Aaron J. Wilk, Charlotte Darby, Michael Zager, et al. Integrated analysis of multimodal single-cell data. Cell, 184(13):3573–3587.e29, 2021. ISSN 0092-8674. doi: 10.1016/j.cell.2021.04.048.

86. F. Alexander Wolf, Philipp Angerer, and Fabian J. Theis. SCANPY: large-scale singlecell gene expression data analysis. Genome Biology, 19(1):15, 2018. doi: 10.1186/s13059-017-1382-0.

87. Romain Lopez, Jeffrey Regier, Michael B. Cole, Michael I. Jordan, and Nir Yosef. Deep generative modeling for single-cell transcriptomics. Nature Methods, 15(12):1053–1058, 2018. ISSN 1548-7091. doi: 10.1038/s41592-018-0229-2.

88. Leland McInnes, John Healy, and James Melville. UMAP: Uniform Manifold Approximation and Projection for Dimension Reduction. arXiv, 2018.

89. Páll Melsted, A. Sina Booeshaghi, Lauren Liu, Fan Gao, Lambda Lu, Kyung Hoi (Joseph) Min, Eduardo da Veiga Beltrame, Kristján Eldjárn Hjörleifsson, Jase Gehring, and Lior Pachter. Modular, efficient and constant-memory single-cell RNA-seq preprocessing. Nature Biotechnology, 39(7):813–818, 2021. ISSN 1087-0156. doi: 10.1038/s41587-021-00870-2.

90. Ilya Korsunsky, Nghia Millard, Jean Fan, Kamil Slowikowski, Fan Zhang, Kevin Wei, Yuriy Baglaenko, Michael Brenner, Po-ru Loh, and Soumya Raychaudhuri. Fast, sensitive and accurate integration of single-cell data with Harmony. Nature Methods, 16(12):1289–1296, 2019. ISSN 1548-7091. doi: 10.1038/s41592-019-0619-0.

91. Mohammad Lotfollahi, Mohsen Naghipourfar, Malte D. Luecken, Matin Khajavi, Maren Büttner, Marco Wagenstetter, Žiga Avsec, Adam Gayoso, Nir Yosef, Marta Interlandi, et al. Mapping single-cell data to reference atlases by transfer learning. Nature Biotechnology, 40(1):121–130, 2022. ISSN 1087-0156. doi: 10.1038/s41587-021-01001-7.

92. Greg Finak, Andrew McDavid, Masanao Yajima, Jingyuan Deng, Vivian Gersuk, Alex K. Shalek, Chloe K. Slichter, Hannah W. Miller, M. Juliana McElrath, Martin Prlic, et al. MAST: a flexible statistical framework for assessing transcriptional changes and characterizing heterogeneity in single-cell RNA sequencing data. Genome Biology, 16(1):278, 2015. doi: 10.1186/s13059-015-0844-5.

93. Edward Y Chen, Christopher M Tan, Yan Kou, Qiaonan Duan, Zichen Wang, Gabriela Vaz Meirelles, Neil R Clark, and Avi Ma’ayan. Enrichr: interactive and collaborative HTML5 gene list enrichment analysis tool. BMC Bioinformatics, 14(1):128, 2013. doi: 10.1186/1471-2105-14-128.

94. Peter Langfelder and Steve Horvath. WGCNA: an R package for weighted correlation network analysis. BMC Bioinformatics, 9(1):559, 2008. doi: 10.1186/1471-2105-9-559.

95. Peter Langfelder, Bin Zhang, and Steve Horvath. Defining clusters from a hierarchical cluster tree: the Dynamic Tree Cut package for R. Bioinformatics, 24(5):719–720, 2007. ISSN 1367-4803. doi: 10.1093/bioinformatics/btm563.

96. Jake R Conway, Alexander Lex, and Nils Gehlenborg. UpSetR: an R package for the visualization of intersecting sets and their properties. Bioinformatics, 33(18):2938–2940, 2017. ISSN 1367-4803. doi: 10.1093/bioinformatics/btx364.

97. Lambda Moses, Pétur Helgi Einarsson, Kayla Jackson, Laura Luebbert, A. Sina Booeshagh, Sindri Antonsson, Páll Melsted, and Lior Pacther. Voyager: exploratory single-cell genomics data analysis with geospatial statistics. bioRxiv, 2023. doi: 10.1101/2023.07.20.549945.

98. Junyue Cao, Malte Spielmann, Xiaojie Qiu, Xingfan Huang, Daniel M. Ibrahim, Andrew J. Hill, Fan Zhang, Stefan Mundlos, Lena Christiansen, Frank J. Steemers, et al. The singlecell transcriptional landscape of mammalian organogenesis. Nature, 566(7745):496–502, 2019. ISSN 0028-0836. doi: 10.1038/s41586-019-0969-x.

