## Supplementary Figures for "Spatial and single-nucleus transcriptomic analysis of genetic and sporadic forms of Alzheimer’s Disease"

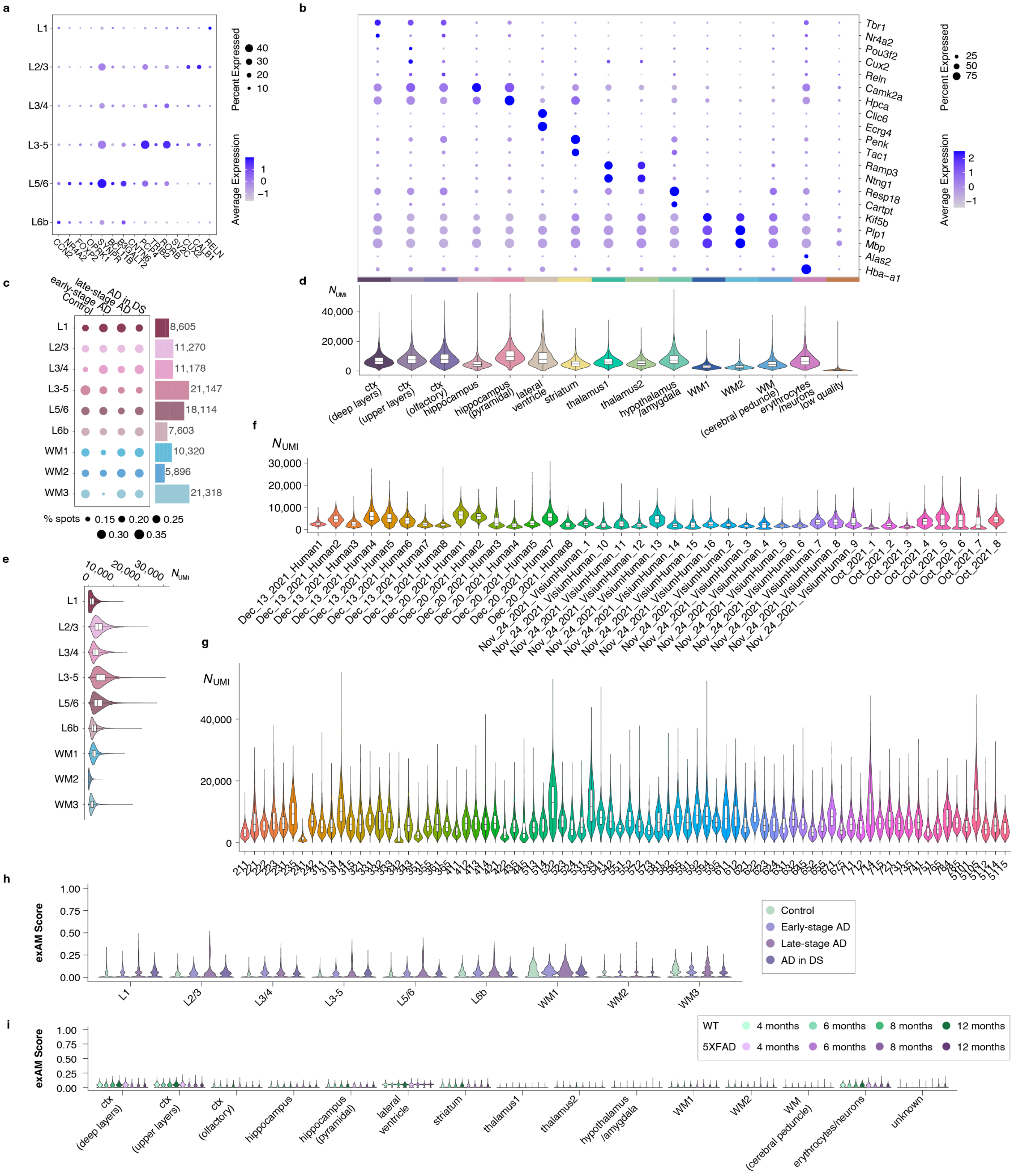


**Fig. S1. Quality control of the human and mouse spatial transcriptomics datasets.** **a**, Dot plot showing the expression of selected marker genes in the cortical layer clusters from the human ST dataset. **b**, Dot plot showing the expression of selected marker genes in the clusters from the mouse ST dataset. **c**, The number of spots profiled in each cluster, and the proportion of diagnosis attributed to each of the clusters. **d-e**, Distributions of the number of UMI captured in each of the mouse (**d**) and human (**e**) ST clusters. **f-g**, Distributions of the number of UMI captured in each of the human (**f**) and mouse (**g**) samples. **h-i**, Distributions of ex-vivo activation[1] UCell[2] scores in each of the ST clusters in the human (**h**) and mouse (**i** clusters). For box and whisker plots, box boundaries and line correspond to the interquartile range (IQR) and median, respectively. Whiskers extend to the lowest or highest data points that are no further than 1.5 times the IQR from the box boundaries.

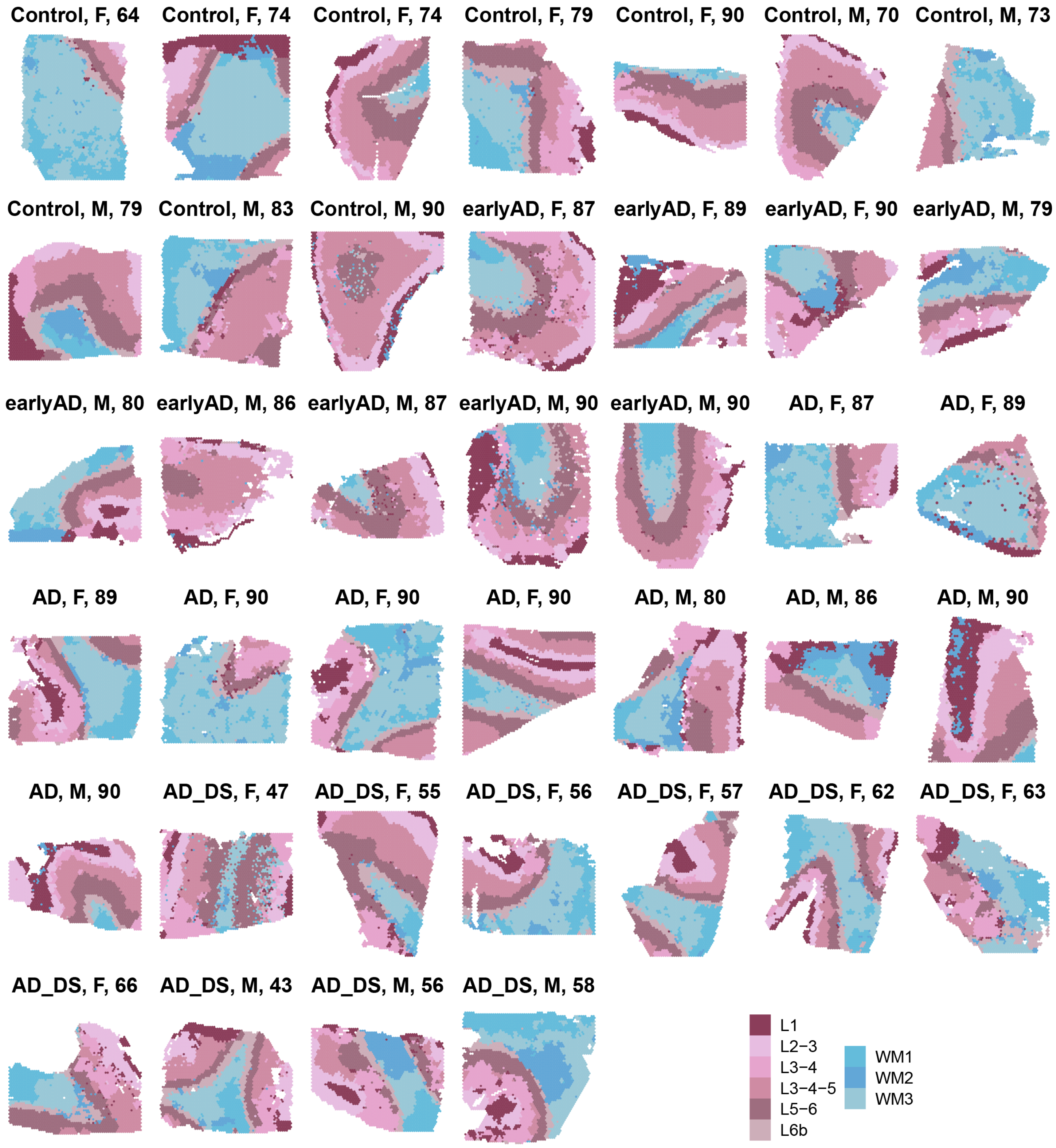


**Fig. S2. Clustering results in the human ST dataset.** We used BayesSpace[3] to jointly cluster spatial transcriptomic spots in the 39 samples from the human dataset based on their transcriptomic content as well as their spatial information (Methods). This process resulted in nine clusters, which we annotated based on marker gene expression and anatomic location. Each human ST sample is shown here with each spot colored by BayesSpace cluster assignment. We organized the samples in this plot by disease status, sex, and age.

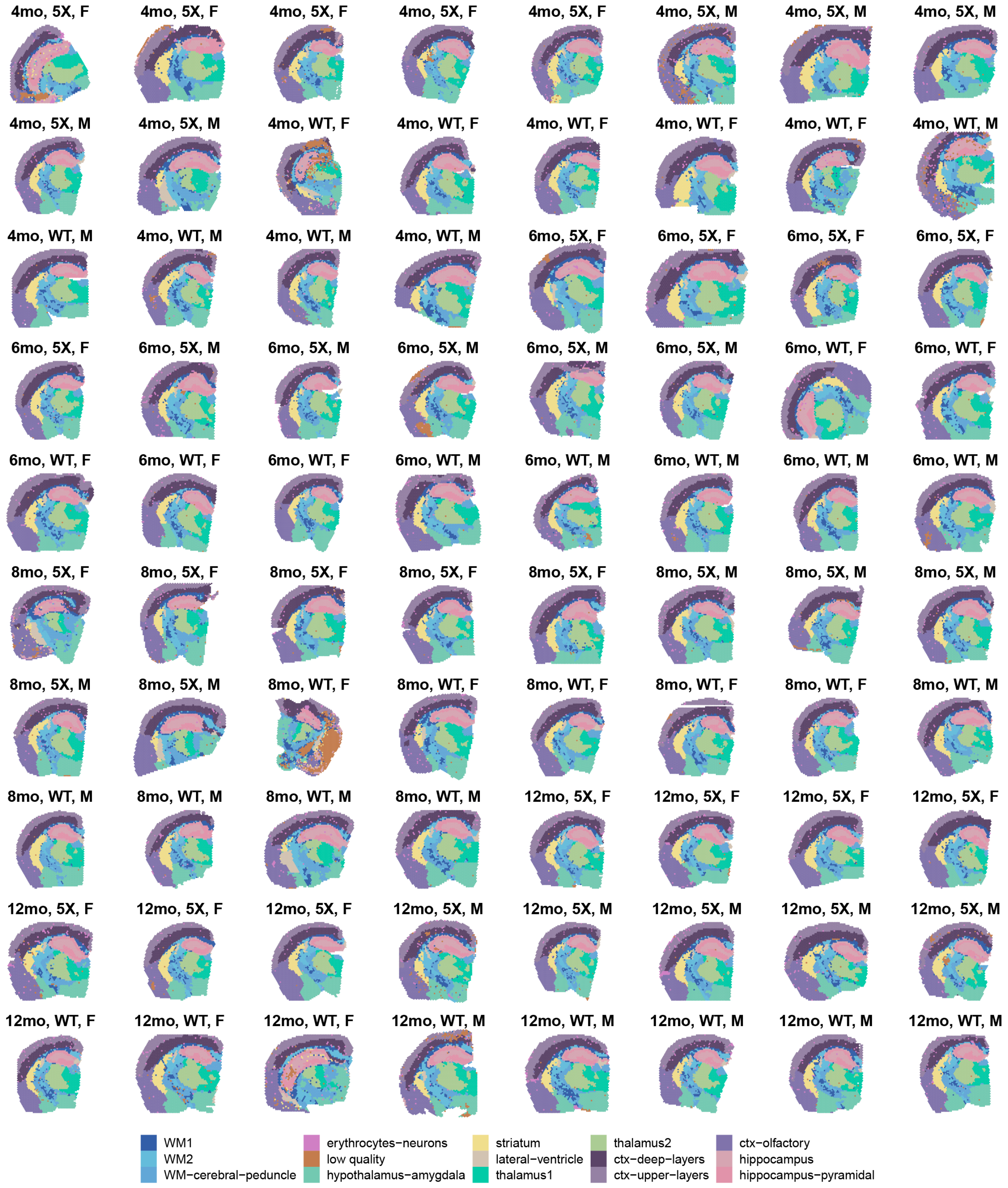


**Fig. S3. Clustering results in the mouse ST dataset.** We used BayesSpace[3] to jointly cluster spatial transcriptomic spots in the 80 samples from the mouse dataset based on their transcriptomic content as well as their spatial information (Methods). This process resulted in 15 clusters, which we annotated based on marker gene expression and anatomic location. Each mouse ST sample is shown here with each spot colored by BayesSpace cluster assignment. We organized the samples in this plot by disease age, genotype, and sex.

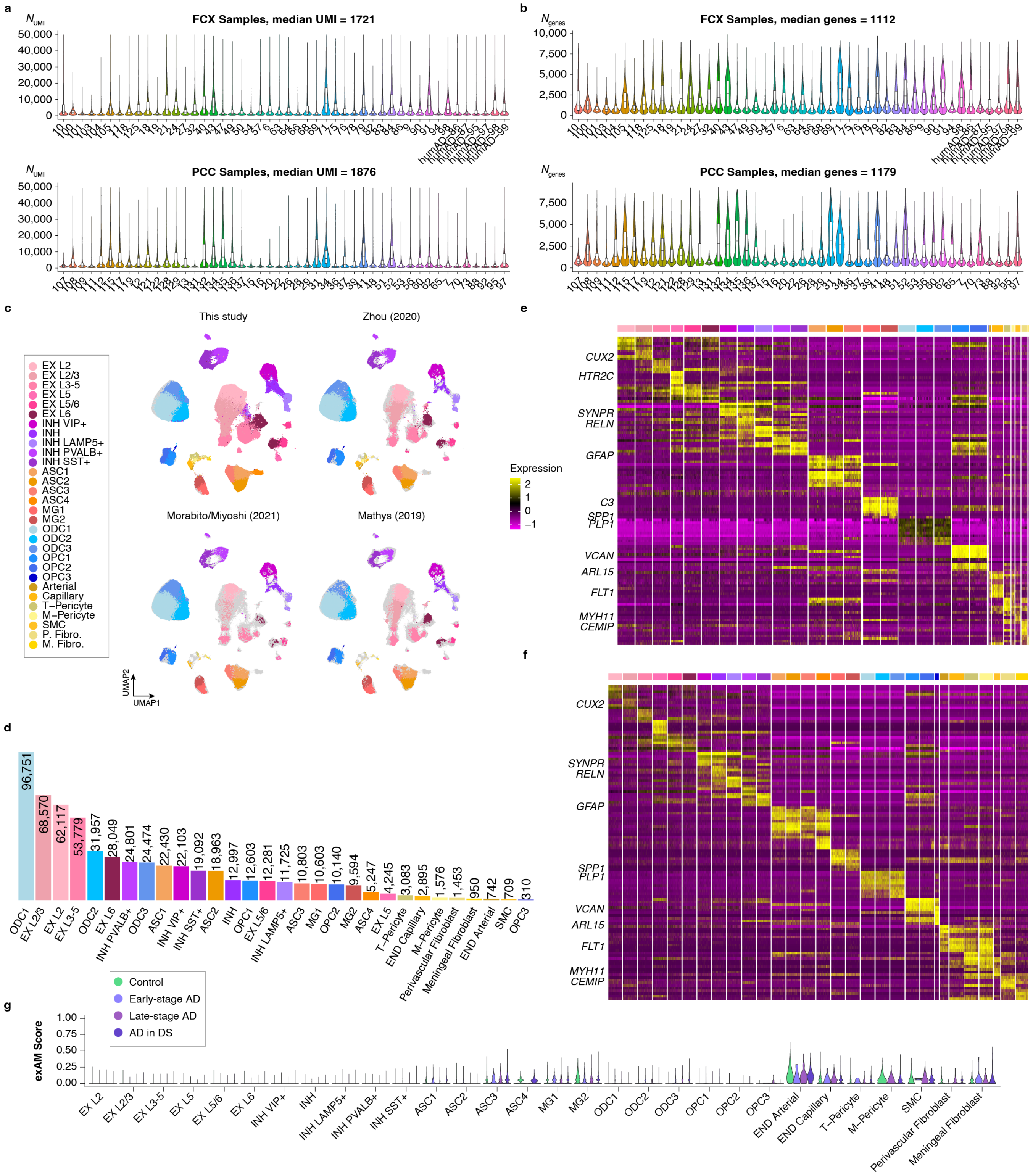


**Fig. S4. Quality control of the integrated snRNA-seq dataset. a**, Distributions of the number of UMI captured in each snRNA-seq sample for the frontal cortex (FCX, top) and the posterior cingulate cortex (PCC, bottom). **b**, Distributions of the number of genes captured in each snRNA-seq sample for the FCX (top) and the PCC (bottom). **c**, UMAP dimensionality reduction plot of the integrated snRNA-seq dataset comprised of the data generated in this study and three previous snRNA-seq datasets of AD. The UMAP plot is faceted by study of origin, and colored by cluster annotations. **d**, Bar plot showing the number of nuclei in each of the snRNA-seq clusters. **e**, Heatmap showing the expression of the top 5 marker genes ranked by effect size in the snRNA-seq dataset. These marker genes were identified using only the data generated in this study, and this heatmap only shows expression data from these nuclei (Methods). **f**, Heatmap showing the expression of the same genes as in panel (**e**) in the three other snRNA-seq datasets. **g**, Distributions of ex-vivo activation [1] UCell [2] scores in each of the snRNA-seq clusters. For box and whisker plots, box boundaries and line correspond to the interquartile range (IQR) and median, respectively. Whiskers extend to the lowest or highest data points that are no further than 1.5 times the IQR from the box boundaries.

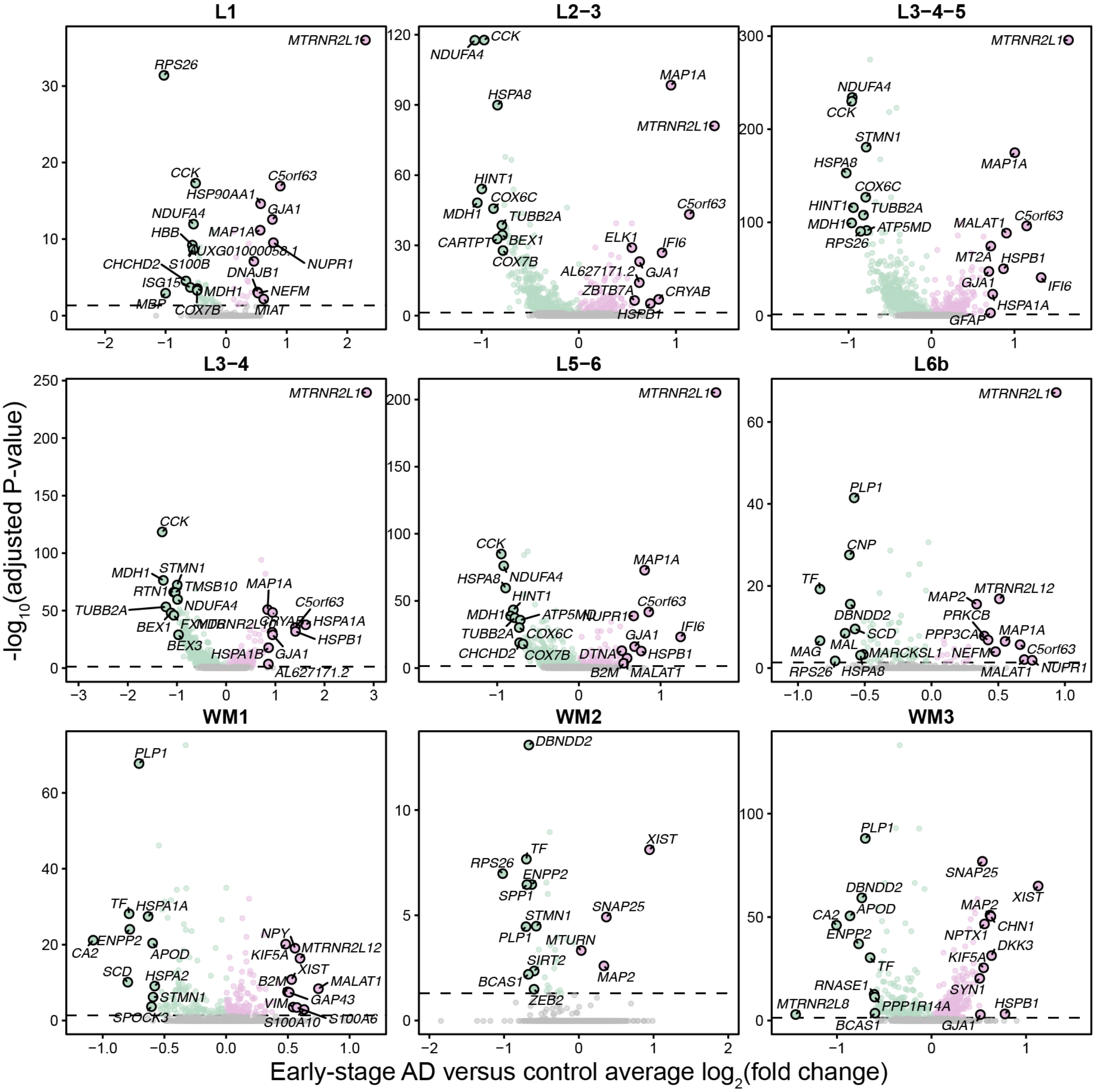


**Fig. S5. Early-stage AD versus control DEGs in the ST dataset.** Volcano plots show the adjusted significance levels and effect sizes from the differential gene expression comparisons between early-stage AD cases versus cognitively normal controls in the human ST dataset. The results are shown for the differential gene expression analysis in each of the nine spatial clusters. The top and bottom ten genes by effect size are annotated.

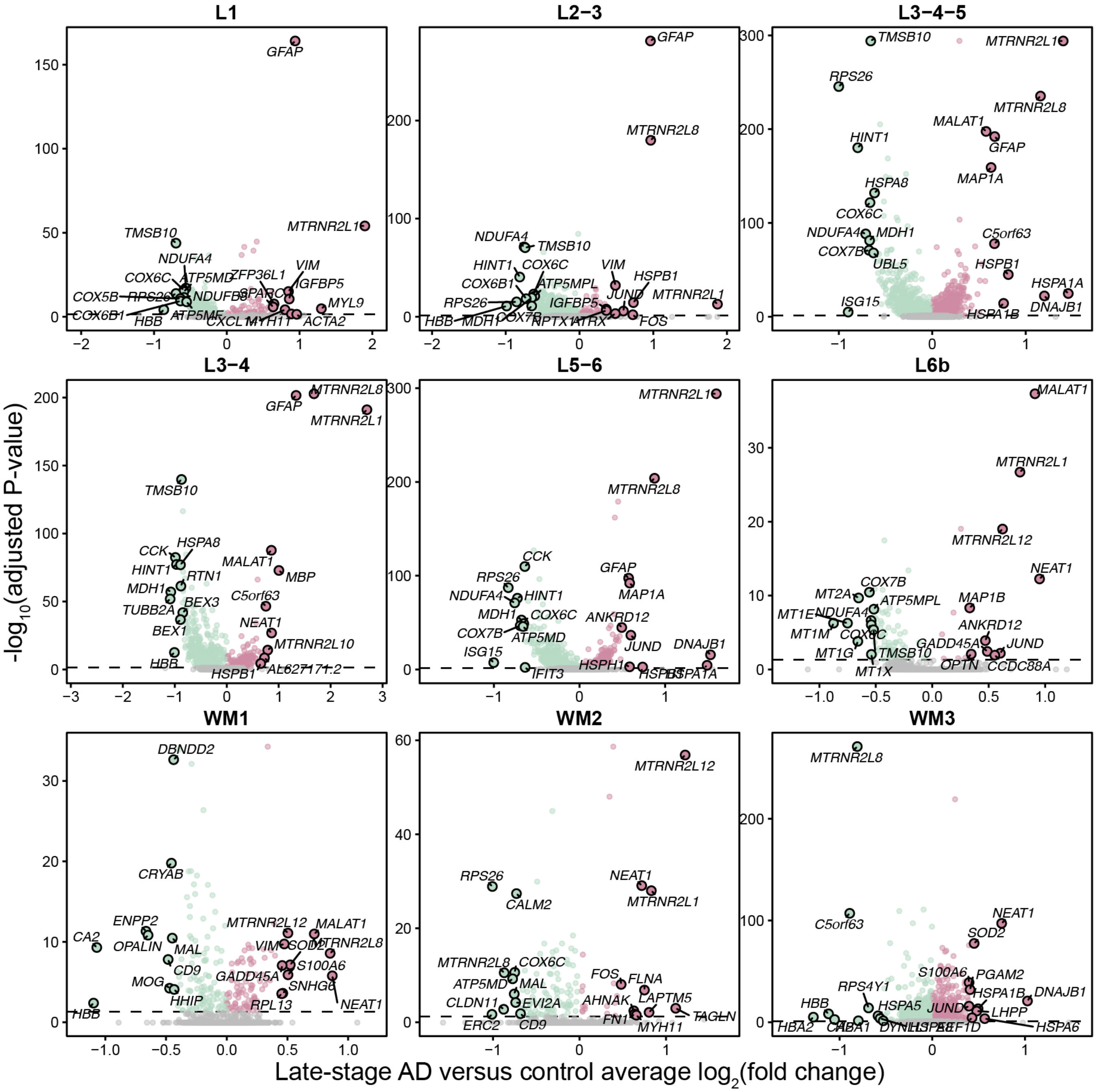


**Fig. S6. Late-stage AD versus control DEGs in the ST dataset.** Volcano plots show the adjusted significance levels and effect sizes from the differential gene expression comparisons between late-stage AD cases versus cognitively normal controls in the human ST dataset. The results are shown for the differential gene expression analysis in each of the nine spatial clusters. The top and bottom ten genes by effect size are annotated.

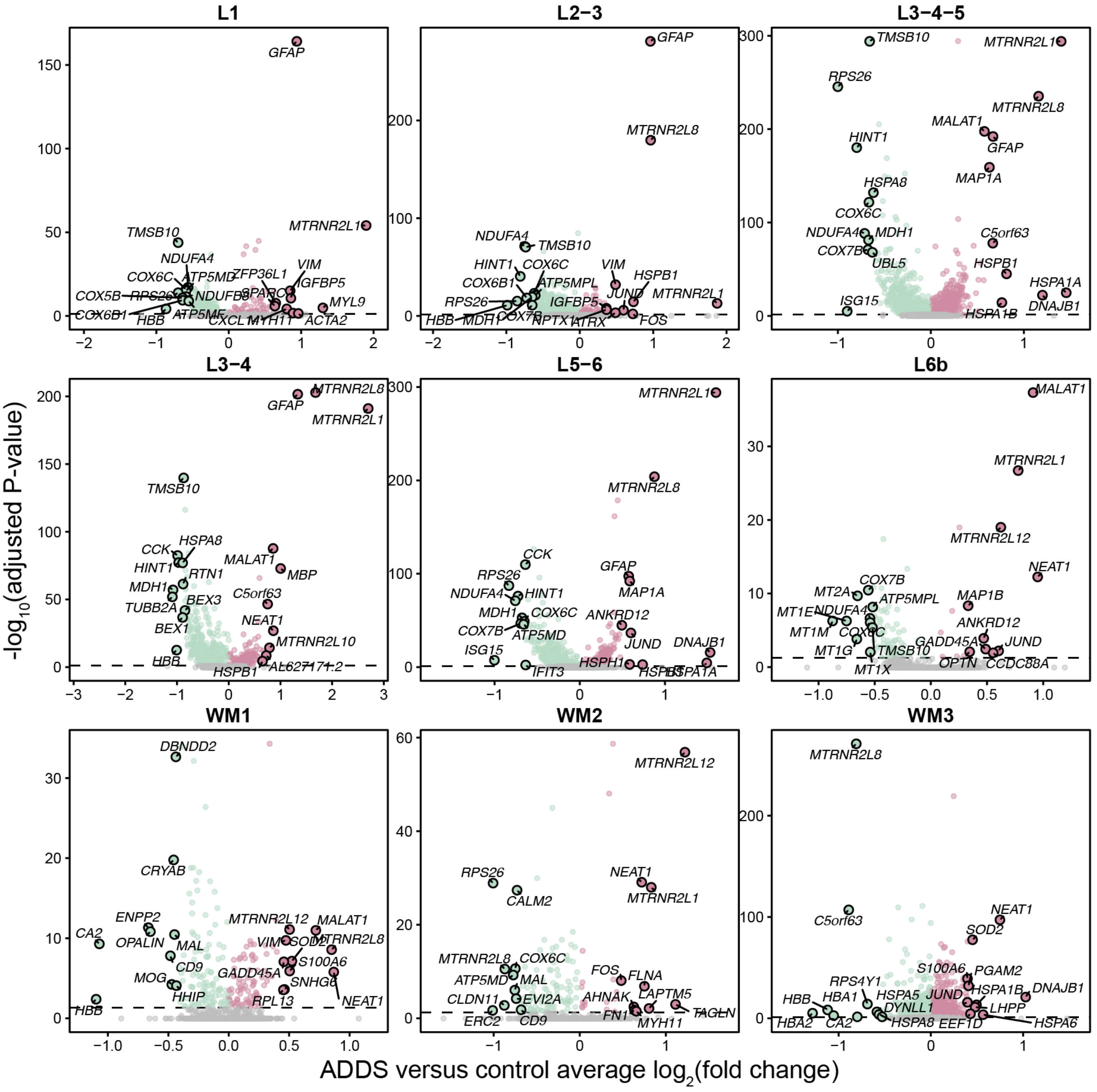


**Fig. S7. AD in DS versus control DEGs in the ST dataset.** Volcano plots show the adjusted significance levels and effect sizes from the differential gene expression comparisons between AD in DS cases versus cognitively normal controls in the human ST dataset. The results are shown for the differential gene expression analysis in each of the nine spatial clusters. The top and bottom ten genes by effect size are annotated.

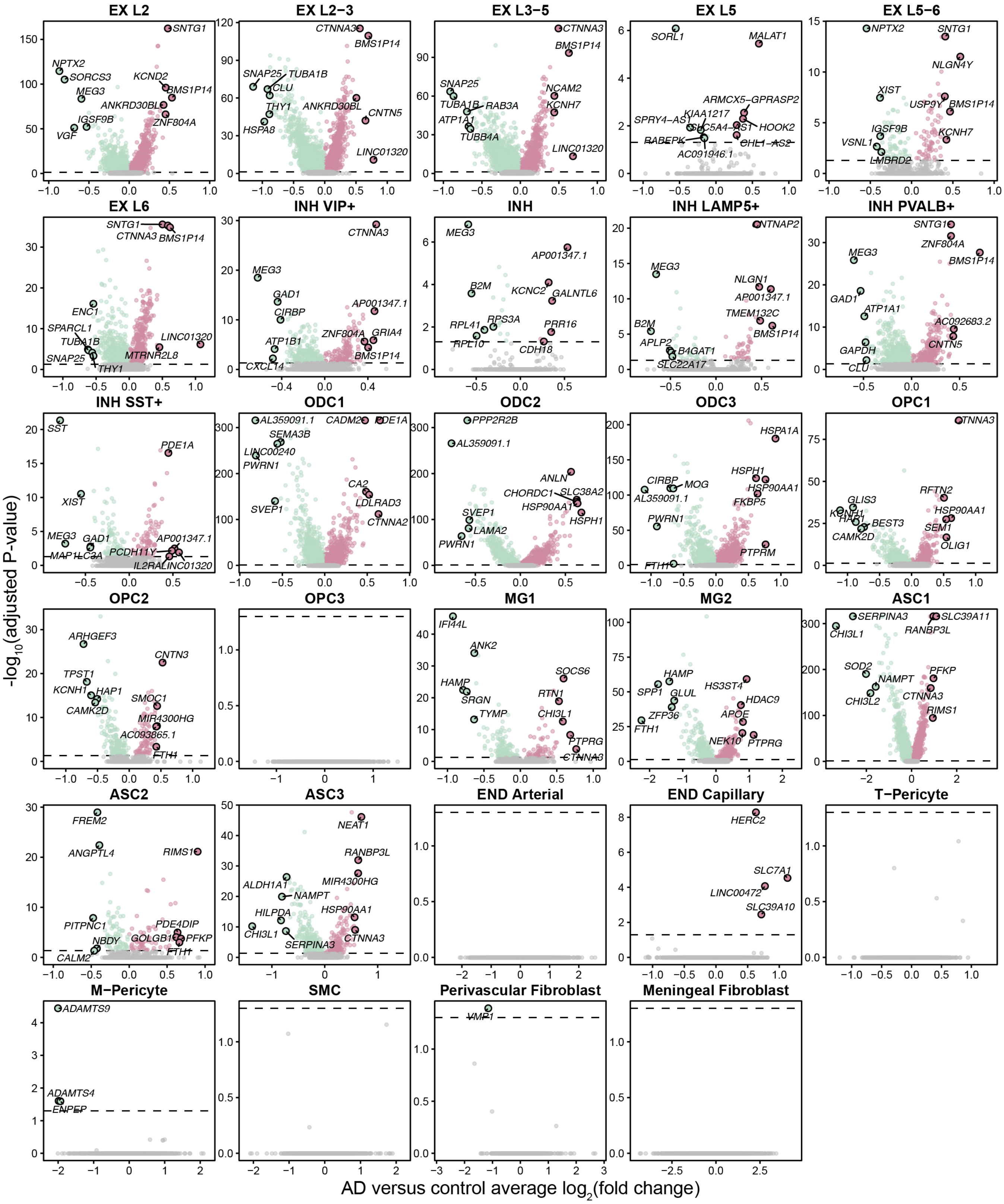


**Fig. S8. Late-stage AD versus control DEGs in the snRNA-seq dataset.** Volcano plots show the adjusted significance levels and effect sizes from the differential gene expression comparisons between late-stage AD cases versus cognitively normal controls from the integrated analysis of the three previously published snRNA-seq datasets. The results are shown for the differential gene expression analysis in each of the snRNA-seq clusters. The top and bottom five genes by effect size are annotated.

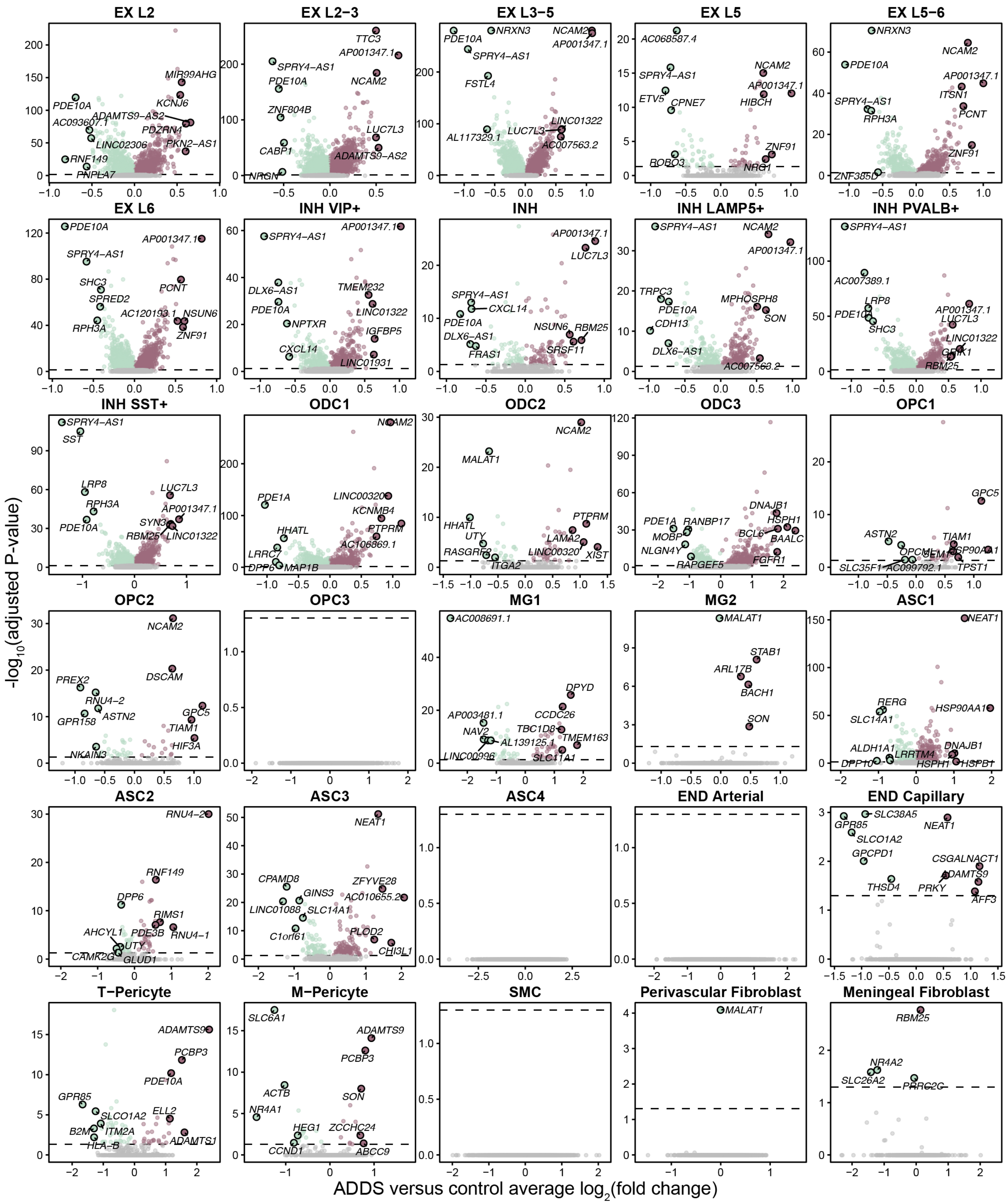


**Fig. S9. AD in DS versus control DEGs in the FCX snRNA-seq dataset.** Volcano plots show the adjusted significance levels and effect sizes from the differential gene expression comparisons between AD in DS cases versus cognitively normal controls in the frontal cortex (FCX) snRNA-seq dataset. The results are shown for the differential gene expression analysis in each of the snRNA-seq clusters. The top and bottom five genes by effect size are annotated.

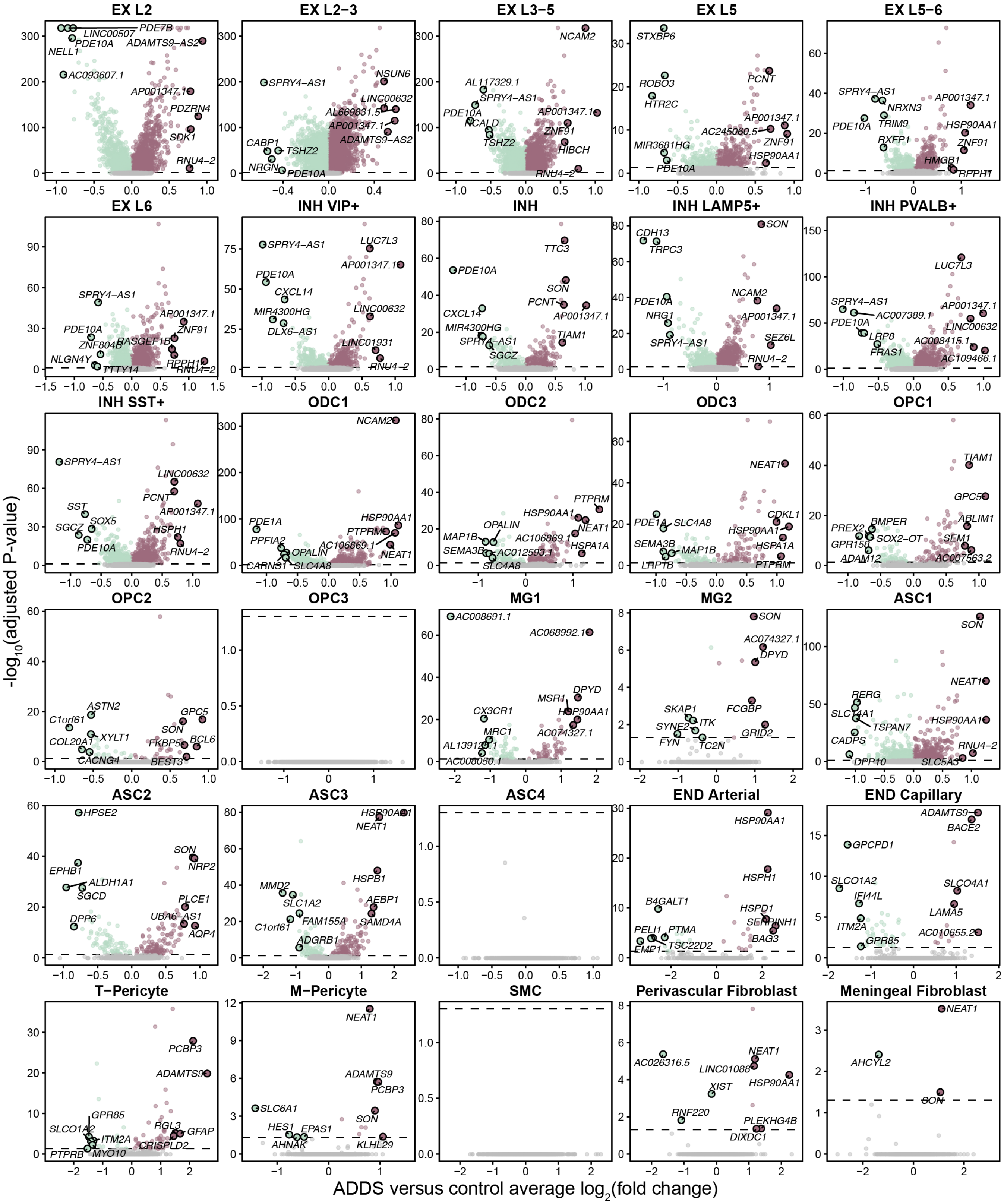


**Fig. S10. AD in DS versus control DEGs in the PCC snRNA-seq dataset** Volcano plots show the adjusted significance levels and effect sizes from the differential gene expression comparisons between AD in DS cases versus cognitively normal controls in the posterior cingulate cortex (PCC) snRNA-seq dataset. The results are shown for the differential gene expression analysis in each of the snRNA-seq clusters. The top and bottom five genes by effect size are annotated.

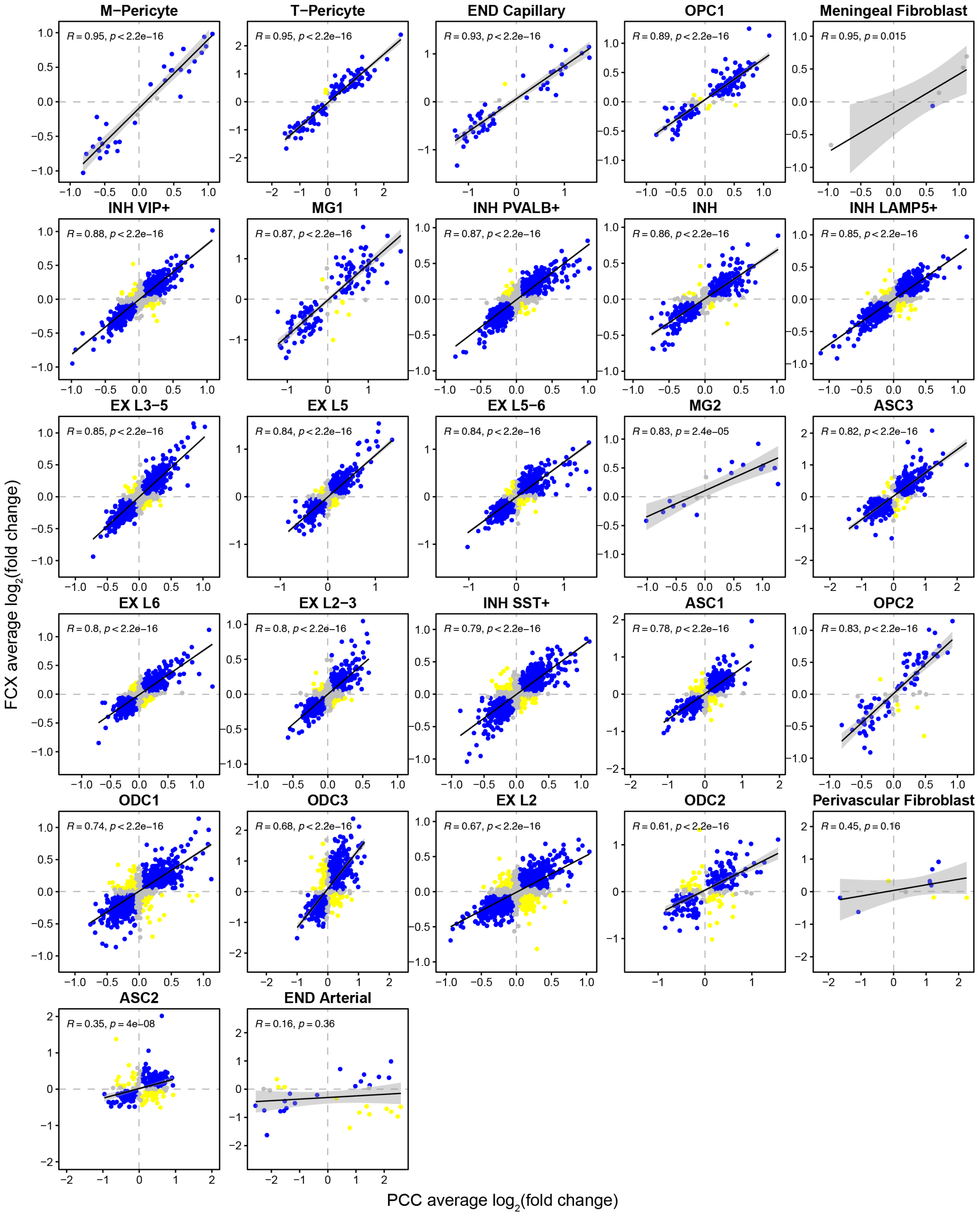


**Fig. S11. Comparison of snRNA-seq AD in DS versus control DEG effect sizes in the PCC and FCX** Comparison of differential expression effect sizes from AD in DS versus control in the PCC and FCX snRNA-seq data. Genes that were statistically significant (adjusted p-value < 0.05) in either comparison were included in this analysis. Genes are colored blue if the direction is consistent, yellow if inconsistent, and grey if the absolute effect sizes were smaller than 0.05. Black line represents a linear regression with a 95% confidence interval shown in grey. Pearson correlation coefficients are shown in the upper left corner of each panel.

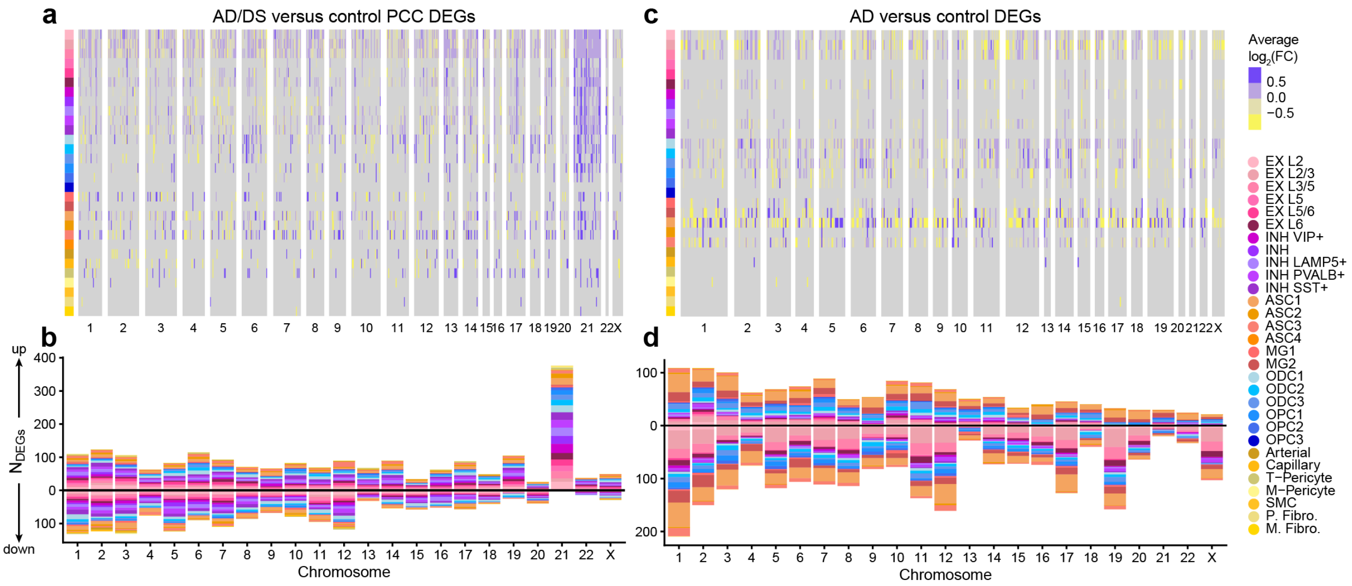


**Fig. S12. snRNA-seq DEGs examined by chromosome a**, Heatmap colored by effect size from the PCC AD in DS versus control differential gene expression analysis, with genes stratified by chromosome and by spatial region. Statistically significant (FDR < 0.05) genes with an absolute average log2(fold-change) >= 0.25 in at least one region are shown. **b**, Stacked bar chart showing the number of PCC AD in DS versus control DEGs in each snRNA-seq cluster stratified by chromosome. **c**, Heatmap colored by effect size from the late-stage AD versus control differential gene expression analysis, with genes stratified by chromosome and by spatial region. Statistically significant (FDR < 0.05) genes with an absolute average log2(fold-change) >= 0.25 in at least one region are shown. **d**, Stacked bar chart showing the number of late-stage AD versus control DEGs in each snRNA-seq cluster stratified by chromosome.

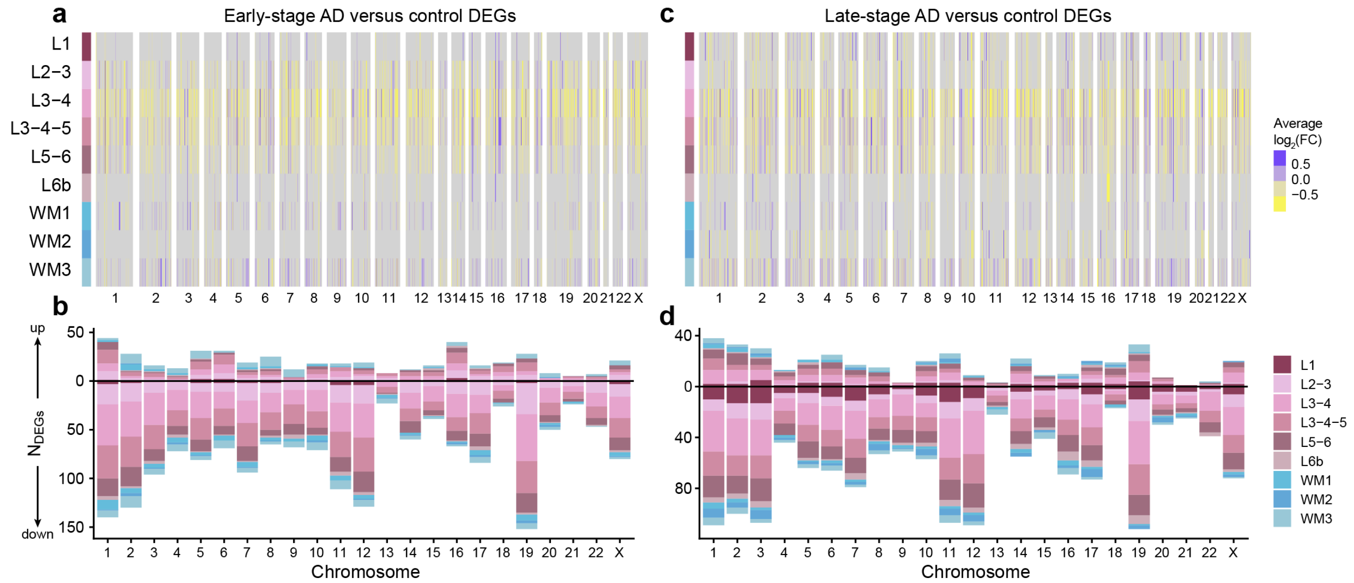


**Fig. S13. Gene set overlap analysis of spatial transcriptomics DEGs from different disease groups a**, Heatmap colored by effect size from the spatial transcriptomic early-stage AD versus control differential gene expression analysis, with genes stratified by chromosome and by spatial region. Statistically significant (FDR < 0.05) genes with an absolute average log2(fold-change) >= 0.25 in at least one region are shown. **b**, Stacked bar chart showing the number of spatial transcriptomic early-stage AD control DEGs in each spatial cluster stratified by chromosome. **c**, Heatmap colored by effect size from the spatial transcriptomic early-stage AD versus control differential gene expression analysis, with genes stratified by chromosome and by spatial region. Statistically significant (FDR < 0.05) genes with an absolute average log2(fold-change) >= 0.25 in at least one region are shown. **d**, Stacked bar chart showing the number of spatial transcriptomic early-stage AD versus control DEGs in each spatial cluster stratified by chromosome.

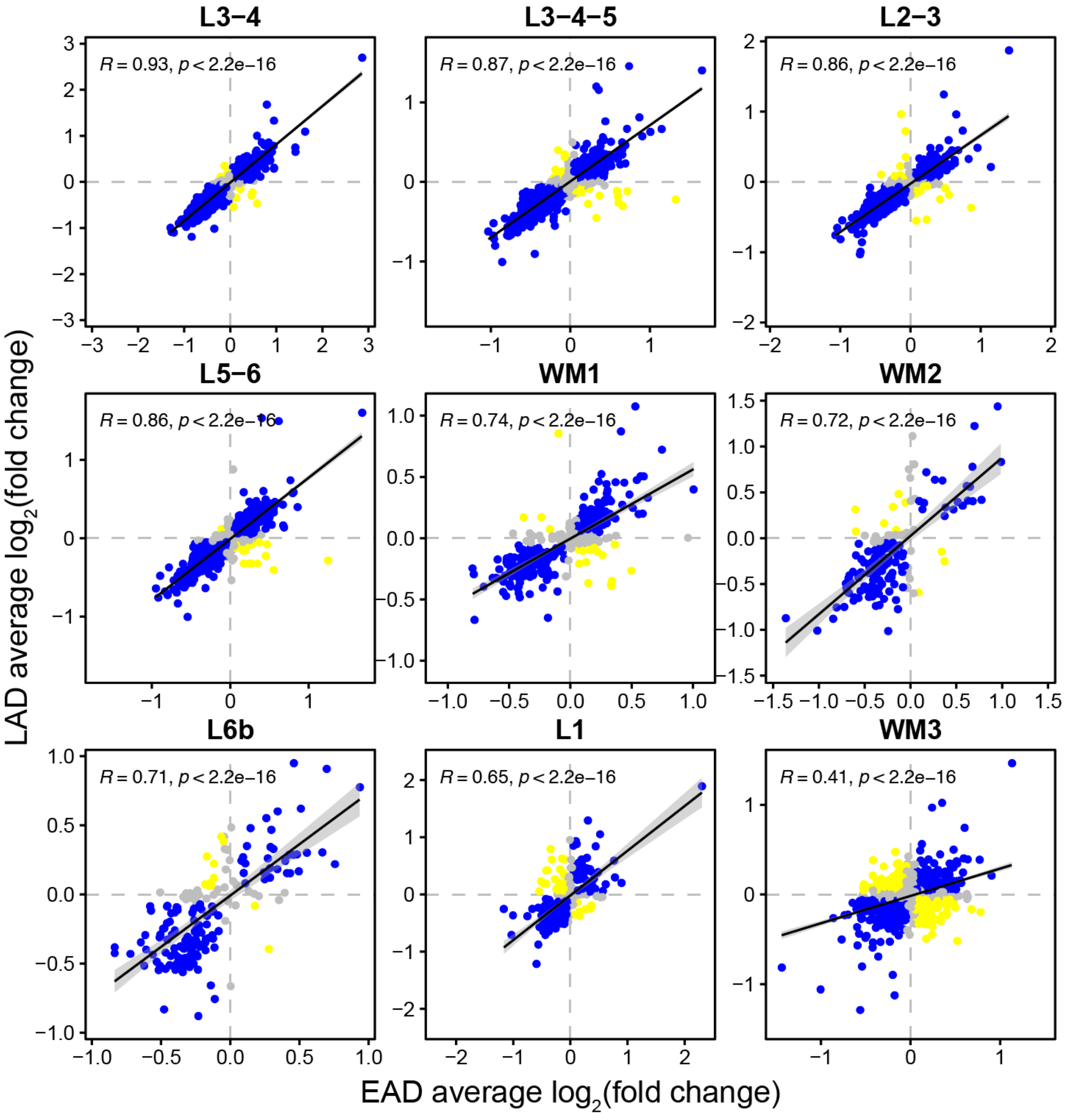


**Fig. S14. Comparison of spatial transcriptomic DEG effect sizes between early-stage AD and late-stage AD** Comparison of differential expression effect sizes from early-stage AD versus control and late-stage AD versus control. Genes that were statistically significant (adjusted p-value < 0.05) in either comparison were included in this analysis. Genes are colored blue if the direction is consistent, yellow if inconsistent, and grey if the absolute effect sizes were smaller than 0.05. Black line represents a linear regression with a 95% confidence interval shown in grey. Pearson correlation coefficients are shown in the upper left corner of each panel.

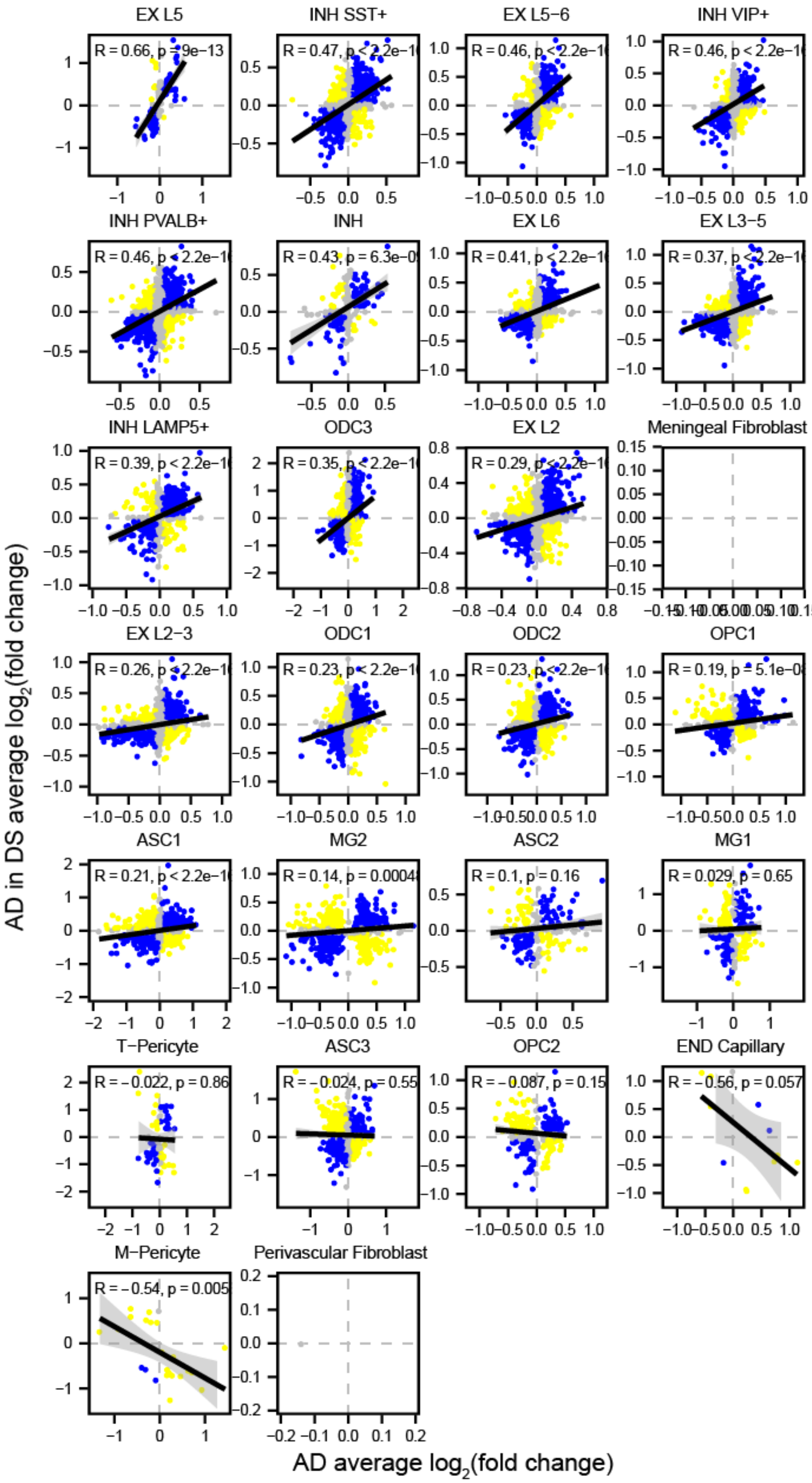


**Fig. S15. Comparison of snRNA-seq DEG effect sizes between sporadic AD and AD in DS** Comparison of differential expression effect sizes from sAD versus control and AD in DS versus control. Genes that were statistically significant (adjusted p-value < 0.05) in either comparison were included in this analysis. Genes are colored blue if the direction is consistent, yellow if inconsistent, and grey if the absolute effect sizes were smaller than 0.05. Black line represents a linear regression with a 95% confidence interval shown in grey. Pearson correlation coefficients are shown in the upper left corner of each panel.

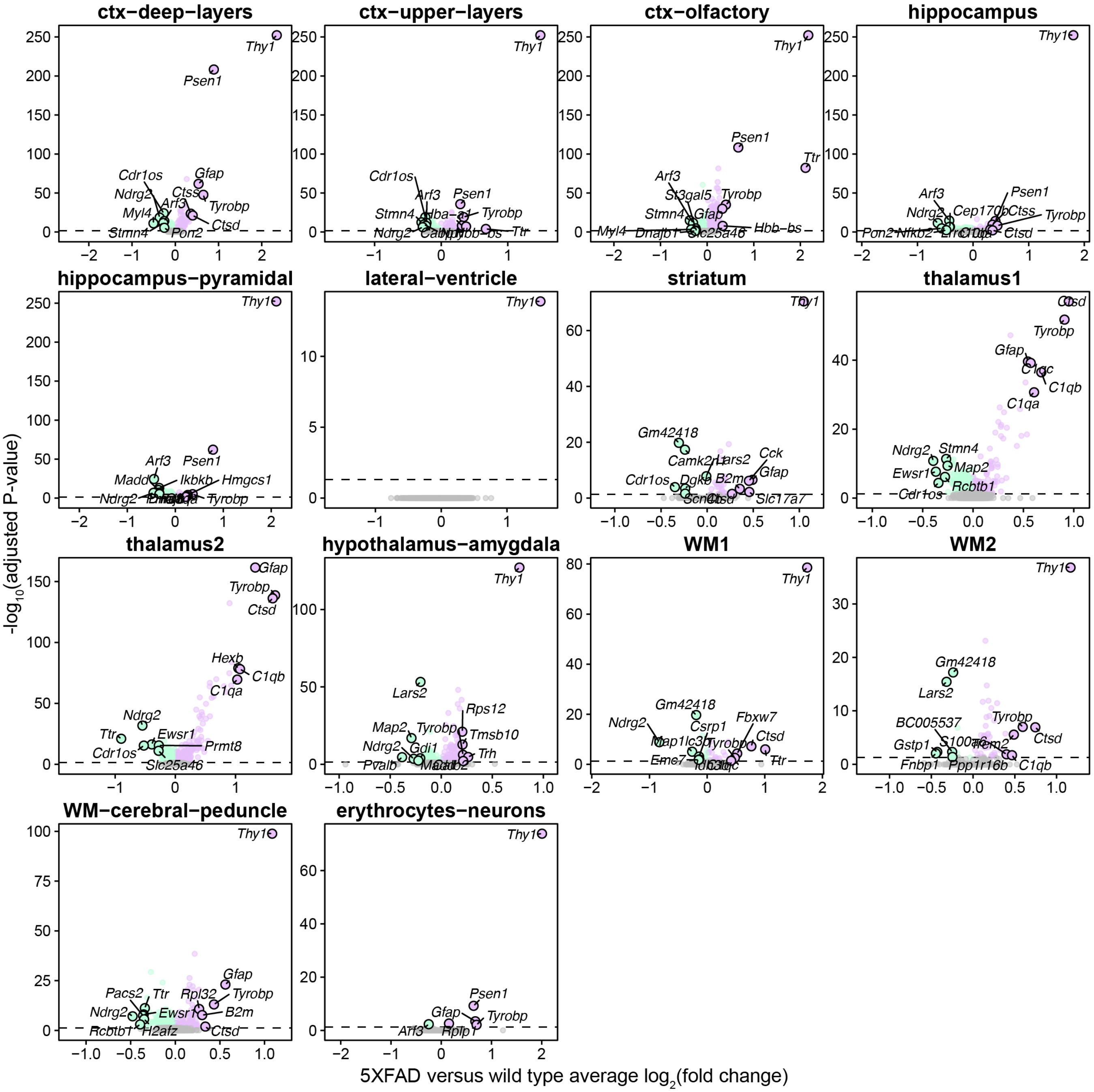


**Fig. S16. 5xFAD versus wild type DEGs at four months of age.** Volcano plots show the adjusted significance levels and effect sizes from the differential gene expression comparisons between 5xFAD versus wild type mice at four months of age. The results are shown for the differential gene expression analysis in each of the spatial transcriptomic clusters. The top and bottom six genes by effect size are annotated.

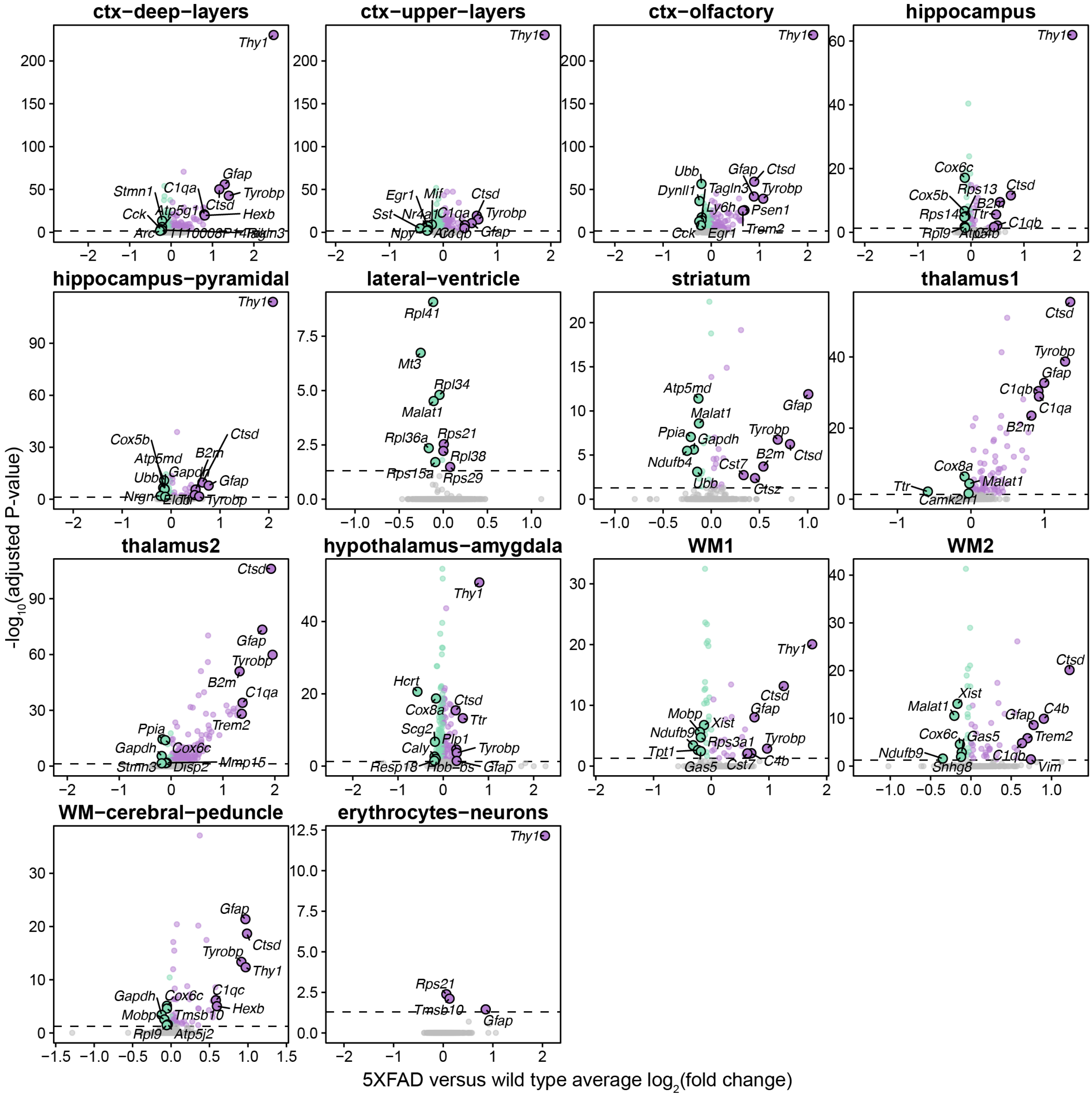


**Fig. S17. 5xFAD versus wild type DEGs at six months of age.** Volcano plots show the adjusted significance levels and effect sizes from the differential gene expression comparisons between 5xFAD versus wild type mice at six months of age. The results are shown for the differential gene expression analysis in each of the spatial transcriptomic clusters. The top and bottom six genes by effect size are annotated.

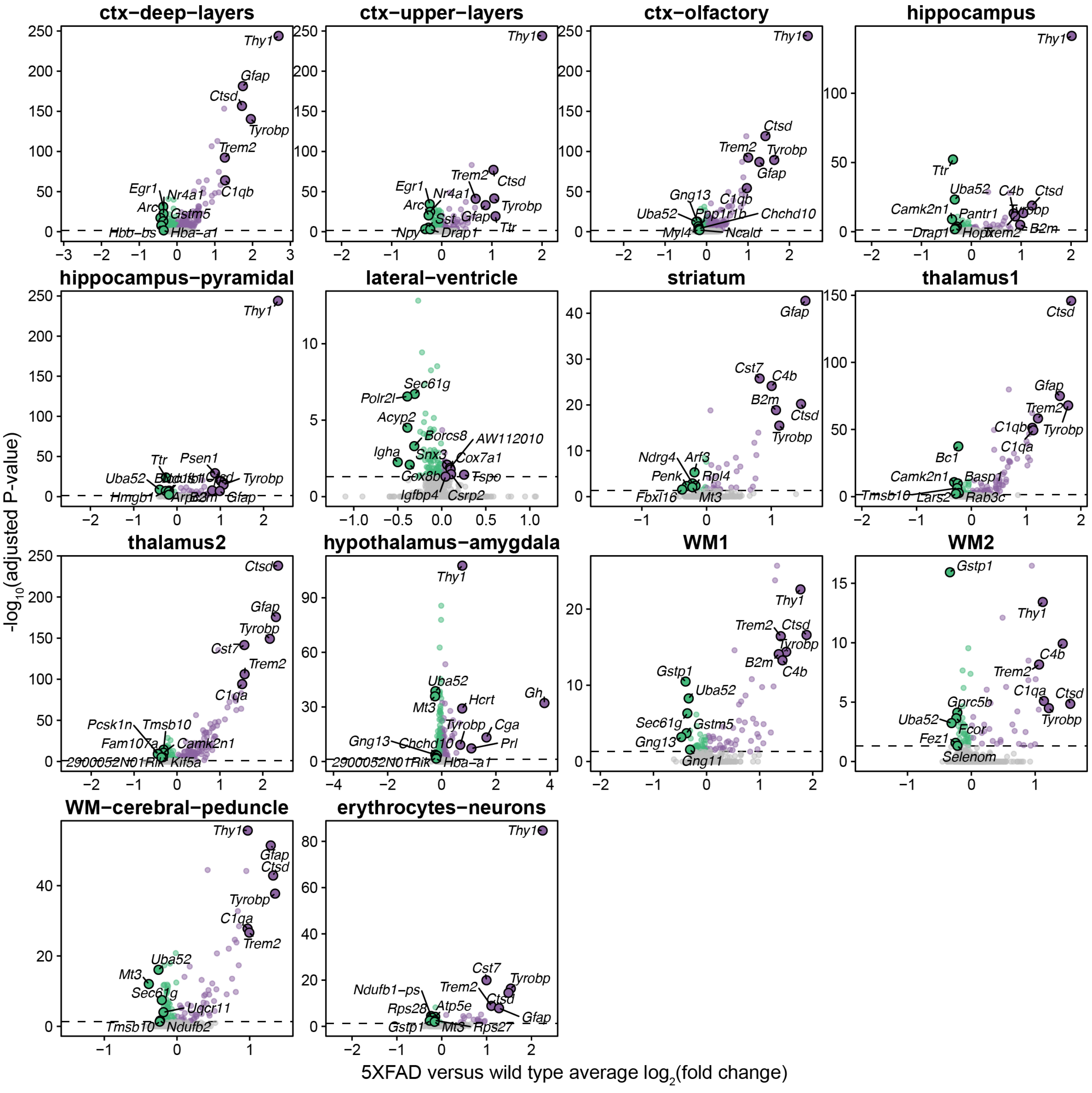


**Fig. S18. 5xFAD versus wild type DEGs at eight months of age.** Volcano plots show the adjusted significance levels and effect sizes from the differential gene expression comparisons between 5xFAD versus wild type mice at six months of age. The results are shown for the differential gene expression analysis in each of the spatial transcriptomic clusters. The top and bottom six genes by effect size are annotated.

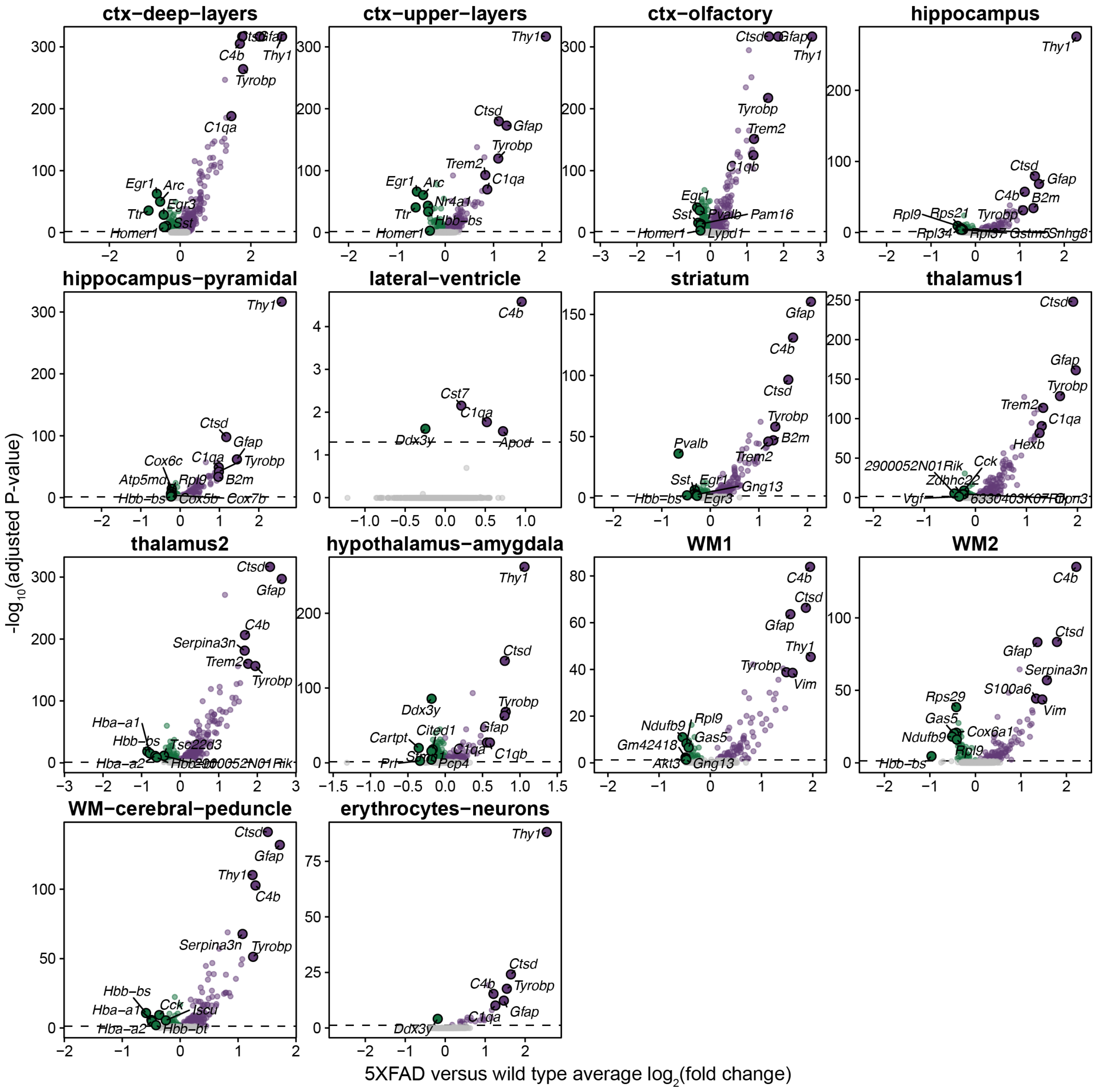


**Fig. S19. 5xFAD versus wild type DEGs at twelve months of age.** Volcano plots show the adjusted significance levels and effect sizes from the differential gene expression comparisons between 5xFAD versus wild type mice at six months of age. The results are shown for the differential gene expression analysis in each of the spatial transcriptomic clusters. The top and bottom six genes by effect size are annotated.

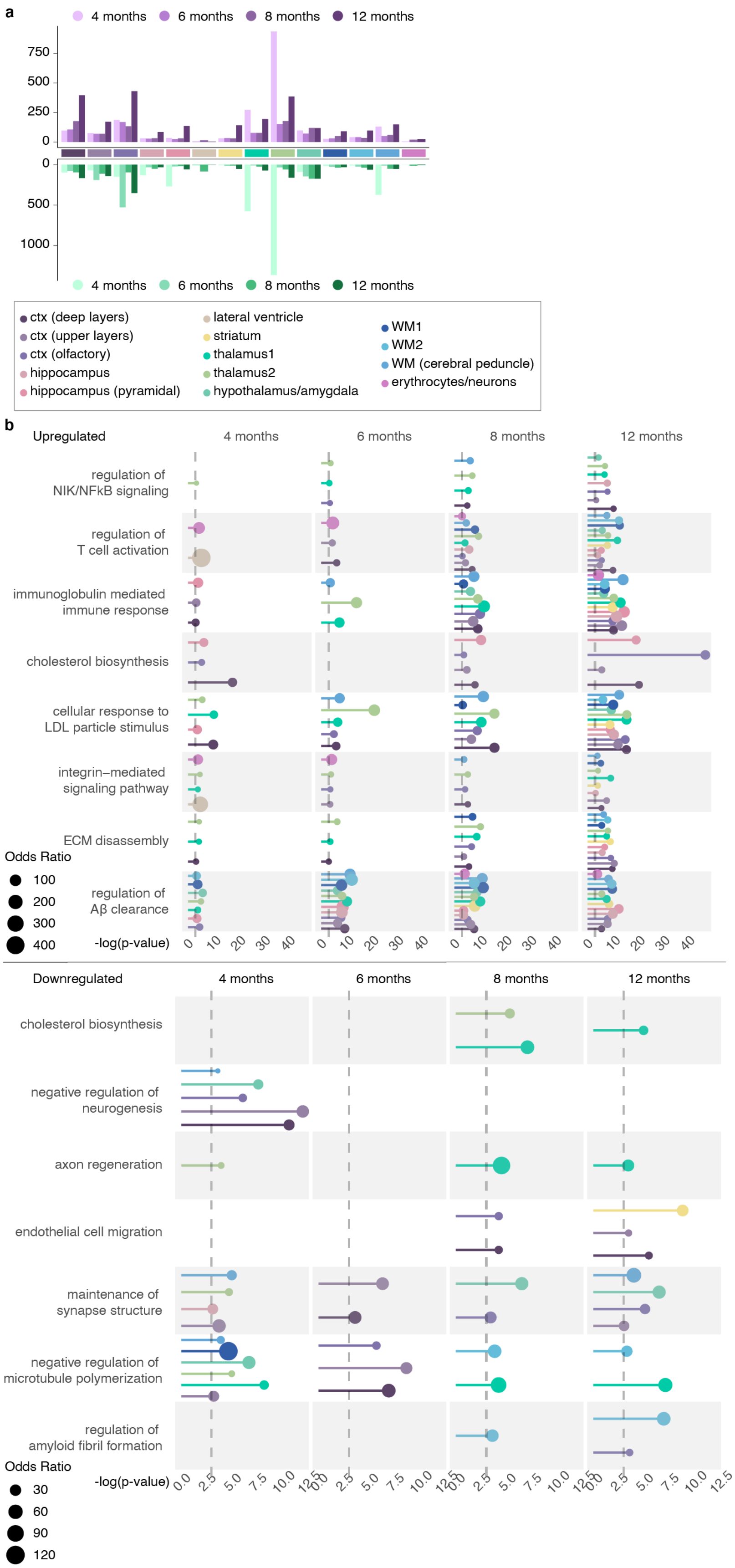


**Fig. S20. Number of 5xFAD DEGs and GO term enrichment analysis. a**, Barplot showing the number of DEGs in each group. **b**, Selected pathway enrichment analysis for the 5xFAD DEGs.

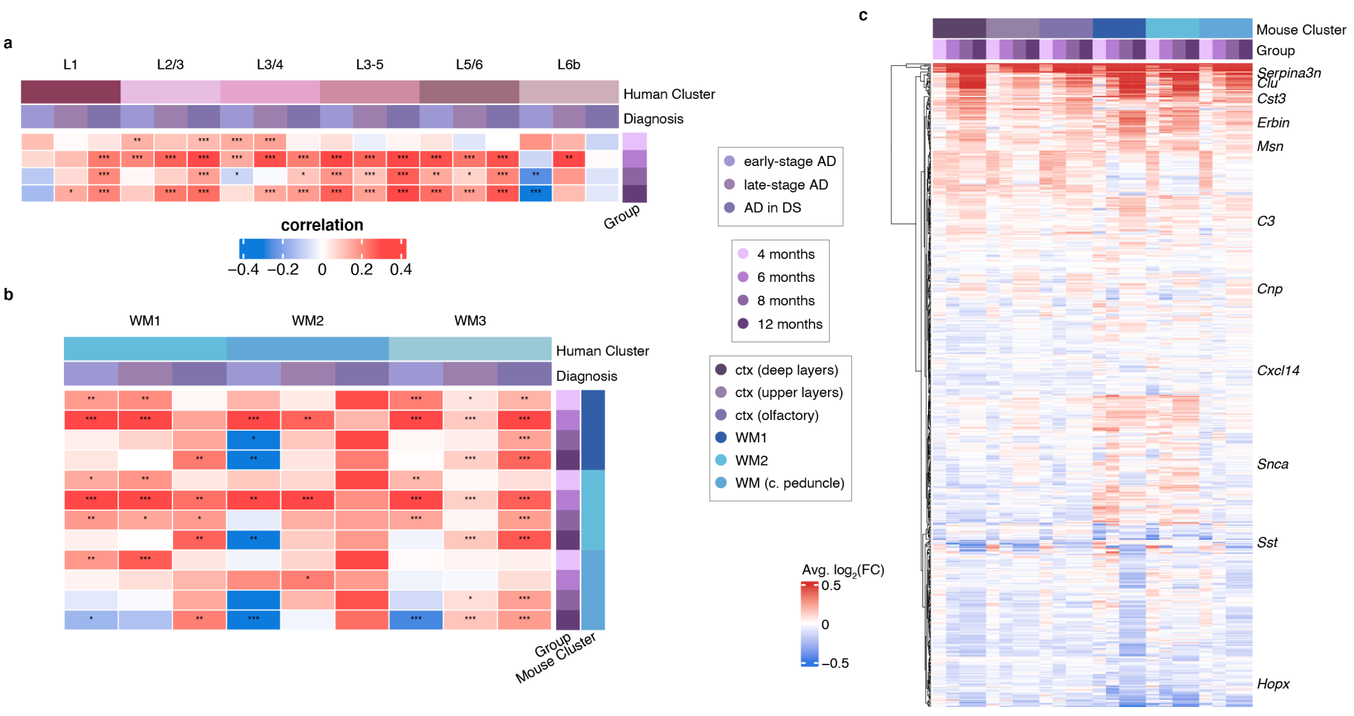


**Fig. S21. Comparisons of human and mouse DEGs. a-b**, Heatmaps showing correlations of effect sizes between human and mouse ST datasets in orthologous genes between human and mouse for cortical layer clusters **a** and white matter **b** clusters. **c**, Heatmap showing differential gene expression effect size results in the 5xFAD versus WT comparisons.

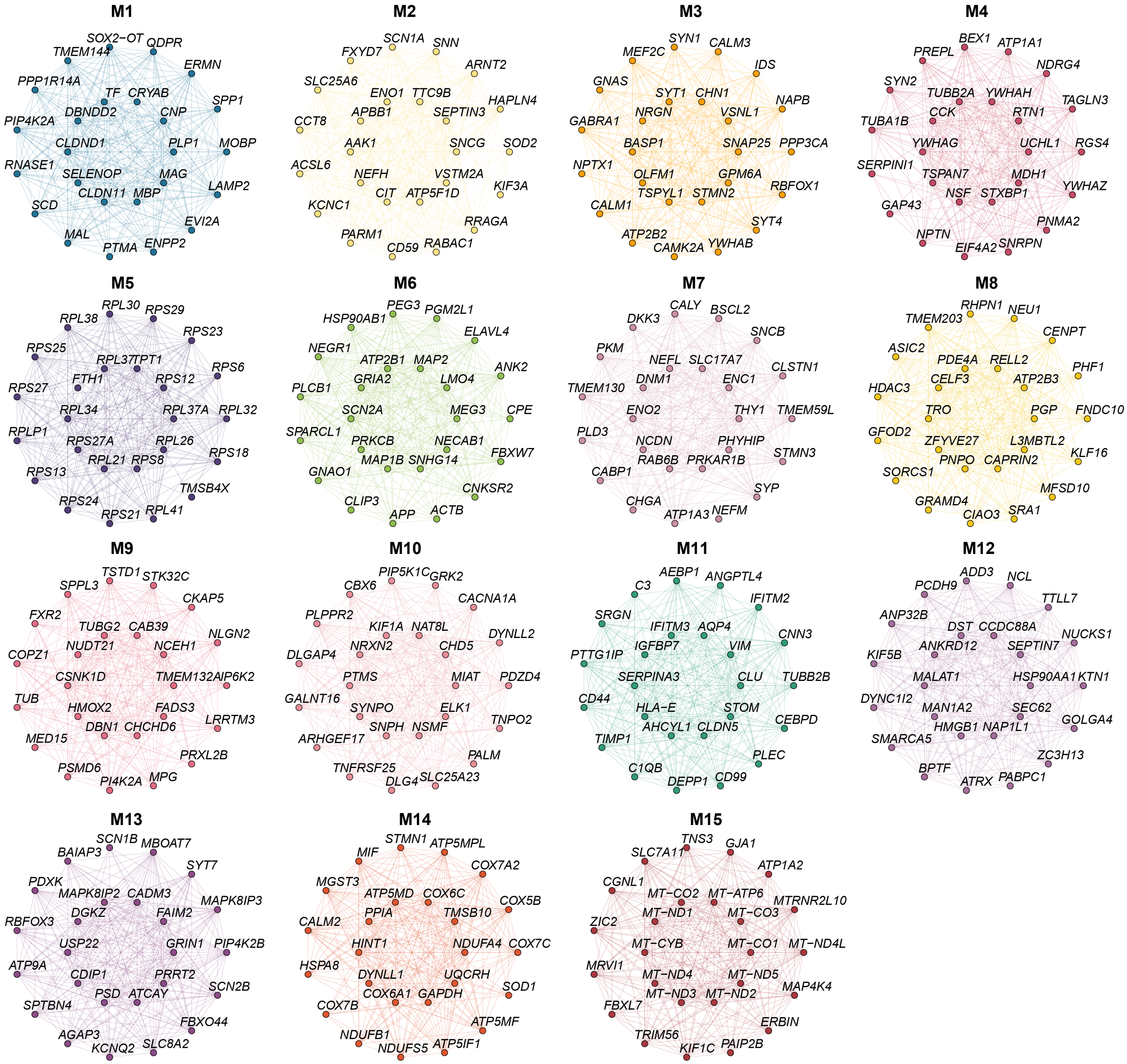


**Fig. S22. Module hub gene networks for the human ST co-expression meta-modules.** Hub gene networks for each of the 15 human spatial co-expression meta-modules. The top 25 hub genes ranked by kME are visualized. Nodes represent genes, and edges represent co-expression links.

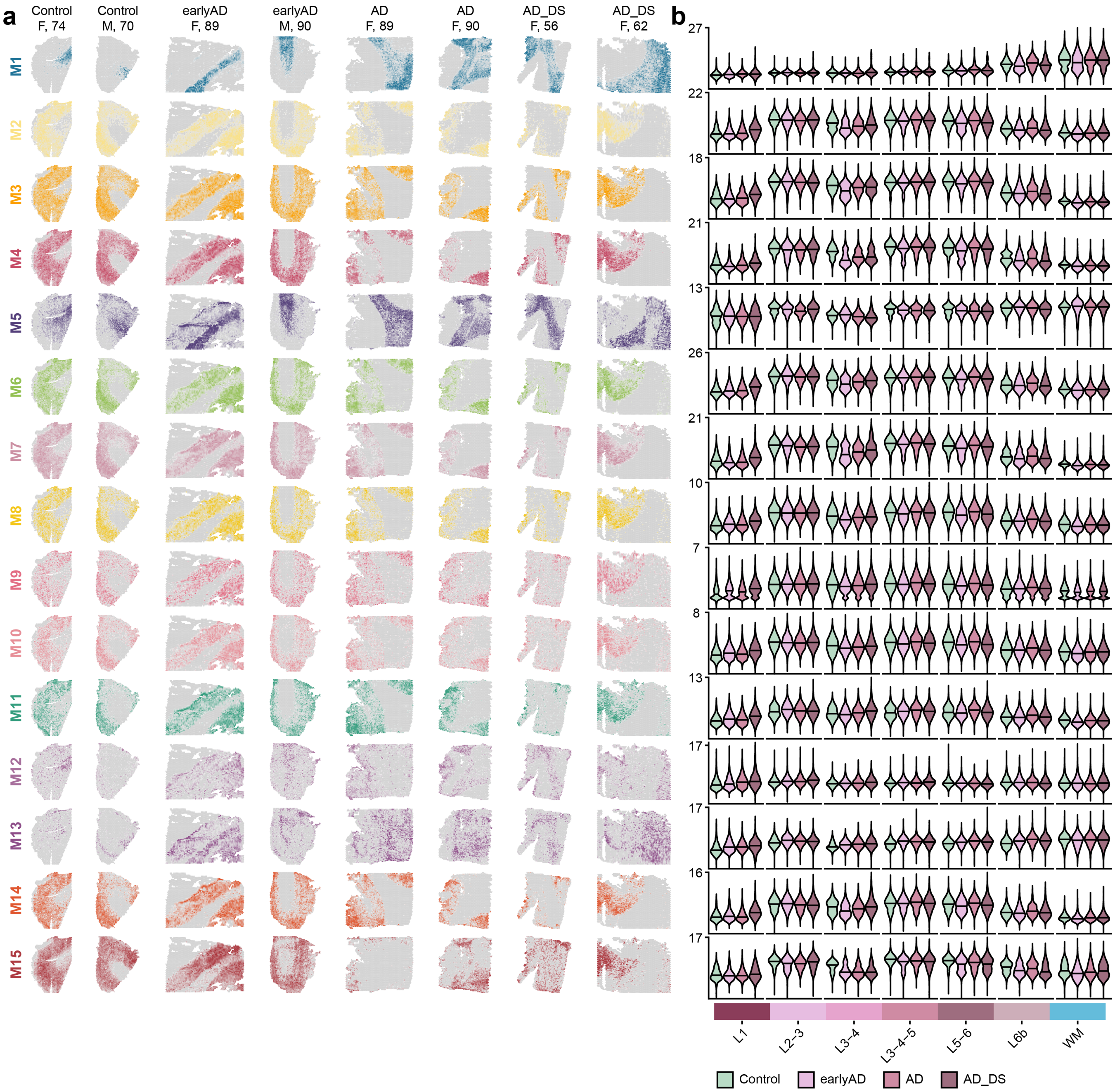


**Fig. S23. Module eigengenes for the human ST meta-modules. a**, Spatial feature plots showing module eigengenes (MEs) for the 15 human co-expression meta-modules shown in eight representative samples. **b**, ME distributions in each disease group (control, early-stage AD, late-stage AD, AD in DS) stratified by cortical layer clusters and white matter.

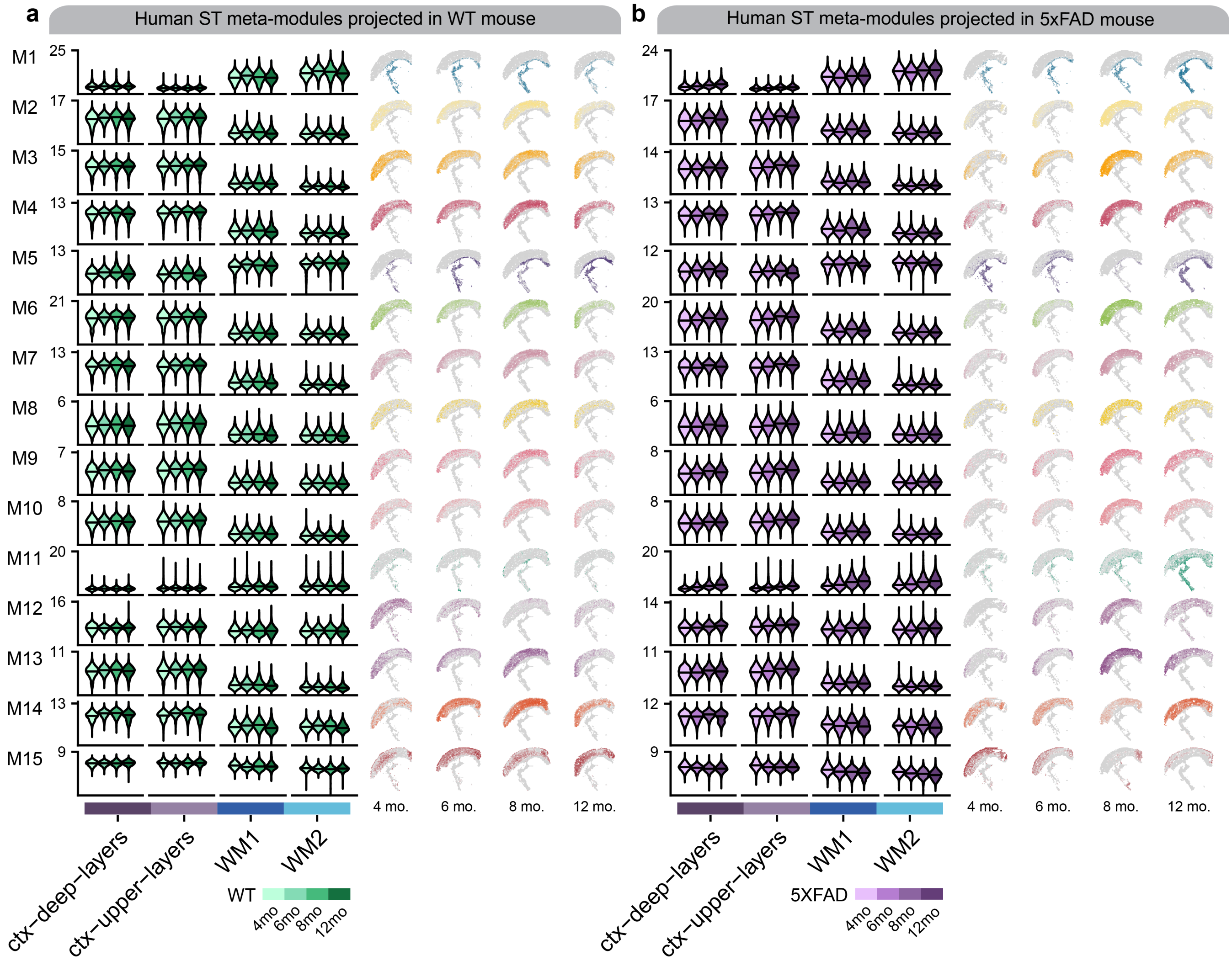


**Fig. S24. Module eigengenes of human co-expression meta-modules in the mouse ST dataset. a-b**, Left: Module eigengene (ME) distributions for the 15 human co-expression meta-modules in each mouse age group (control, early-stage AD, late-stage AD, AD in DS) stratified by cortical and white matter clusters. Right: Spatial feature plots showing MEs in four representative samples. Panel a shows the results in wild type mice and panel **b** shows the results in 5xFAD.

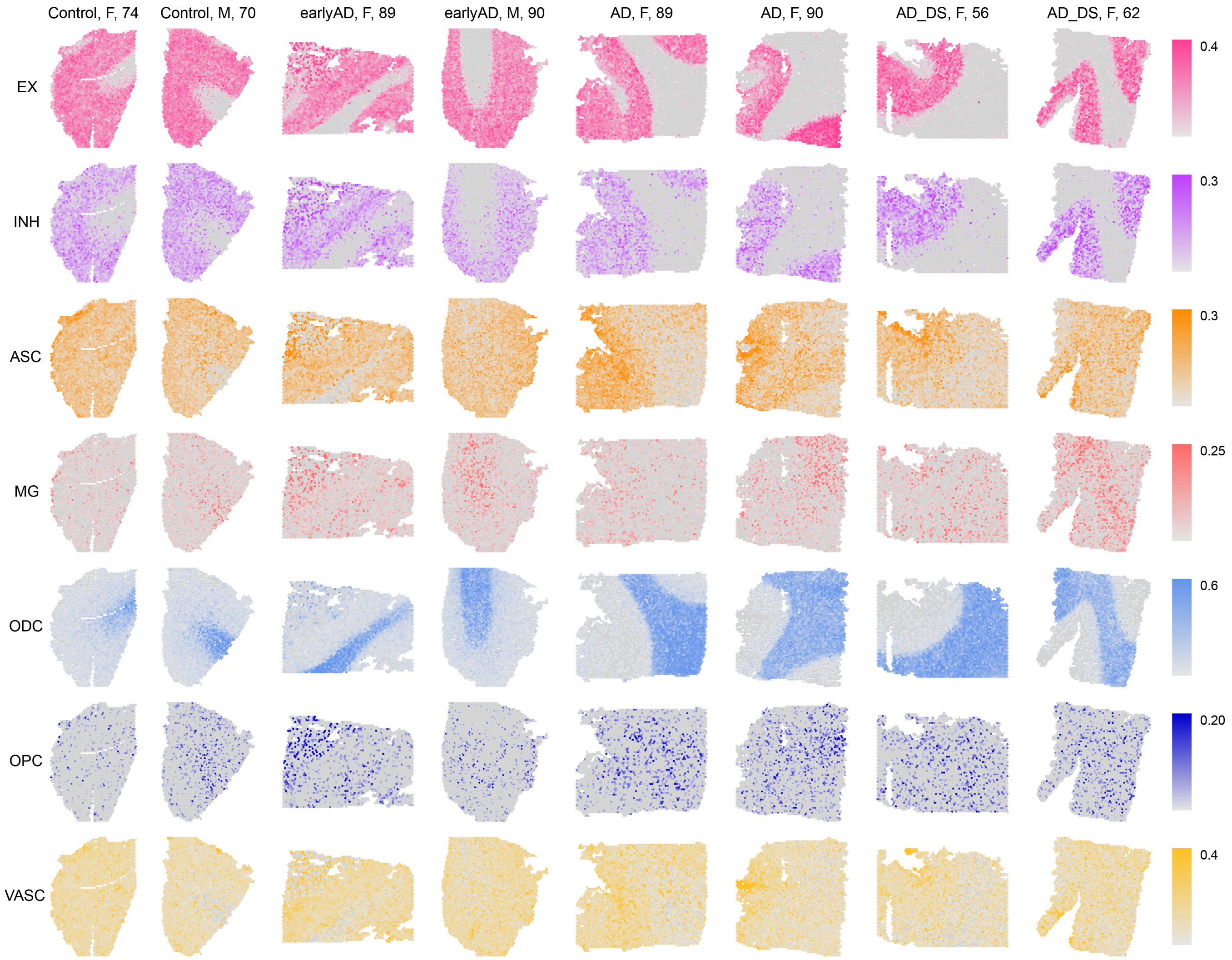


**Fig. S25. Cell-type deconvolution results in the human ST dataset.** Spatial feature plots in eight representative samples from the human ST dataset showing the inferred proportion of different major cell types based on our deconvolution analysis conducted with SPOTLight[4]..

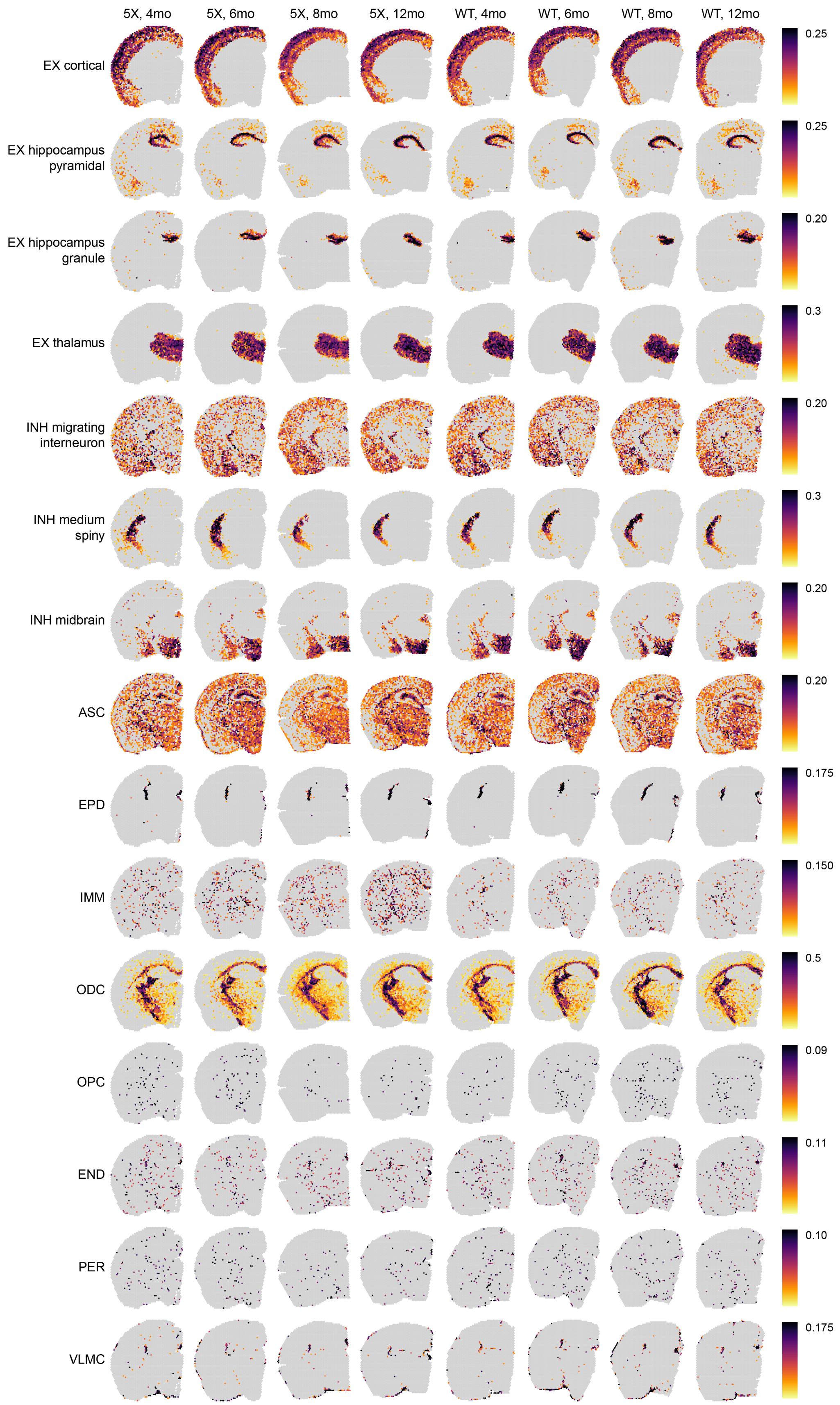


**Fig. S26. Cell-type deconvolution results in the mouse ST dataset.** Spatial feature plots in eight representative samples from the mouse ST dataset showing the inferred proportion of different major cell types based on our deconvolution analysis conducted with SPOTLight[4].

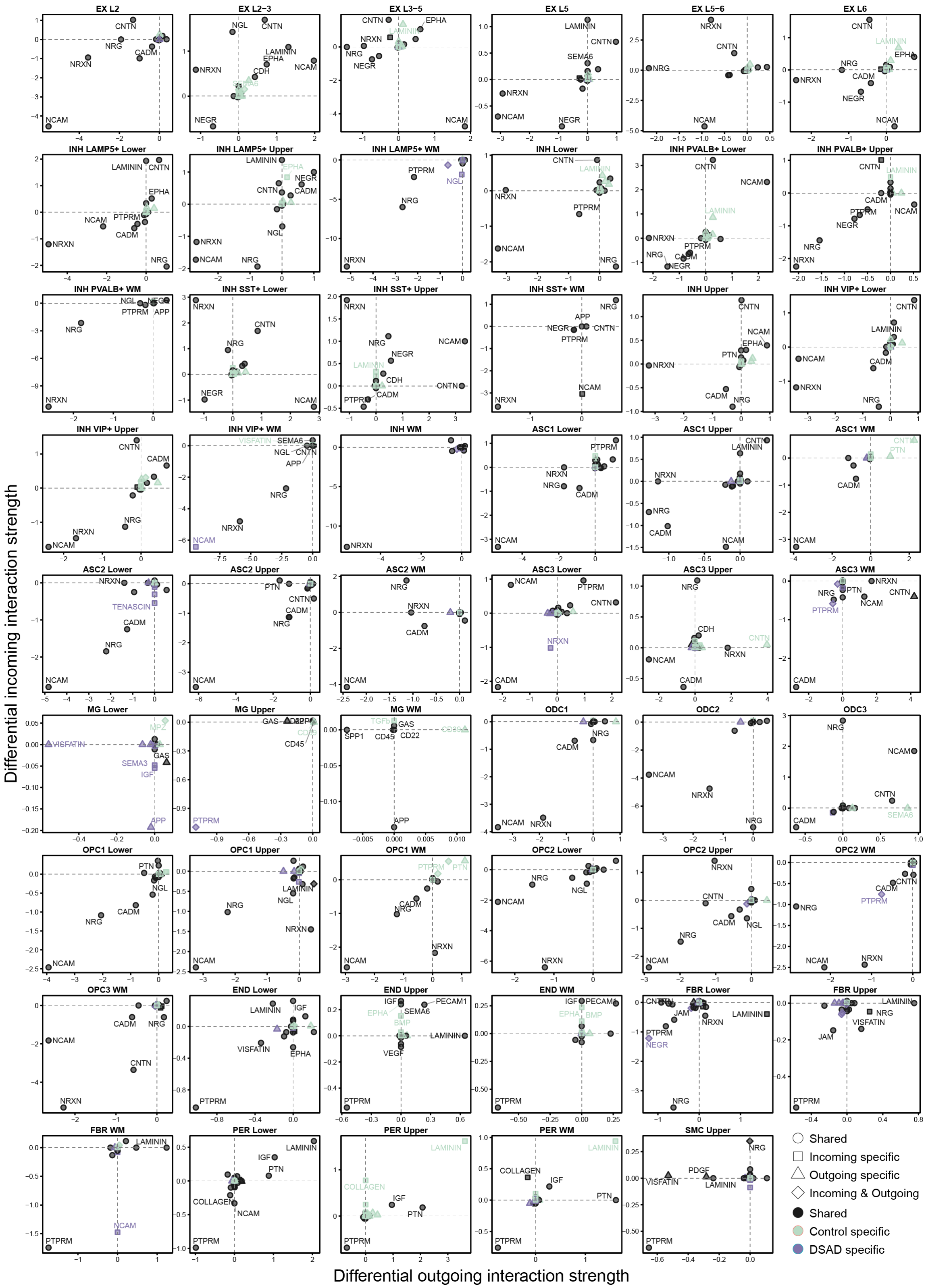


**Fig. S27. Differential cell-cell signaling between AD in DS and control.** Scatter plots showing the differential outgoing interaction strength versus the differential incoming interaction strength from the differential cell-cell signaling network analysis between AD in DS cases versus cognitively normal controls. Pathways were shown in each cluster where there was a statistically significant ($p$-value < 0.05) difference between AD in DS and control based on a permutation test[5].

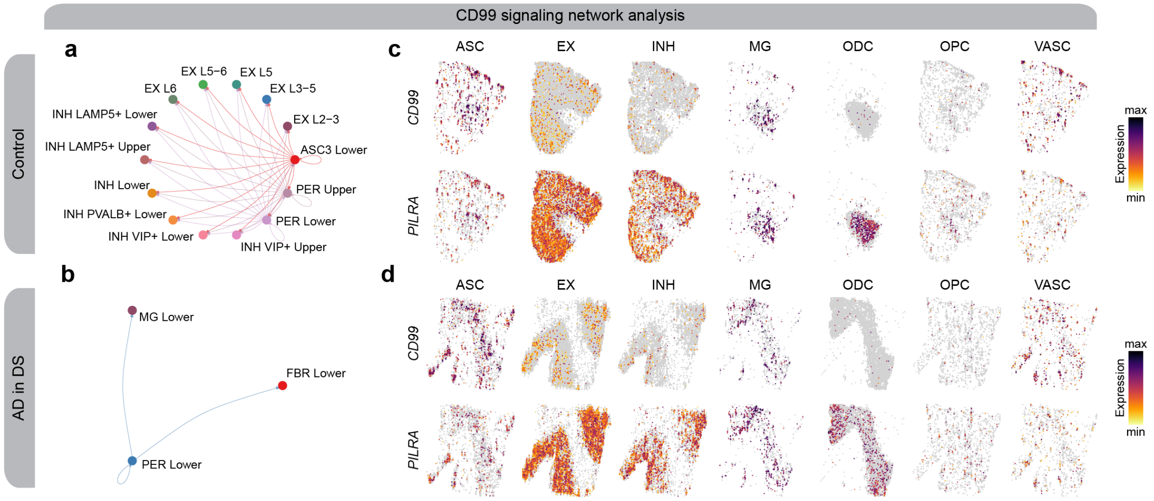


**Fig. S28. CD99 signaling changes between AD in DS and control.** textbfa-b, Network plot showing the CCC signaling strength between different cell populations in controls (**a**) and AD in DS (**b**) for the CD99 signaling pathway. **c-d**, Spatial feature plots of the snRNA-seq in predicted spatial coordinates for one control sample (**c**) and one AD in DS sample (**d**) for one ligand and one receptor in the CD99 pathway.

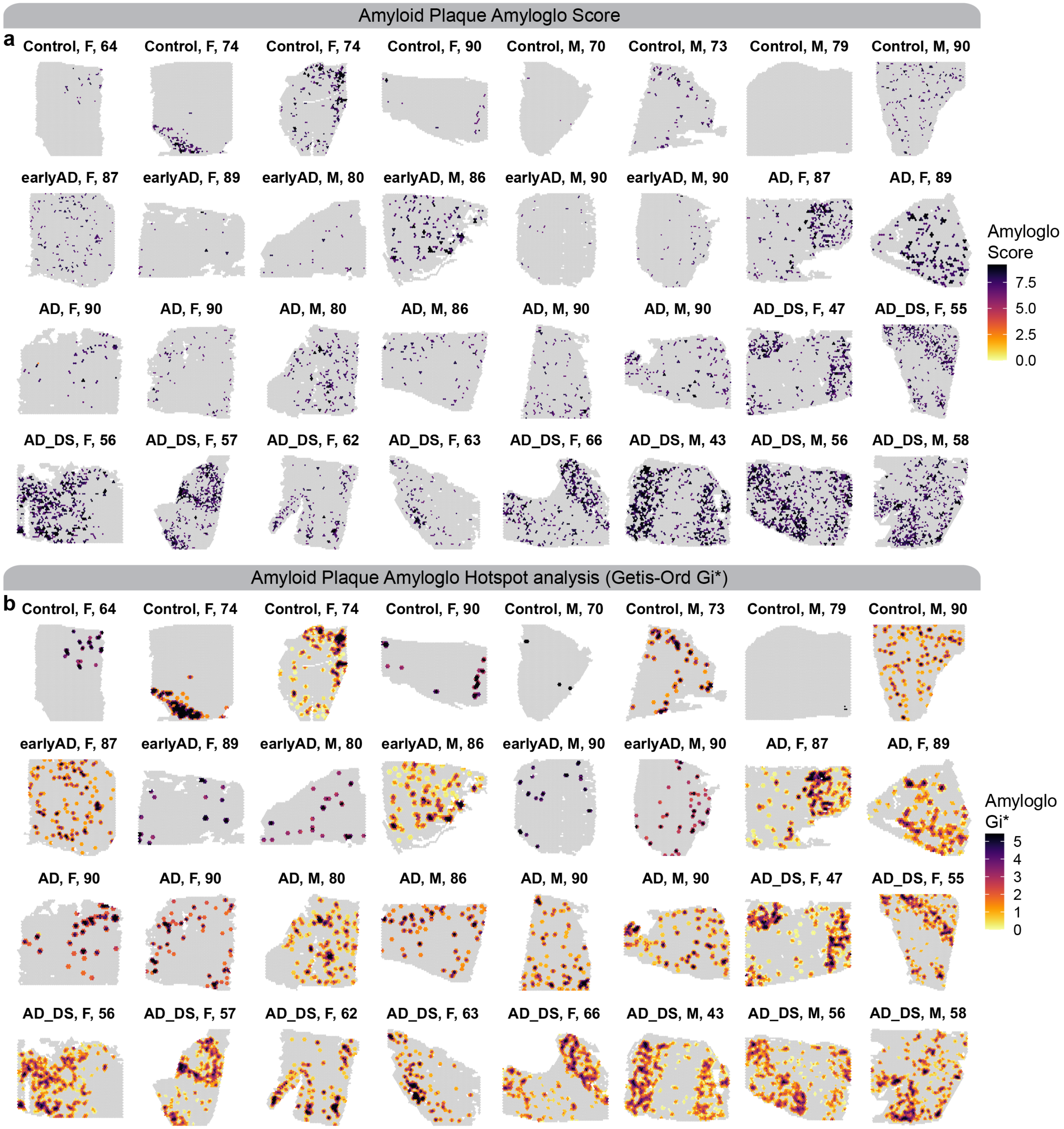


**Fig. S29. Integration of amyloid plaque imaging (Amylo-glo) and hotspot analysis in the human ST dataset** Spatial feature plots for the human ST dataset showing the integration of amyloid plaque imaging of Amylo-glo. a, The Amylo-glo score, computed as the sum of the areas (size) for all overlapping amyloid plaques with each ST spot, and then log normalized. b, Getis-Ord Gi* hotspot analysis based on the Amylo-glo score.

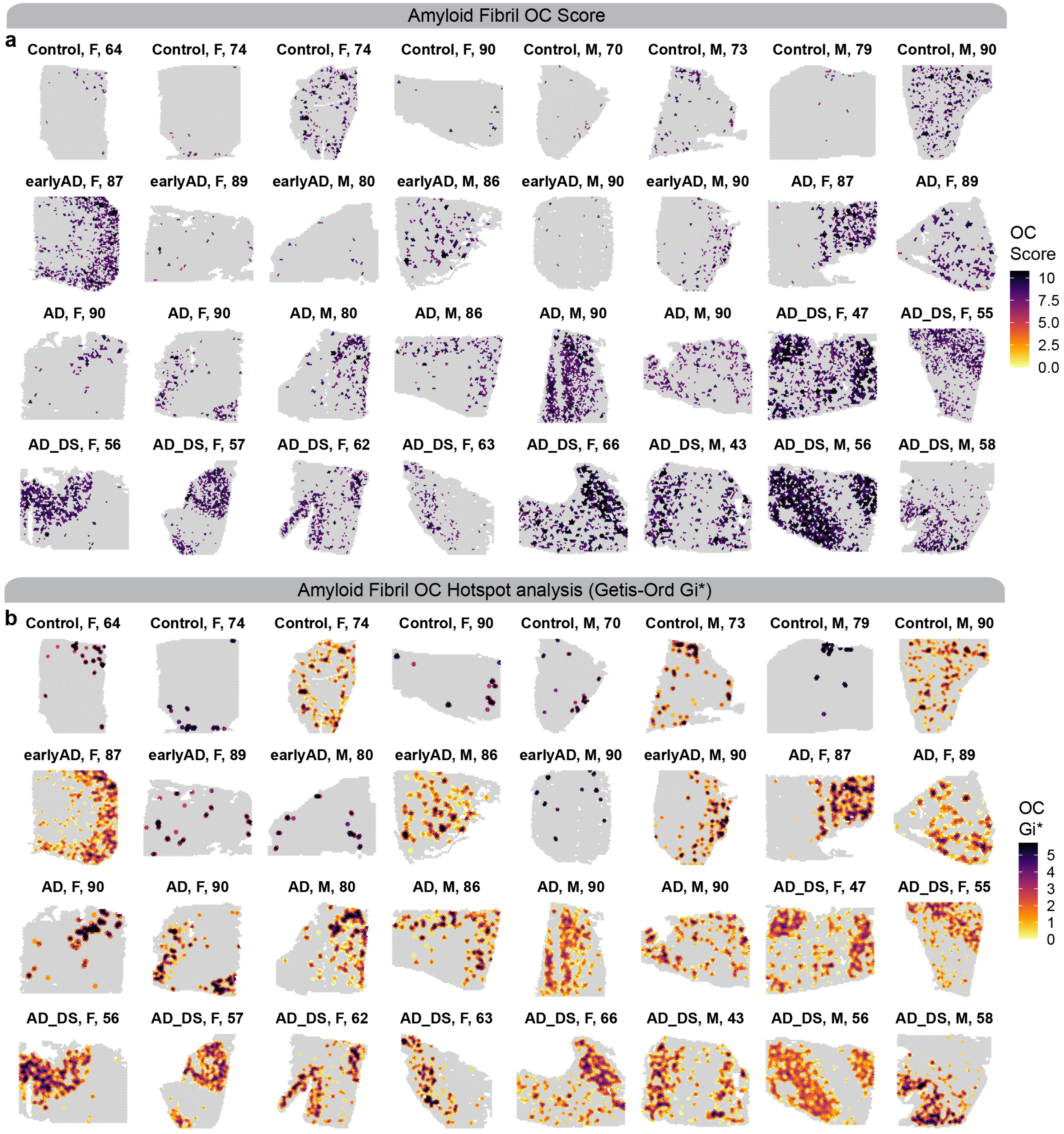


**Fig. S30. Integration of amyloid fibril imaging (OC) and hotspot analysis in the human ST dataset.** Spatial feature plots for the human ST dataset showing the integration of amyloid fibril imaging of OC. a, The OC score, computed as the sum of the areas (size) for all overlapping amyloid plaques with each ST spot, and then log normalized. b, Getis-Ord Gi* hotspot analysis based on the OC score.

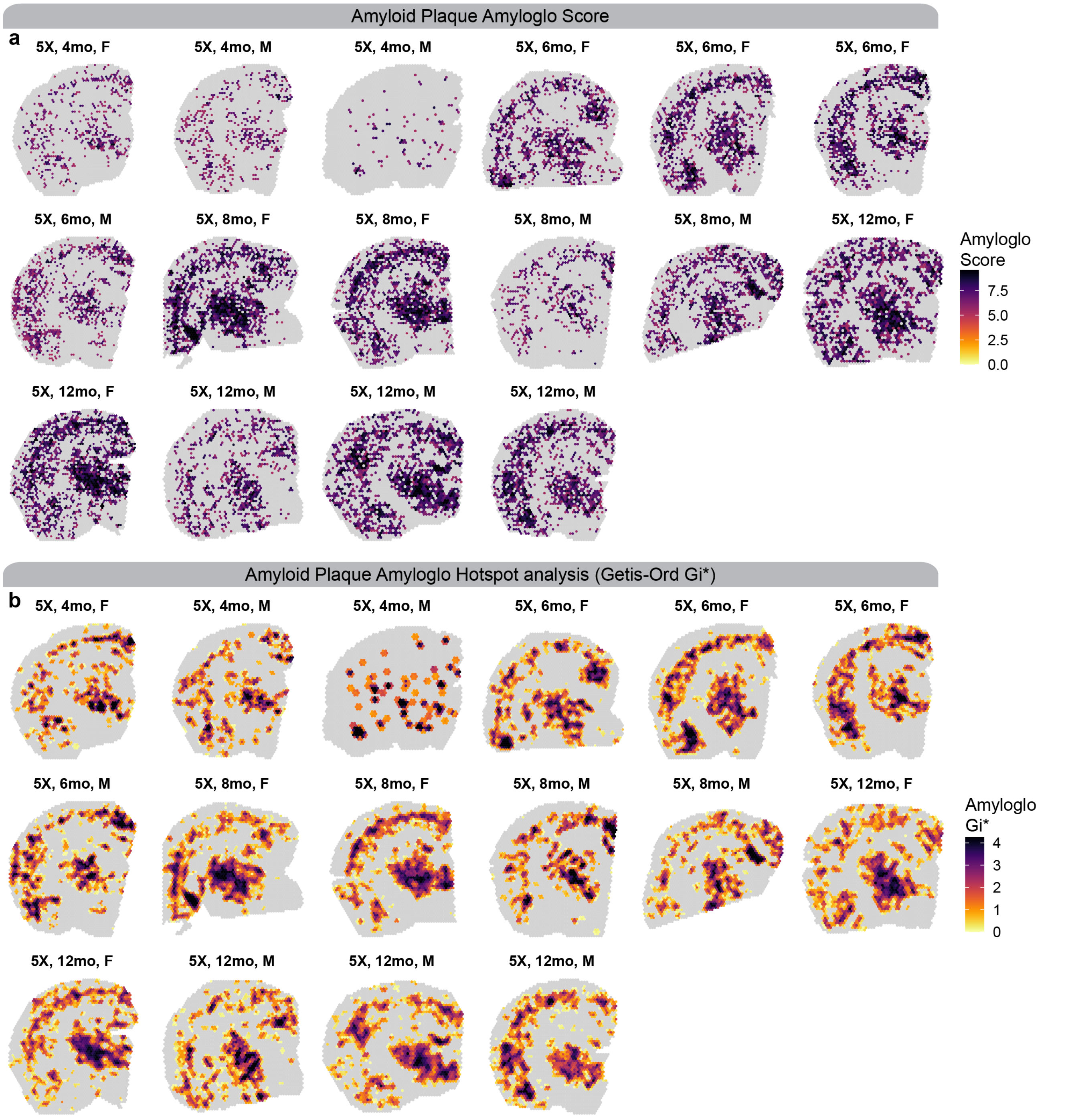


**Fig. S31. Integration of amyloid plaque imaging (Amylo-glo) and hotspot analysis in the mouse ST dataset** Spatial feature plots for the mouse ST dataset showing the integration of amyloid plaque imaging of Amylo-glo. a, The Amylo-glo score, computed as the sum of the areas (size) for all overlapping amyloid plaques with each ST spot, and then log normalized. b, Getis-Ord Gi* hotspot analysis based on the Amylo-glo score.

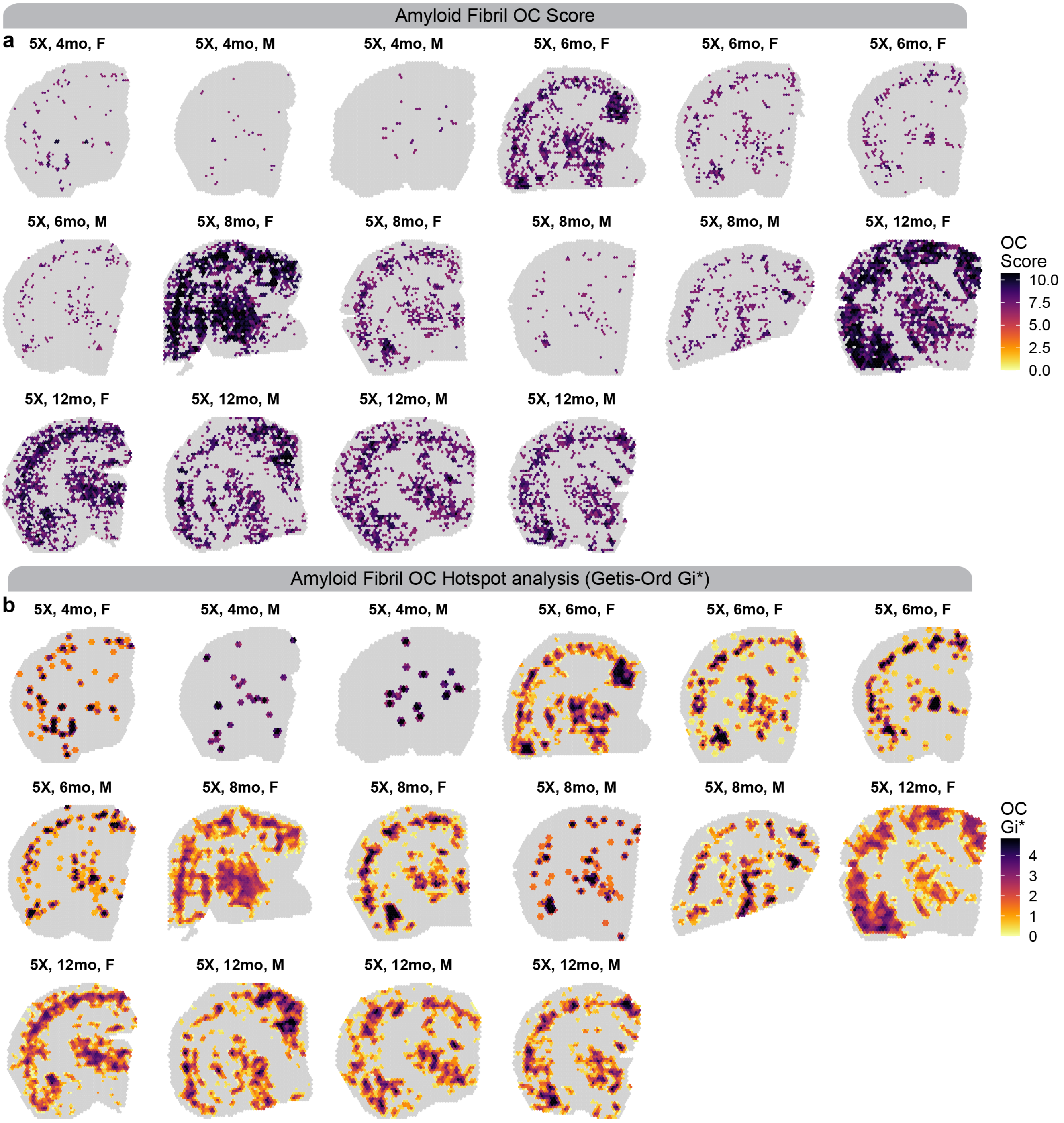


**Fig. S32. Integration of amyloid fibril imaging (OC) and hotspot analysis in the mouse ST dataset** Spatial feature plots for the mouse ST dataset showing the integration of amyloid fibril imaging of OC. a, The OC score, computed as the sum of the areas (size) for all overlapping amyloid plaques with each ST spot, and then log normalized. b, Getis-Ord Gi* hotspot analysis based on the OC score.

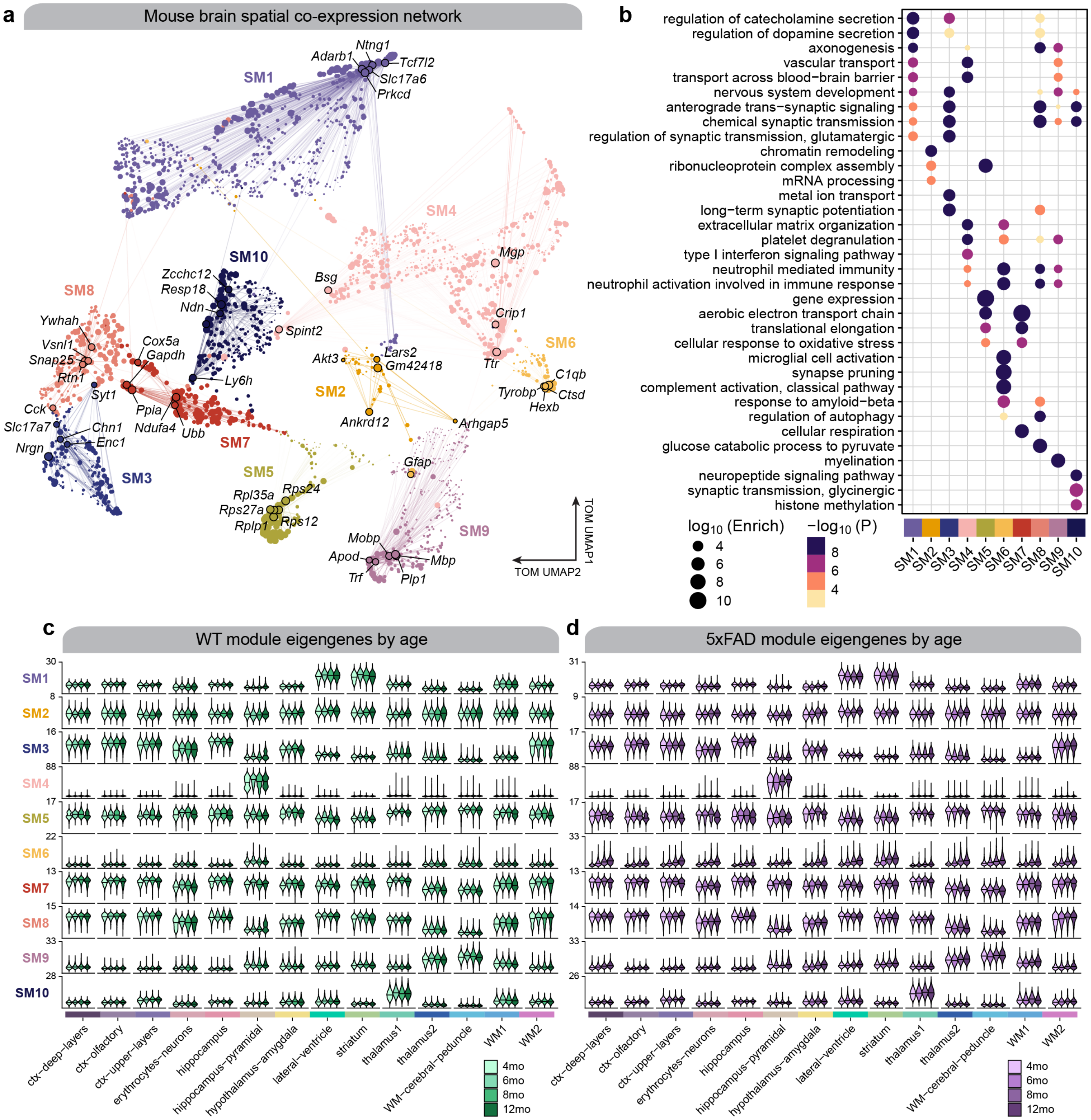


**Fig. S33. Co-expression network analysis in the mouse ST dataset a**, UMAP plot of the mouse spatial co-expression network. Each node represents a single gene, and edges represent co-expression links between genes and module hub genes. Point size is scaled by eigengene-based connectivity. Nodes are colored by co-expression module assignment. The top five hub genes per module are labeled. Network edges were downsampled for visual clarity. **b**, Dot plot showing selected GO enrichment results for each co-expression module. **c-d**, Module eigengene (ME) distributions for the ten mouse co-expression modules in each mouse age group (control, early-stage AD, late-stage AD, AD in DS) stratified by cluster for wild type (**c**) and 5xFAD mice (**d**).

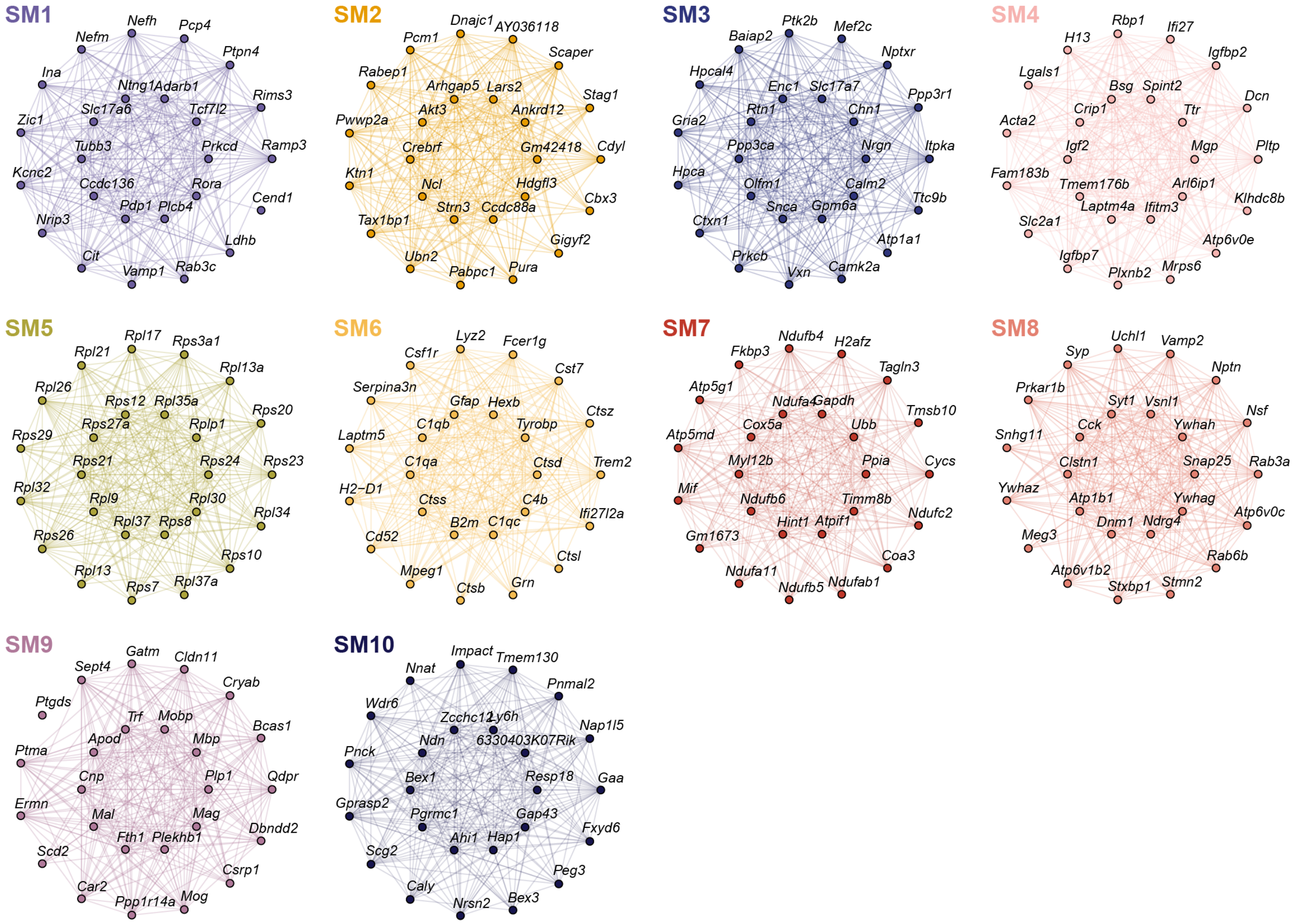


**Fig. S34. Module hub gene networks from the mouse co-expression network analysis** Hub gene networks for each of the 10 mouse spatial co-expression modules. The top 25 hub genes ranked by kME are visualized. Nodes represent genes, and edges represent co-expression links.

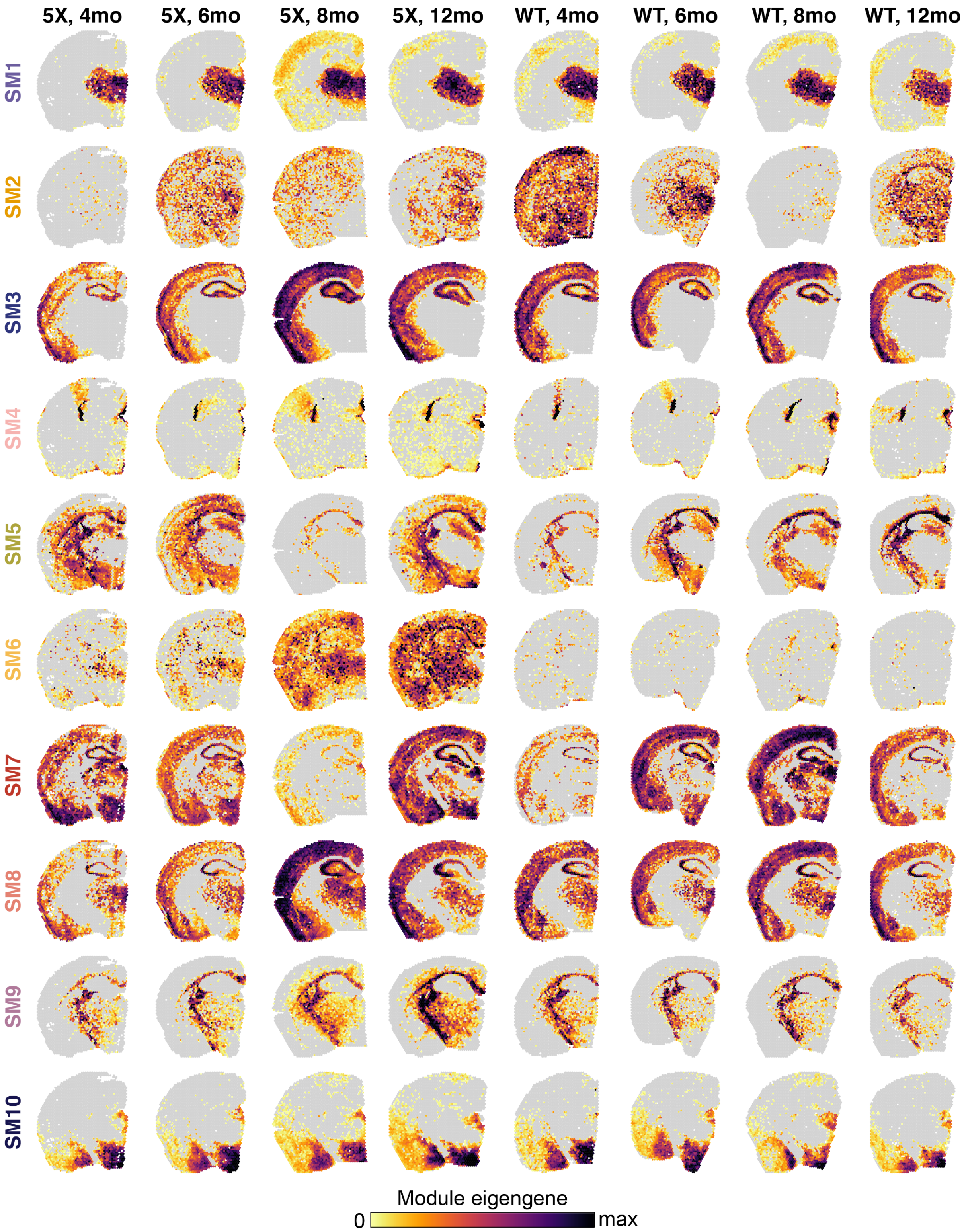


**Fig. S35. Module eigengenes in representative samples from the mouse ST dataset** Spatial feature plots showing module eigengenes (MEs) for the ten mouse co-expression modules shown in eight representative samples.

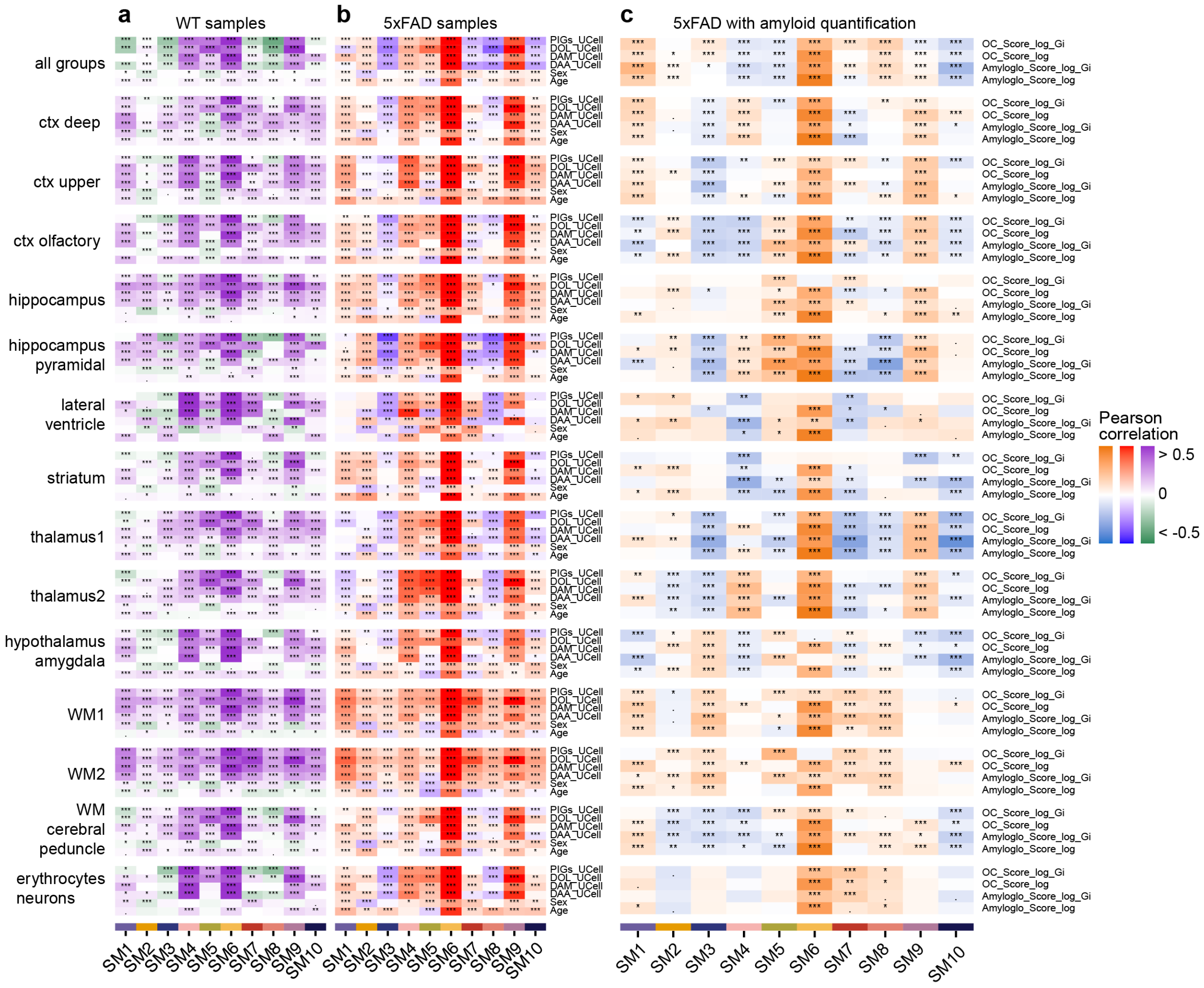

**Fig. S36. Module-trait correlation analysis in the mouse co-expression network** a-b, Heatmap showing the module-trait correlation results for age, sex (positive correlation corresponds to female), and gene signatures for plaque-induced genes (PIGs), disease-associated oligodendrocytes (DOL), disease-associated microglia (DAM), and disease-associated astrocytes (DAA) in wild type (a) and in 5xFAD (b) mice. c, Heatmap showing the module-trait correlation results with the amyloid imaging analysis (Amylo-glo and OC) in 5xFAD with amyloid quantifications. Not significant (ns), p > 0.05; * p <= 0.05; ** p <= 0.01; *** p <= 0.001, **** p<= 0.0001.

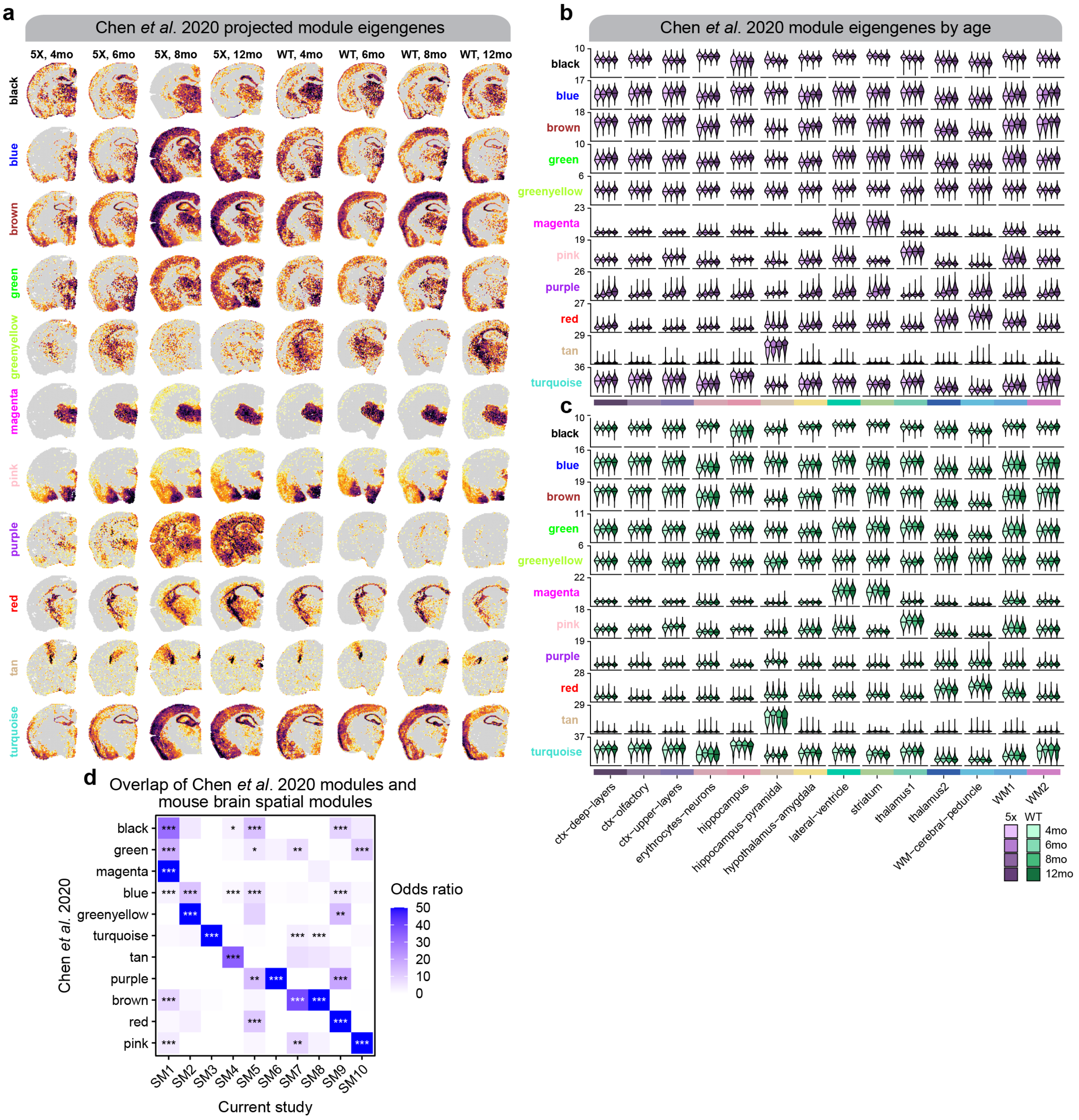


**Fig. S37. Investigating the gene co-expression modules from Chen et al in our 5xFAD ST dataset a**, Spatial feature plots showing module eigengenes (MEs) for the ten mouse co-expression modules shown in eight representative samples. **b-c**, Module eigengene (ME) distributions for the ten mouse co-expression modules in each mouse age group (control, early-stage AD, late-stage AD, AD in DS) stratified by cluster for 5xFAD (**b**) and 5xFAD mice (**c**). **d**, Heatmap showing gene set overlap analysis results comparing the sets of genes from the mouse ST co-expression modules with the co-expression modules identified from the Chen et al.[6] study. Fisher's exact test results shown as follows: Not significant (ns), p > 0.05; * p <= 0.05; ** p <= 0.01; *** p <= 0.001, **** p<= 0.0001.

### Bibliography


1. Samuel E. Marsh, Alec J. Walker, Tushar Kamath, Lasse Dissing-Olesen, Timothy R. Hammond, T. Yvanka de Soysa, Adam M. H. Young, Sarah Murphy, Abdulraouf Abdulraouf, Naeem Nadaf, et al. Dissection of artifactual and confounding glial signatures by single-cell sequencing of mouse and human brain. *Nature Neuroscience*, 25(3):306–316, 2022. ISSN 1097-6256. doi: 10.1038/s41593-022-01022-8.
2. Massimo Andreatta and Santiago J. Carmona. UCell: Robust and scalable single-cell gene signature scoring. *Computational and Structural Biotechnology Journal*, 19:3796–3798, 2021. ISSN 2001-0370. doi: 10.1016/j.csbj.2021.06.043.
3. Edward Zhao, Matthew R. Stone, Xing Ren, Jamie Guenthoer, Kimberly S. Smythe, Thomas Pulliam, Stephen R. Williams, Cedric R. Uytingco, Sarah E. B. Taylor, Paul Nghiem, et al. Spatial transcriptomics at subspot resolution with BayesSpace. *Nature Biotechnology*, pages 1–10, 2021. ISSN 1087-0156. doi: 10.1038/s41587-021-00935-2.
4. Marc Elosua-Bayes, Paula Nieto, Elisabetta Mereu, Ivo Gut, and Holger Heyn. SPOTlight: seeded NMF regression to deconvolute spatial transcriptomics spots with single-cell transcriptomes. *Nucleic Acids Research*, 49(9):e50–e50, 2021. ISSN 0305-1048. doi: 10.1093/nar/gkab043.
5. Suoqin Jin, Christian F. Guerrero-Juarez, Lihua Zhang, Ivan Chang, Raul Ramos, Chen-Hsiang Kuan, Peggy Myung, Maksim V. Plikus, and Qing Nie. Inference and analysis of cell-cell communication using CellChat. *Nature Communications*, 12(1):1088, 2021. doi: 10.1038/s41467-021-21246-9.
6. Wei-Ting Chen, Ashley Lu, Katleen Craessaerts, Benjamin Pavie, Carlo Sala Frigerio, Nikky Corthout, Xiaoyan Qian, Jana Laláková, Malte Kühnemund, Iryna Voytyuk, et al. Spatial Transcriptomics and In Situ Sequencing to Study Alzheimer's Disease. *Cell*, 182(4):976–991.e19, 2020. ISSN 0092-8674. doi: 10.1016/j.cell.2020.06.038.
